# A family of unusual immunoglobulin superfamily genes in an invertebrate histocompatibility complex

**DOI:** 10.1101/2022.03.04.482883

**Authors:** Aidan L. Huene, Steven M. Sanders, Zhiwei Ma, Anh-Dao Nguyen, Sergey Koren, Manuel H. Michaca, James C. Mullikin, Adam M. Phillippy, Christine E. Schnitzler, Andreas D. Baxevanis, Matthew L. Nicotra

**Affiliations:** Starzl Transplantation Institute, University of Pittsburgh, Pittsburgh, PA, USA; Center for Evolutionary Biology and Medicine, University of Pittsburgh, Pittsburgh, PA, USA; Computational and Statistical Genomics Branch, National Human Genome Research Institute, National Institutes of Health, Bethesda, MD, USA; NIH Intramural Sequencing Center, National Institutes of Health, Rockville, MD, USA; Whitney Laboratory for Marine Bioscience, University of Florida, St. Augustine, FL, USA; Department of Biology, University of Florida, Gainesville, FL USA; Department of Immunology, University of Pittsburgh, Pittsburgh, PA, USA

**Keywords:** allorecognition, *Hydractinia*, AlphaFold, gene complex, non-self recognition

## Abstract

Most colonial marine invertebrates are capable of allorecognition, the ability to distinguish between themselves and conspecifics. One long-standing question is whether invertebrate allorecognition genes are homologous to vertebrate histocompatibility genes. In the cnidarian *Hydractinia symbiolongicarpus*, allorecognition is controlled by at least two genes, *Allorecognition 1* (*Alr1*) and *Allorecognition 2* (*Alr2*), which encode highly polymorphic cell surface proteins that serve as markers of self. Here, we show that *Alr1* and *Alr2* are part of a family of 41 *Alr* genes, all of which reside a single genomic interval called the Allorecognition Complex (ARC). Using sensitive homology searches and highly accurate structural predictions, we demonstrate that the Alr proteins are members of the immunoglobulin superfamily (IgSF) with V-set and I-set Ig domains unlike any previously identified in animals. Specifically, their primary amino acid sequences lack many of the motifs considered diagnostic for V-set and I-set domains, yet they adopt secondary and tertiary structures nearly identical to canonical Ig domains. Thus, the V-set domain, which played a central role in the evolution of vertebrate adaptive immunity, was present in the last common ancestor of cnidarians and bilaterians. Unexpectedly, several Alr proteins also have immunoreceptor tyrosine-based activation motifs (ITAMs) and immunoreceptor tyrosine-based inhibitory motifs (ITIMs) in their cytoplasmic tails, suggesting they could participate in pathways homologous to those that regulate immunity in humans and flies. This work expands our definition of the IgSF with the addition of a family of unusual members, several of which play a role in invertebrate histocompatibility.

**Significance Statement:** The immunoglobulin superfamily (IgSF) is one of the largest and most functionally versatile domain families in animal genomes. Although their amino acid sequences can vary considerably, IgSF domains have been traditionally defined by conserved residues at several key positions in their fold. Here, we sequenced an invertebrate histocompatibility complex and discovered a family of IgSF genes with amino acid sequences that lack most of these residues yet are predicted to adopt folds virtually identical to canonical V-set and I-set IgSF domains. This work broadens the definition of the IgSF and shows that the V-set domain was present earlier in animal evolution than previously appreciated.

## Introduction

Allorecognition is the ability to distinguish self from non-self within the same species. Most encrusting colonial marine invertebrates, including sponges, corals, hydroids, bryozoans, and ascidians, are capable of allorecognition (1). This ability enables colonies to compete with conspecifics for space but prevents them from competing with themselves as they grow on three-dimensional surfaces (2). Allorecognition also reduces the risk of stem cell parasitism, which can occur if unrelated colonies fuse and one colony’s germline contributes disproportionately to the gametic output of the chimera (3).

Allorecognition has long attracted the attention of marine ecologists interested in spatial competition (2), population geneticists interested in the generation and maintenance of allelic diversity (4), and evolutionary biologists interested in units of selection and the origins of multicellularity (5, 6). In addition, ever since immunologists learned that corals and sea squirts exhibit allorecognition, they have wondered whether the genes that underlie this ability might be homologous to vertebrate histocompatibility genes (7). If so, studying invertebrate allorecognition could help resolve the evolutionary history of immunity and perhaps lead to novel therapies in immunity and transplantation. Together, these interests have motivated the study of allorecognition genes in several species, including the poriferan *Amphimedon queenslandica* (8), the protochordate *Botryllus schlosseri* (9–11), and the cnidarian *Hydractinia symbiolongicarpus* (12–15).

In *Hydractinia*, allorecognition is controlled by the allorecognition complex (ARC), which encodes two linked genes called *Allorecognition 1* (*Alr1*) and *Allorecognition 2* (*Alr2*) (12, 13). In laboratory strains, each gene has two alleles, and together they control allorecognition using a “missing-self” strategy. Colonies that share at least one allele at both genes recognize each other as self and fuse to create a larger colony (**Fig. 1A**). Colonies that do not share alleles at either locus recognize each other as non-self and fight by discharging harpoon-like organelles called nematocysts into their opponents (**Fig. 1B**). Colonies that share alleles only at one locus—either *Alr1* or *Alr2—*fuse but later separate.

**Fig. 1.**
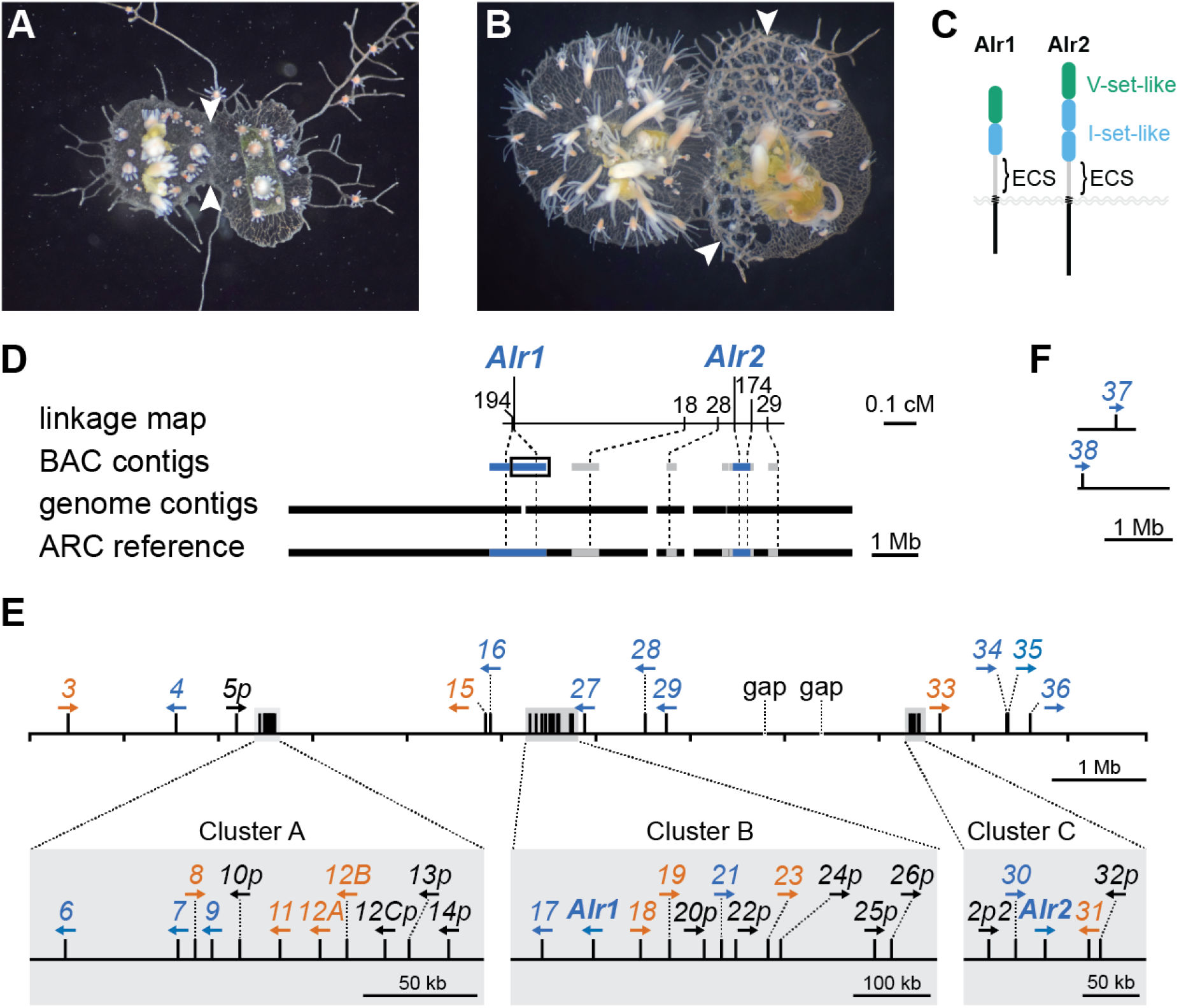
Allorecognition in *Hydractinia* and the assembly of the *Alr* gene complex. (A) Fusion between two compatible colonies. Arrows point to the region of fusion. (B) Rejection between two incompatible colonies. The colony on the left has grown over the colony on the right. Arrows point to specialized structures called hyperplastic stolons, which are destroying the underlying tissue. (C) Domain architecture of Alr1 and Alr2. (D) Chromosome walks from the ARC linkage map (top) generated six BAC contigs (below; blue = previously published; gray = unpublished). These were aligned to contigs from the assembled genome of an animal homozygous across the ARC (black). The resulting 11.83 Mb reference sequence was constructed by concatenating the BAC and genome assemblies (bottom). (E) Identity, location, and orientation of *Alr* family members within the ARC reference (blue, *bona fide* gene; orange, putative gene; black, pseudogene). (F) Two *Alr* genes located in genome contigs that could not be physically linked to the ARC reference sequence.

*Alr1* and *Alr2* encode transmembrane proteins with highly polymorphic extracellular domains (14, 15). In nature, there are tens to hundreds of alleles for each gene (15, 16). *In vitro* studies have shown that the Alr1 protein is capable of *trans* (cell-to-cell) homophilic binding, which only occurs between allelic variants with similar extracellular sequences (17). The same is true for Alr2 (17, 18). This variant-specific homophilic binding is hypothesized to be the mechanism of self/non-self discrimination *in vivo*.

The homology of *Alr1* and *Alr2* to other genes is unresolved. When they were originally identified, it was not possible to identify orthologs for either gene outside of *Hydractinia*. However, BLAST searches did return statistically significant alignments between domains in the Alr1/2 extracellular regions and immunoglobulin (Ig) domains (**Fig. 1C**). These hits had low sequence identity, making it difficult to determine whether the Alr domains belonged to the immunoglobulin superfamily (IgSF).

It is also unclear how many *Alr* genes exist in *Hydractinia. Alr1* is flanked by several *Alr1-*like sequences (15), and the genomic region immediately upstream *Alr2* contains two *Alr2* pseudogenes (14, 19). However, the full extent of this gene family is unknown because only a fraction of the ARC has been sequenced. Identifying additional *Alr* genes is of particular interest because the ARC most likely contains at least one additional allodeterminant. Evidence for this comes from the fact that *Alr1* and *Alr2* can fail to predict allorecognition responses in field-collected colonies (14, 15). The unidentified allodeterminant(s) likely reside in the ARC because genetic studies have shown that all *Hydractinia* allodeterminants, including dominant and codominant modifiers, are linked to *Alr1* and *Alr2* (20).

Here, we report the discovery of a family of 41 *Alr* loci, all encoded in the ARC. These genes show evidence of ancient and recent duplications, exon shuffling, and alternative splicing that could give rise to functionally distinct isoforms. A majority encode single-pass transmembrane proteins with V-set and I-set Ig domains with highly unusual amino acid sequences. Unexpectedly, most Alr proteins also encode a fibronectin III (Fn3)-like fold. Several Alr proteins also have immunoreceptor tyrosine-based activation motifs (ITAMs) or immunoreceptor tyrosine-based inhibitory motifs (ITIMs) in their cytoplasmic tails, suggesting a role for a conserved ITAM/ITIM-mediated signaling pathway in *Hydractinia*. Together, our results outline the full extent of a novel family of IgSF proteins, several of which are candidates for additional allodeterminants in *Hydractinia*.

## Results

### The ARC spans at least 11.8 Mb

In previous work, we generated a BAC library from a colony homozygous for the ARC-F haplotype (colony 833-8 in *SI Appendix* Fig. S1), and used it to perform chromosome walks starting from markers in the ARC linkage (**Fig. 1D**) (13–15). The minimum tiling path of each walk was sequenced, resulting in six BAC contigs with a total length of 2.9 Mb (**Fig. 1D** and *SI Appendix* Fig. S2A). Here, we sequenced and assembled the genome of colony 236-21, an ARC-F homozygote and a descendant of colony 833-8 (*SI Appendix* Fig. S1). Genomic DNA was sequenced via PacBio long-read sequencing and polished with Illumina data to create a genome assembly that was 431 Mb long, with 5,697 contigs and an N50 of 224 Kb. We then aligned the original BAC contigs to this new assembly using Nucmer (Kurtz et al. 2004). We identified five genome contigs that overlapped the BAC contigs with >99% sequence identity (**Fig. 1D** and *SI Appendix* Fig. S3, Table S1). The only major discrepancies between the BAC contigs and the genome contigs were in repeat regions. Therefore, we merged these sequences by filling the gaps between the BAC contigs with sequences from the genome assembly. The resulting ARC-F reference sequence spans 11.83 Mb and contains two gaps of unknown size (**Fig. 1D** and Dataset S1).

### The ARC contains a large family of *Alr* genes and pseudogenes

We next annotated all *Alr*-like genes, guided by *ab initio* gene predictions, sequence similarity to *Alr1* and *Alr2*, and RNAseq data. Sequences without homology to *Alr1* or *Alr2* were not annotated. As new gene models were created, we used them in iterative TBLASTX searches to identify *Alr* genes that might not have been detected in earlier similarity searches. Finally, to identify *Alr* genes that exist outside the ARC-F reference sequence, we used TBLASTX to query the full genome assembly with the amino acid translation of each *Alr* gene model. All gene models were then numbered sequentially, with pseudogenes receiving a lowercase ‘*p*’ at the end of their name (e.g., *Alr5p*). Alternative splice variants were indicated with a decimal number (e.g., *Alr1.2*). Gene models whose full-length predicted amino acid sequences had >80% sequence identity were given the same number followed by a letter (e.g., *Alr12A* and *Alr12B*).

In total, we created 41 gene models (**Fig. 1E**). All but two were in the ARC reference sequence (Dataset S2). More than half (27/41) were encoded in one of three *Alr* clusters, which we named A, B, and C (**Fig. 1E**). The remaining genes, *Alr37* and *Alr38*, were on contigs not contiguous with the reference sequence (**Fig. 1E** and Datasets S3-S6). The expression level of each gene model was estimated from our RNAseq data (**Fig. 2A**). Gene models with less than 1 FKPM were deemed unexpressed. To aid our analysis, we then classified each gene model as a *bona fide* gene, putative gene, or pseudogene.

**Fig. 2.**
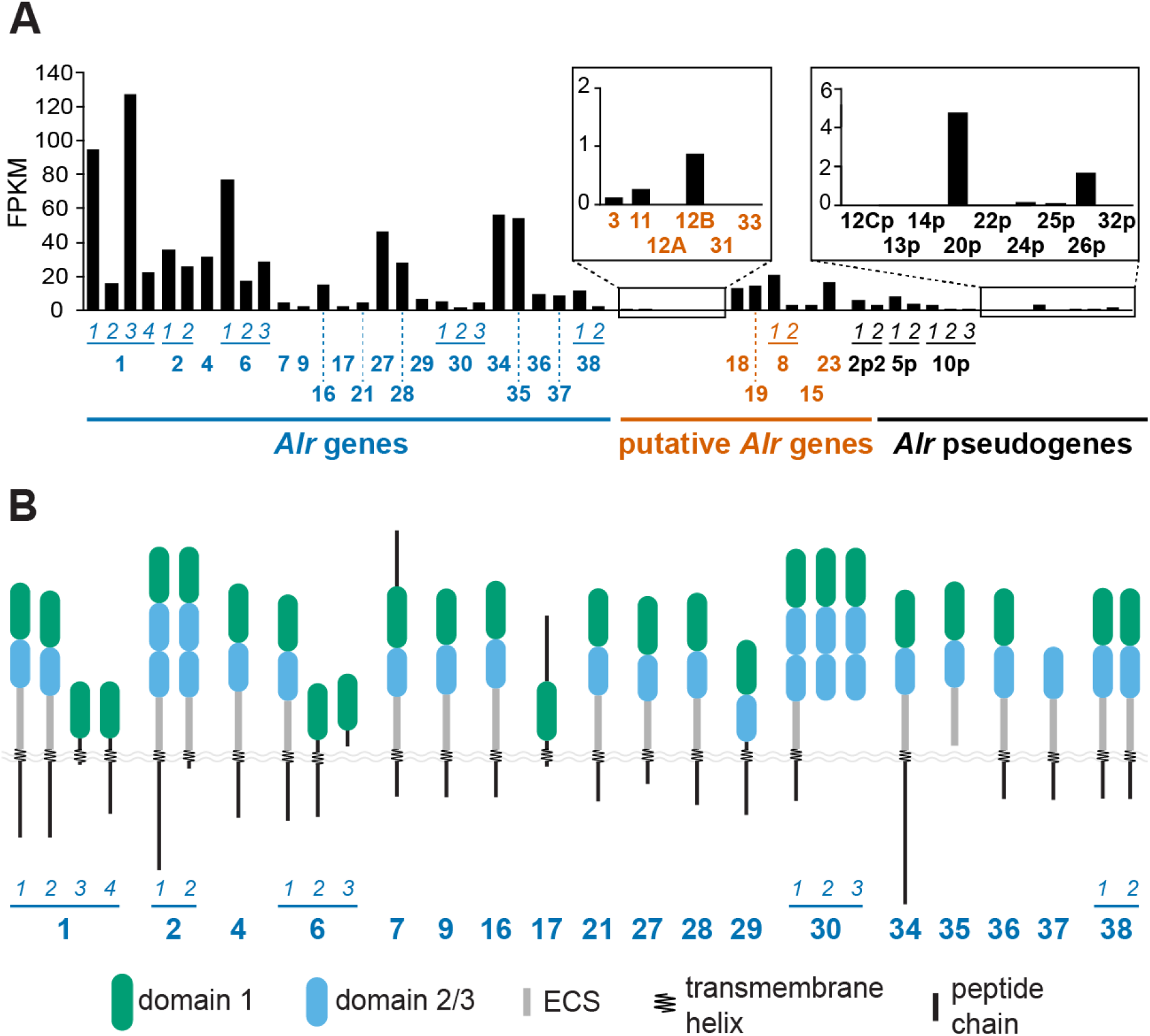
Expression, domain architecture, and alternative splicing of *Alr* genes. (A) Estimated expression level of each *bona fide* gene, putative gene, and pseudogene. FPKM, fragments per kilobase mapped. Genes are identified by bold numbers. Splice variants are indicated by horizontal lines and numbers in italics. The order of the putative genes follows their appearance in panels C-E. (B) Domain architecture of *Alr* genes.

A gene model was classified as a *bona fide* gene if it had one open reading frame (ORF) and each exon was expressed and properly spliced. Eighteen models fit this definition, including *Alr1* and *Alr2* (Datasets S7 and S8). As shown in **Fig. 2B**, *Alr* genes generally encoded single-pass transmembrane proteins with 1-3 domains similar to the *Alr1* and *Alr2* Ig-like domains, an extracellular spacer (ECS), a transmembrane helix, and a cytoplasmic tail. Without exception, each Ig-like domain, ECS, and transmembrane helix was encoded by a single exon.

A gene model was classified as a putative gene if it had features that made us hesitant to call it a *bona fide* gene, but was not obviously a pseudogene. Eleven gene models fit this definition (Datasets S9 and S10). Six had clear sequence similarity to *bona fide Alr* genes but were unexpressed (**Fig. 2A** and *SI Appendix* Fig. S3A). We did not call them pseudogenes because they could be expressed at developmental time points or in tissues not represented in our RNAseq dataset. Two gene models had ORFs that would encode a full Alr protein, but there was no evidence of splicing between exons 1, 2, and 3 (*SI Appendix* Fig. S3B). Three gene models did not encode a signal peptide (*SI Appendix*, Fig. S3C).

A gene model was classified as a pseudogene if it was similar to a *bona fide* or putative *Alr* gene but was truncated by nonsense or frameshift mutations. Twelve gene models fit this definition (*SI Appendix*, Table S2). Several pseudogenes were expressed at modest levels relative to other *Alr* genes (**Fig. 2A**).

We paid particular attention to the region directly upstream of *Alr2* because it contained two pseudogenes reported in previous publications (14, 19). The first, immediately upstream of *Alr2*, was named CDS6P by Nicotra *et al*. (2009) and *alr2P1* by Rosengarten *et al*. (2011). It was assumed to be a non-functional partial duplication of *Alr2*. Here, we identified additional exons encoding a transmembrane domain, cytoplasmic tail, and 3’ UTR. We therefore classified this locus as a *bona fide* gene and named it *Alr30*. The second pseudogene was called CDS5P by Nicotra *et al*. (2009) and *alr2P2* by Rosengarten *et al*. (2011). Here, we also concluded the locus was a pseudogene. For consistency with the previous work, we have named this pseudogene *Alr2p2*.

### Alternative splicing alters the domain architecture of several *Alr* gene products

Several *Alr* genes were alternatively spliced in ways that would change their gene product’s domain architecture. *Alr1*, for example, had four splice variants. *Alr1.1* and *Alr1.2* were previously described (15), but in *Alr1.3* and *Alr1.4*, exon 2 was spliced to new exons encoding alternative transmembrane domains and cytoplasmic tails (*SI Appendix*, Fig. S4A). This resulted in two isoforms lacking domain 2 and the ECS (**Fig. 2B**). *Alr6* was also alternatively spliced in a similar manner (*SI Appendix*, Fig. S4B). Notably, in *Alr6.3*, exon 2 was spliced to exon 11 and lacked a transmembrane helix, raising the possibility that its gene product is secreted (**Fig 2B**). A similar splicing pattern, potentially leading to secreted gene products, was observed in Alr30 and Alr35 (Fig. 2B and *SI Appendix* Fig. S4C). At *Alr2*, transcripts lacking the 22-bp exon 7 were also detected, which would introduce a frameshift that truncated the cytoplasmic tail (**Fig. 2B** and *SI Appendix* Fig. S4D).

### Sequences of Alr family members are highly diverse and undergo exon shuffling

The shared domain architecture of *Alr* proteins suggested a history of gene duplication. To investigate the evolutionary relationships between *Alr* family members, we attempted to create a single multiple sequence alignment for the gene products of all *Alr* genes and putative genes. However, their sequences were so divergent that it was impossible to obtain a high-quality alignment even after restricting ourselves to sequences of similar length. This drove us to assess overall sequence similarity within the family by performing all possible pairwise alignments. We found the average percent identity between any two amino acid sequences (excluding splice isoforms of the same gene) was 24.3% ± 8.6% (*SI Appendix*, Fig. S5). Only 2% of pairwise alignments had more than 50% identity. Thus, a substantial amount of sequence evolution has occurred since the origin of the *Alr* family.

Closer analysis of the pairwise alignments suggested a history of exon shuffling between *Alr* genes. Specifically, we found several cases in which two predicted proteins were very similar in some domains but very divergent in others (**Fig. 3A**). For example, Alr12A and Alr12B were >90% identical along their entire length except for domain 1, which was only 64% identical (**Fig. 3A**). This pattern of domain-level variation is consistent with previous studies indicating a history of exon shuffling between *Alr1* and nearby *Alr*-like sequences (15) and between *Alr2* and nearby *Alr2* pseudogenes (16). Our data indicate exon shuffling could occur between several other *Alr* genes.

**Fig. 3.**
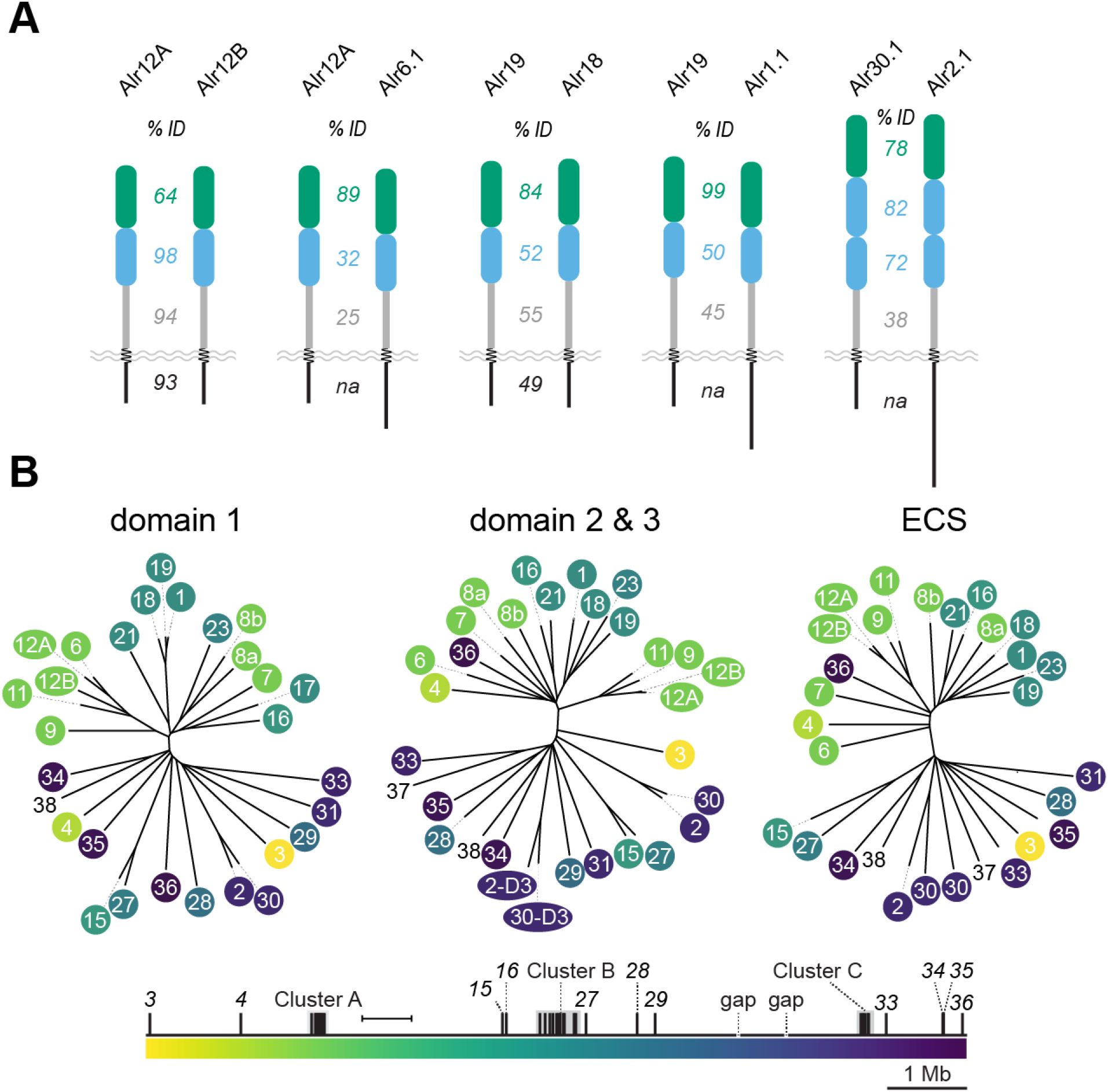
Sequence similarity between Alr extracellular domains. (A) Evidence of exon shuffling between *Alr* genes. Numbers between protein pairs indicate sequence identity within that domain/region. (B) Neighbor-joining trees of Alr extracellular domains. Leaves of each tree are color coded according to their genomic position. Domains from *Alr37* and *Alr38* are not color coded. Branch lengths calculated according to the BLOSUM26 matrix. Scale bar = 100 units.

This pattern of exon shuffling led us to compare Alr proteins at the domain level. We subdivided each sequence into its constitutive extracellular domains, then produced multiple sequence alignments (Datasets S11-S13) and neighbor-joining trees (**Fig. 3B**) for each domain type. This revealed a pattern in which domains were more similar if they were encoded close to each other in the genome (**Fig. 3B**). While this analysis has limited power to elucidate the history of the *Alr* gene family, it does suggest that the duplications within Cluster C occurred after it split from Clusters A/B.

### The cytoplasmic tails of many Alr proteins contain ITAMs or ITIMs

Unlike their extracellular domains, the Alr cytoplasmic tails were too diverse to be included in a single alignment. Therefore, we used CD-HIT to cluster them at 20% sequence identity. This placed 16/32 into three groups, for which we created separate alignments (*SI Appendix*, Fig. S6). Of the remaining 16 tails, three were <14 aa and the rest could not be grouped with other sequences. Thus, the cytoplasmic tails of Alr proteins are more divergent than their extracellular domains.

The domain architecture of most Alr proteins suggested they might be receptors with intracellular signaling functions. To investigate this, we searched their cytoplasmic tails for signaling motifs. We found immunoreceptor tyrosine-based activation motifs (ITAMs) in the tails of six *bona fide* and eight putative Alr proteins (**Fig. 4A** and SI Appendix Fig. S7). ITAMs, which have a consensus sequence of Yxx[I/L]x_(6-9)_Yxx[L/I] (21), are found in receptors that activate immune responses in vertebrates (22) and phagocytosis of damaged cells in *Drosophila* (23).

**Fig. 4.**
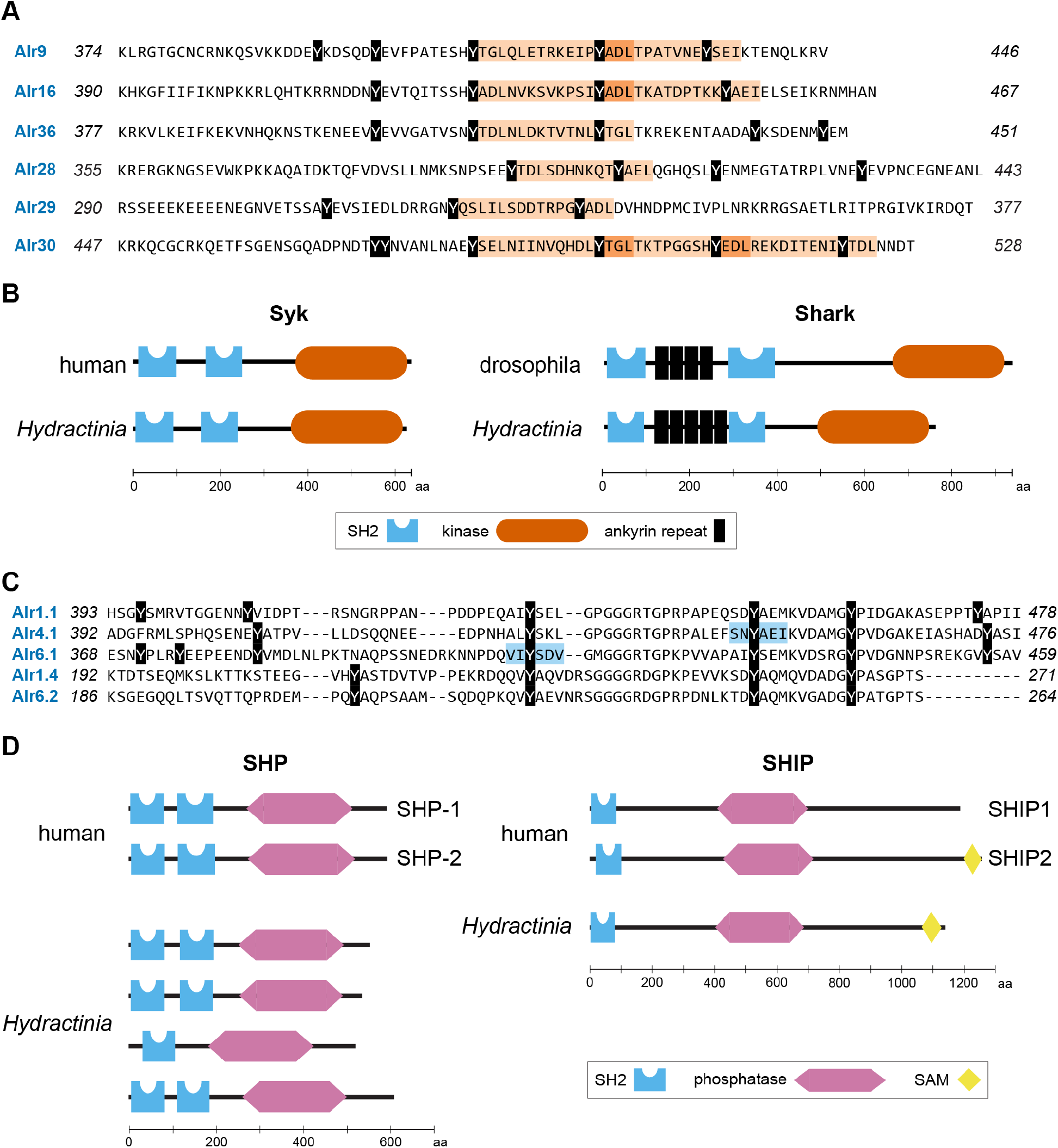
Alr ITAM and ITIM motifs and presence of select signaling molecules. (A) Cytoplasmic tails of Alr proteins with ITAMs (orange background). Overlapping ITAMs are indicated with heavier shading. (B) Comparison of human Syk and *Drosophila* Shark to *Hydractinia* Syk and Shark. (C) Truncated alignment of cytoplasmic tails with ITIMs (blue background). All tyrosines have a black background. (D) Comparison of human SHP and SHIP proteins to *Hydractinia* SHP and SHIP homologs.

Phosphorylated ITAMs are bound by the dual SH2 domains of a kinase called *Syk* in vertebrates and *Shark* in insects (24). Syk and Shark are related proteins and differ in that Shark has a set of ankyrin repeats between its two SH2 domains. To determine whether the *Hydractinia* genome encodes *Syk* or *Shark-*like kinases that might bind to these ITAMs, we performed a TBLASTN search of the complete genome assembly with the amino acid sequences of human *Syk* and *Drosophila Shark*, identifying *Hydractinia* homologs of each (**Fig. 4B**). This is consistent with previous work that has identified *Syk*-like and *Shark-*like kinases in *Hydra* (25, 26).

A second motif in vertebrates called the immunoreceptor tyrosine-based inhibitory motif (ITIM) is found in receptors that counteract ITAM-mediated signaling and downregulate immune responses (27). We found ITIMs, defined as [I/L/V/S]xYxx[I/V/L] (28), in two Alr tails, both from group 1 (**Fig. 4C**). In mammals, phosphorylated ITIMs are bound by the SH2 domains of SHP-1 and SHP-2, two phosphatases that dephosphorylate ITAM-bearing receptors, Syk-like proteins, and other components of activating pathways (29, 30). ITIMs are also bound by the phosphoinositide phosphatases SHIP1 and SHIP2, which dampen immune cell activation via the PI3K pathway (31, 32). We searched the *Hydractinia* genome and identified four homologs of SHP-1/2 and one SHIP1/2 homolog (**Fig. 4D**).

Together, these data show that many *Alr* genes have ITAMs and ITIMs, motifs that regulate the recognition of self and non-self in other animals. Moreover, the *Hydractinia* genome includes homologs of the enzymes that bind phosphorylated ITAMs or ITIMs and act as effectors of cellular activation or inhibition.

### Domain 1 is a V-set Ig domain

Domain 1 of Alr1 and Alr2 was originally described as V-set-like (14, 15). To determine whether domain 1 was similar to V-set Ig domains in Alr3-Alr38, we first used HMMER to compare each sequence to Pfam, a database of hidden Markov models for protein families (33). At an E-value cutoff of <0.01, only 6/29 sequences were similar to V-set domains (*SI Appendix*, Table S3). We also used HHpred, a method that is more sensitive to remote homologies (34) to search the Structural Classification of Proteins extended (SCOPe) database, which classifies protein domains according to structural and evolutionary relationships (35). Using this approach, we found that 19/29 sequences had a >95% probability of homology to V-set domains (*SI Appendix*, Table S3).

Homologous proteins can evolve such that their primary sequences become highly divergent but their structures remain relatively unchanged (36). Therefore, we predicted the tertiary structure of each domain 1 with Colabfold (37) to further investigate its homology (Dataset S14). Colabfold is a Google Colaboratory implementation of AlphaFold2 (38), which is capable of producing structural predictions with sub-angstrom root mean square deviation from experimental structures (39). Each residue in a model produced by Colabfold is assigned a predicted local distance difference test (pLDDT) score, which estimates how well the prediction would agree with an experimental structure. Residues with pLDDT > 90 are considered highly accurate and have their side chains oriented correctly 80% of the time (38, 39). Residues with pLDDT > 70 generally have their backbones predicted correctly. For the Alr domain 1 sequences, Colabfold produced the structural predictions with average (model-wide) pLDDT scores ranging from 80.6 to 97.4 (**Fig. 5A** and *SI Appendix*, Table S4).

**Fig. 5.**
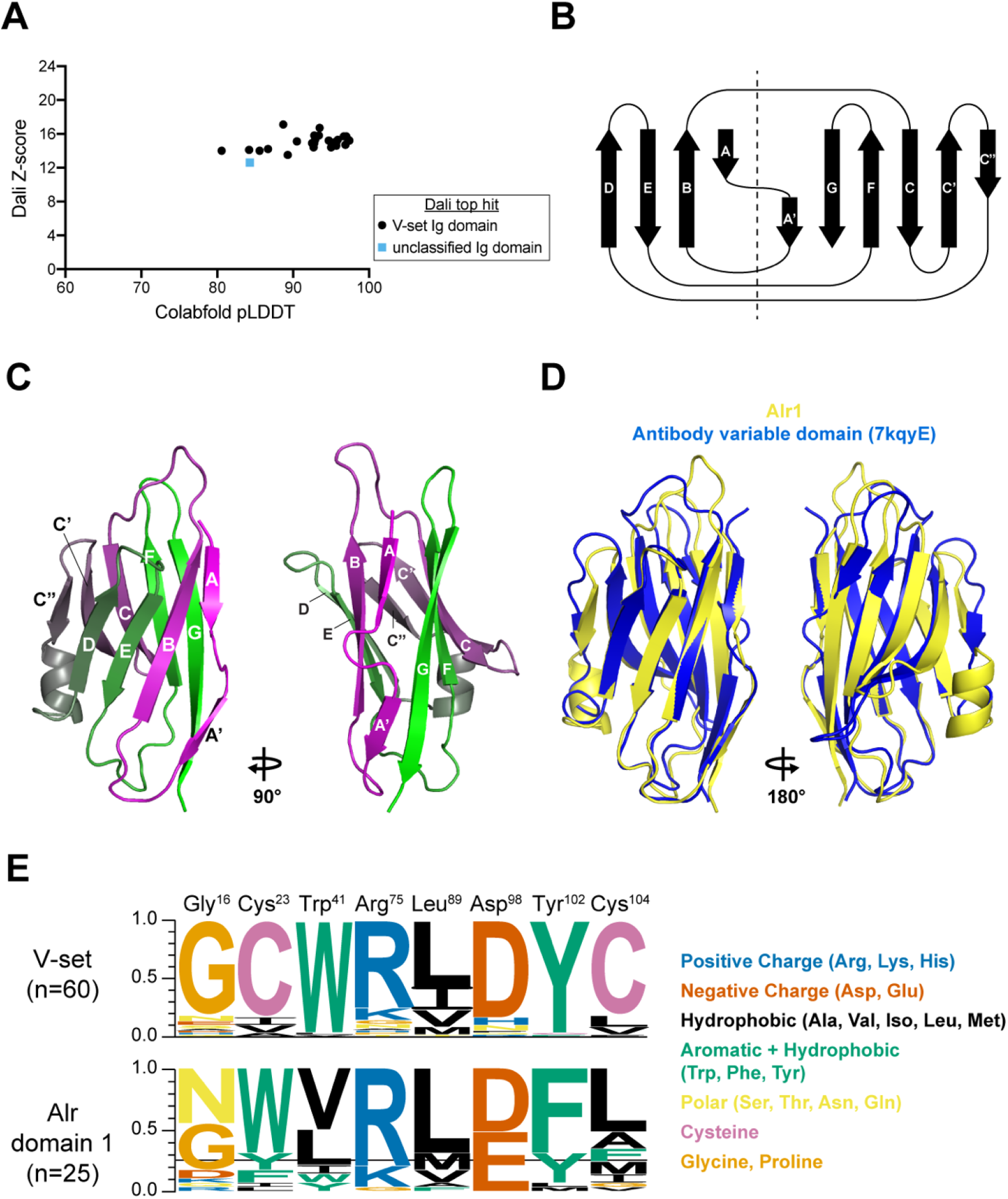
Sequence analysis and structural predictions of domain 1. (A) Plot of each domain 1 model’s average pLDDT score versus its alignment Z-score to the top PDB model identified by DALI. (B) Structural alignment of Alr1 domain 1 to a V-set Ig domain from human heavy-chain antibody (PDB 7KQY). (C) Topology of β-strands in V-set folds. Dotted line separates the strands within the same β-sheet. (D) V-set β-strands labeled on the predicted structure of Alr1 domain 1. (E) Sequence logo comparing frequency of amino acids at eight conserved positions in V-set Ig domains (top) and Alr domain 1 sequences (bottom).

Next, we performed structural alignments against the Protein Data Bank (rcsb.org; 40) with Dali (41) and PDBeFOLD (42). The top hits from both methods were to V-set Ig domains (e.g., Fig. 5B) (Table S4). Structural alignments produced by Dali are assigned a Z-score, which is used to estimate the likelihood that the two proteins are homologous. Z-scores between 8-20 indicate likely homologs (41). The domain 1 alignments had Z-scores ranging from 12.6-16.7, indicating probable homology with V-set domains (**Fig. 5A**), even though their overall sequence identities were 9-20% (*SI Appendix*, Table S4). Thus, our analysis of the primary, secondary, and tertiary structures of domain 1 all indicated they belong to the V-set family of Ig domains.

V-set domains have nine β-strands named A, B, C, C’, C”, D, E, F, and G according to their position in the primary amino acid sequence. Strand C” is only found in V-set domains. Strand A is split into A and A’. These β-strands are arranged as a Greek key to form a β-sandwich, with one β-sheet consisting of strands A, B, E, and D, and the other consisting of strands A’, G, F, C, C’, and C” (**Fig. 5C**). This is often referred to as the V-frame (43).

To determine whether domain 1 has the V-frame, we used STRIDE (44) to determine the secondary structure of our models, then assigned letters to the β-strands (*SI Appendix*, Fig. S8). Twenty five models had all nine β-strands (e.g., **Fig. 5D**). Four models were with the missing strand being in either the A or A’ position (*SI Appendix*, Fig. S8). Notably, all models had the V-set specific C” strand (*SI Appendix*, Fig. S8). In addition, 17/29 models included a 7 aa α-helix between the C” and D strands, which is not typically found in V-set domains.

These results led us to question why HMMER did not identify many Alr domains as V-set domains. Previous studies of V-set domains have identified a set of eight residues that are highly conserved, even across domains with as little as 20% sequence identity (43, 45).

According to the nomenclature of Cannon et al (45), they are Gly^16^, Cys^23^, Trp^41^, Arg^75^, Leu^89^ (or other hydrophobic), Asp^98^, Tyr^102^, and Cys^104^. To determine whether these residues are conserved in domain 1, we generated a multiple sequence alignment between them and 60 canonical V-set sequences from the Pfam V-set sequence profile (pf07686). We then identified the residues that corresponded to the eight V-set residues (*SI Appendix*, Fig. S9). Our findings are summarized in **Fig. 5F**.

In V-set domains, Cys^23^, Trp^41^, and Cys^104^ form a nearly invariant structural motif called the ‘pin’ (46). The cysteines form a disulfide bridge between β-strands B and F, while the tryptophan packs against the bond to stabilize the hydrophobic core of the β-sandwich. All Alr domain 1 sequences, however, lacked these Cys residues, and only two had the Trp. Instead, Cys^25^ was replaced by bulky, aromatic amino acids (Trp, Phe, or Tyr). Cys^104^ and Trp^41^ were replaced by hydrophobic amino acids. Thus, in domain 1, the ‘pin’ is replaced by a set of bulky hydrophobic residues that might serve a similar function by stabilizing the core of the β-sandwich.

The fourth and fifth V-set residues, Arg^75^ and Asp^98^, form a salt bridge between the CD and EF loops. The salt bridge is thought to stabilize the “bottom” of the domain and is found only in the V-set and I-set immunoglobulin domains (43, 45). We found the salt bridge in all but three (26/29), although the negatively charged Asp was often replaced with similarly-charged Glu. Thus, the salt bridge, a hallmark of V-set and I-set domains, is also present in domain 1.

The sixth canonical residue, Tyr^102^, forms the ‘tyrosine corner’, a structural motif located at the start of the F strand and found only in Greek key proteins (47). While Tyr^102^ is highly conserved in V-set Ig-like domains (97% in our seed alignment), it was found in only 7/29 Alr domains (**Fig. 5E**). Instead, 20/29 had Phe, with its aromatic ring occupying the same location as that of Tyr^102^. Mutational studies have shown that a Tyr❼Phe mutation has no effect on the ability of V-set Ig domains to fold properly (48). Thus, the residues at position 102 are consistent with domain 1 folding like a V-set domain. 277C

The seventh canonical residue is Gly^16^, which is part of a β-turn between strands A’ and B. A β-turn is a series of four residues that reverses 180° on itself such that the distance between C^α^(*i*) and C^α^(*i*+3) is less than 7 Å (49). β-turns often feature a hydrogen bond between CO(*i*) and NH(*i*+3) — the carboxyl and amine groups on the first and fourth residues, respectively — but this is not a requirement (50). We found this β-turn in all Alr domain 1 models (*SI Appendix*, Fig. S10). Twenty-eight featured a hydrogen bond, defined by the criterion that O(*i*) is< 3.5 Å from N(*i*+3) (50). However, position *i + 2* was Gly in only eleven sequences. Thus, like V-set domains, domain 1 is predicted to have a β-turn between strands A’ and B, but in most cases it does not involve a glycine.

The eighth canonical V-set residue is a hydrophobic amino acid, typically leucine, at position 89. This residue resides at the center of the hydrophobic core. In Alr domain 1, 21/29 sequences had a leucine residue at this position. The remaining six had other hydrophobic residues. Thus, this canonical residue is shared between V-set and most domain 1 sequences.

In summary, the Alr V-set domains share some, but not all sequence motifs commonly considered diagnostic of the V-set family. In the case of the pin motif, tyrosine corner, and β-turn, the Alr sequences differ in a way that likely preserves the structural motif. Thus domain 1 appears to be a V-set Ig domain with a novel sequence profile.

### Domains 2 and 3 are I-set Ig domains

We next explored the relationship between Ig domains and the 29 domain 2 and 3 sequences encoded by *bona fide* and putative *Alr* genes. At an E-value cutoff of <0.01, HMMER identified 14 as I-set Ig-like domains (pf07679) and another four as Ig-like domains (pf13927) (*SI Appendix*, Table S5). HHpred indicated 23/32 had a >95% probability of being homologous to the I-set family (*SI Appendix*, Table S5).

To further explore their similarity to I-set Ig domains, we generated structural predictions for each domain, with average pLDDT scores ranging from 81.1 to 95.1 (**Fig. 6A** and *SI Appendix* Table S6 and Dataset S14). I-set domains are similar to V-set domains except that they lack a C” strand, and the C’ strand is often shorter (**Fig. 6B**). We found that 21/31 domain 2 and 3 models had an I-set topology (e.g., **Fig. 6C** and *SI Appendix* Fig. S11), and three more were only missing one of the two split A strands. Six others were missing the C’ strand (e.g., **Fig. 6D** and *SI Appendix* Fig. S11). These six models were I-set-like in having a split A strand but were similar to C1-set Ig domains in having a D strand and lacking a C’ strand (**Fig. 6B**, *SI Appendix* Fig. S11). The remaining domain, from Alr9, was predicted to have only 6 β-strands and lacked both C’ and D strands. When we searched for structural homologs, the top hits from PDBeFOLD and Dali were to I-set domains, although some hits were annotated as both I-set and C2-set domains, and the top hits Alr15 were to a filamin repeat (**Fig. 6A**, *SI Appendix* Table S6). Filamin repeats are in the “Early” (E-set) superfamily of immunoglobulin-like β-folds and are possibly related to the IgSF or fibronectin type III superfamilies (35, 51).

**Fig. 6.**
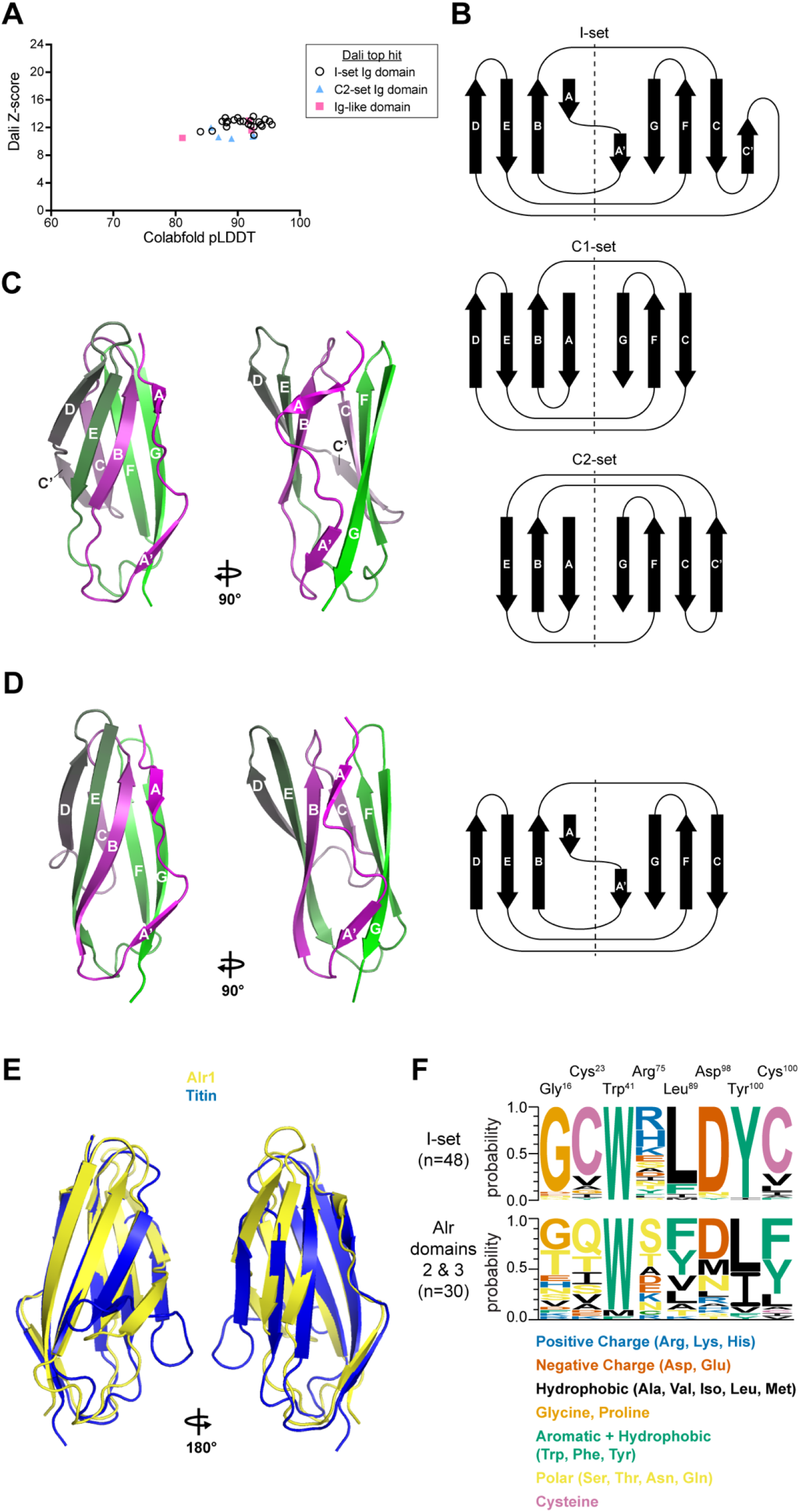
Sequence analysis and structural predictions of domains 2 and 3. (A) Plot of each domain 2 and domain 3 model’s average pLDDT score versus its alignment Z-score to the top PDB model identified by DALI. (B) Topology of β-strands in I-set, C1-set, and C2-set folds. Strand D is part of the EBA sheet in C1 Ig-domains, but is part of the CFG sheet in Fn3 and C2-set domains, where it is often labeled C’. Dotted line separates the strands within the same β-sheet. (C) I-set β-strands labeled on the predicted structure of Alr1 domain 2. (D) β-strands labeled on the predicted structure of Alr2 domain 2. The corresponding Greek key is shown to the right. (E) Structural alignment of Alr1 domain 2 to an I-set Ig domain from titin (PDB 2RJM). (F) Sequence logo comparing frequency of amino acids at conserved positions in I-set Ig domains (top) and Alr domains 2 and 3 (bottom).

We next investigated whether domains 2 and 3 had any of the conserved sequence motifs found in I-set domains. To do so, we aligned the Alr domains to 48 canonical I-set domains from the Pfam I-set sequence profile (pf07679) (*SI Appendix* Fig. S12). We then searched for the sequence motifs common to V-frame Ig-like domains (43, 45). These results are summarized as a sequence logo in Fig. 6F. With respect to the pin motif (C-W-C), all Alr domain 2 sequences had the central tryptophan, but the two domain 3 sequences had methionine in its place. All domains lacked the paired cysteines. One was replaced by a hydrophobic residue, and the second was replaced by residues bearing no consistent physicochemical property (**Fig. 6F** and *SI Appendix* Fig. S12). The Alr domains also lacked the salt bridge and tyrosine corner (**Fig. 6F**). The β turn between β-strands A’ and B was present in all but three structural models (*SI Appendix* Fig. S13). The last of the eight conserved residues, a hydrophobic residue (typically leucine) was present, although in many it was a tyrosine (**Fig. 6F**).

More recently, Wang (52) defined the sequence signature of I-set domains via nine sequence motifs, denoted *i* through *ix*. The first four motifs include the C-W-C pin (in motifs *i, ii*, and *iv*), the tyrosine corner (part of motif *i*), and the tight turn in the A’B loop (motif *iii*) (*SI Appendix* Fig. S12). We searched for the remaining five motifs and found that they were present in a majority of Alr domain 2 and 3 sequences (*SI Appendix* Fig. S12 and Fig. S14).

Taken together, our data indicate most domain 2 and 3 sequences belong to the I-set family of Ig domains. Although these domains lack the disulfide bridge, salt bridge, and tyrosine corner, they have most other sequence motifs associated with I-set domains, including the conserved tryptophan. Most are predicted to have an I-set fold, although several appear to have lost the C’ strand.

### Part of the ECS adopts a fibronectin type III-like fold

When Alr1 and Alr2 were described previously, no domains were identified in the ECS (14, 15). Here, we expanded our analysis to include all 29 ECS regions encoded by *bona fide* and putative *Alr* genes. Although HMMER searches against Pfam only return two hits to domains of unknown function, HHpred indicated part of the ECS had a 60-88% probability of homology to fibronectin type III (Fn3) domains (*SI Appendix*, Table S7). To help us define this potential domain, we aligned the ECS sequences to the 98 Fn3 sequences in the seed alignment of the Pfam Fn3 profile (pf00041.23). We found that the N-terminal portion of the ECS aligned reasonably well to other Fn3 domains, but the C-terminal portion did not (*SI Appendix*, Fig. S15).

Because structure predictions are often better for single domains, we removed the C-terminal portion of the ECS sequences (*SI Appendix* Fig. S15, Dataset S15) then predicted their structures with Colabfold. Twenty-six models had average pLDDT >90, with the remaining three models >80 (**Fig. 7A**, *SI Appendix* Table S8, and Dataset S14). Next, we investigated whether the Alr domains had a similar topology to Fn3 domains. Fn3 domains are an Ig-like fold with seven β-strands arranged in the same topology as C2-set immunoglobulin domains (53, 54) (**Fig. 7B**). All but one ECS model had seven β-strands (e.g., **Fig. 7C**), with the remaining model missing strand G (*SI Appendix* Fig. S16). In all models, the β-strands adopted a C2-set/Fn3 topology. We next searched the PDB for proteins with similar structures. All hits from Dali and PDBeFOLD were to Fn3 domains (**Fig. 7A** and *SI Appendix* Table S8). Thus, the secondary and tertiary structures of this Ig-like fold are predicted to be most similar to Fn3 domains.

**Fig. 7.**
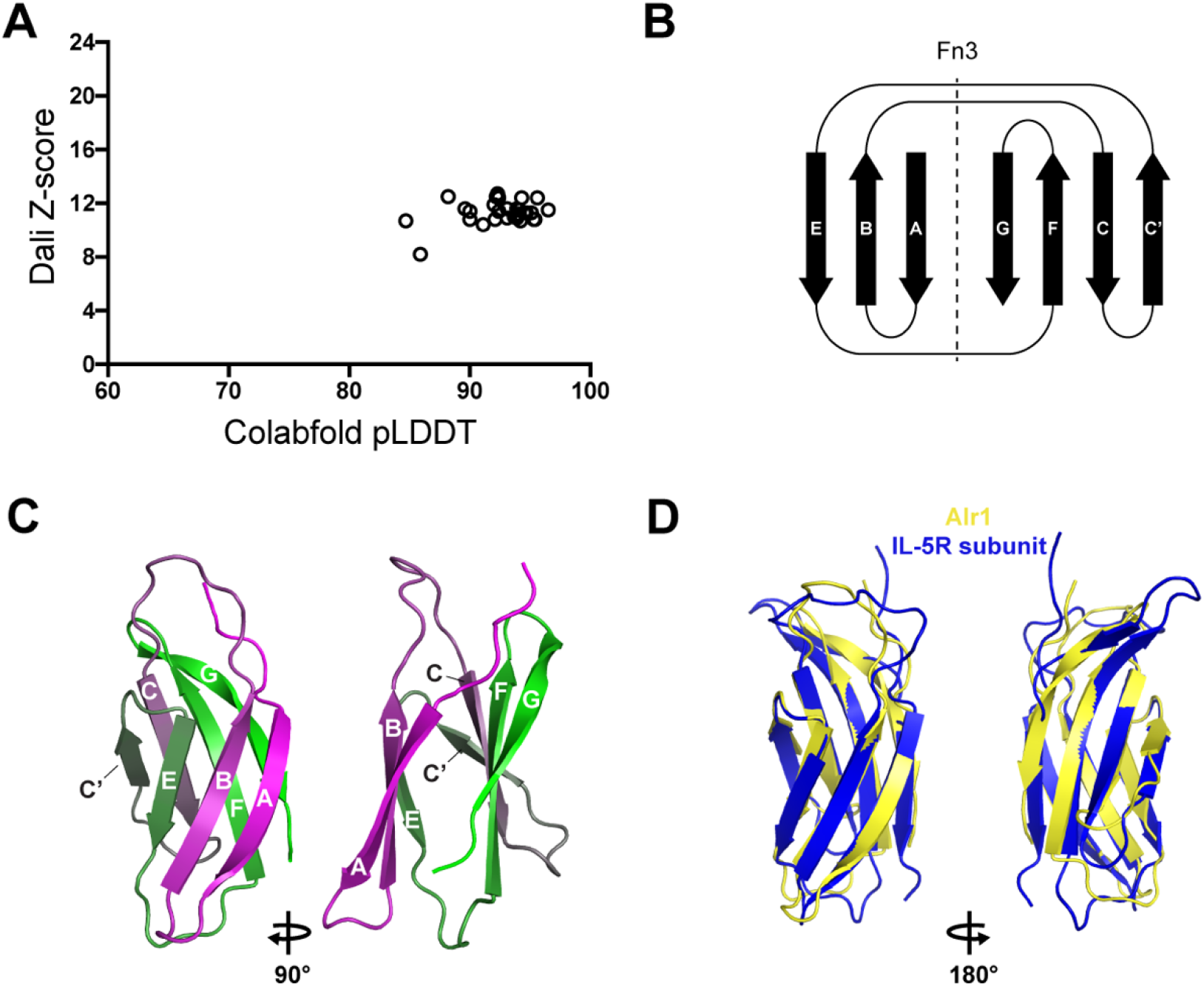
Sequence analysis and structural predictions of the ECS. (A) Plot of each ECS model’s average pLDDT score and corresponding alignment Z-score to the top PDB model identified by DALI. (B) Topology of β-strands in Fn3 folds. Dotted line separates the strands within the same β-sheet, (C) Predicted topology of β-strands in the Alr1 ECS folds. (D) Structural alignment of Alr1 ECS to a Fn3 domain from Interleukin-5 receptor subunit alpha (PDB 6H41).

The primary amino acid sequences of Fn3 domains have six conserved amino acids (53, 54). To determine whether the Alr Fn3-like domain had these residues, we aligned them to sequences from the Pfam Fn3 profile (pf00041.23). The ECS sequences only aligned well to Fn3 domains at their N-terminus (*SI Appendix* Fig. S17). With respect to the six conserved amino acids, ECS and Fn3 sequences shared a proline at the beginning of strand A, a tryptophan at the end of strand B, and a tyrosine at the beginning of strand C (*SI Appendix* Fig. S17 and S18). The fourth conserved residue in Fn3 domains, a tyrosine at the end of strand C, was present in 8/29 ECS sequences, and replaced by phenylalanine in 14/29 ECS sequences. However, unlike Fn3 domains, the ECS sequences were missing the leucine in the EF loop and the tyrosine residue that forms the tyrosine corner in strand F. Fn3 domains also have six additional ‘topohydrophobic’ positions (i.e., positions usually occupied by VILFMWY residues) (54), which were also present more than 50% of the time in the ECS sequences. (*SI Appendix* Fig. S17 and S18).

Taken together, these data indicate that most Alr proteins have a Fn3-like fold between their I-set domain and transmembrane helix.

### Alr proteins have six invariant cysteines likely to form disulfide bridges

While investigating protein alignments of the Alr domains, we identified six cysteine residues that were conserved across all sequences. These residues are not typically found in immunoglobulin or Fn3 folds. Three were in the Ig domain immediately preceding the Fn3-like domain, and three were located within the Fn3-like domain itself (**Fig. 8A,B**). Within the Ig domain, the first two cysteines were in β-strands A and B and were predicted to form a disulfide-bridge in 28/31 models. Within the Fn3-like domain, the first and third cysteines, located in β-strands B and E, were predicted to form a disulfide bridge in 28/31 models. Thus, an intra-domain disulfide bridge appears to stabilize the fold of most Alr I-set and Fn3-like folds.

**Fig. 8.**
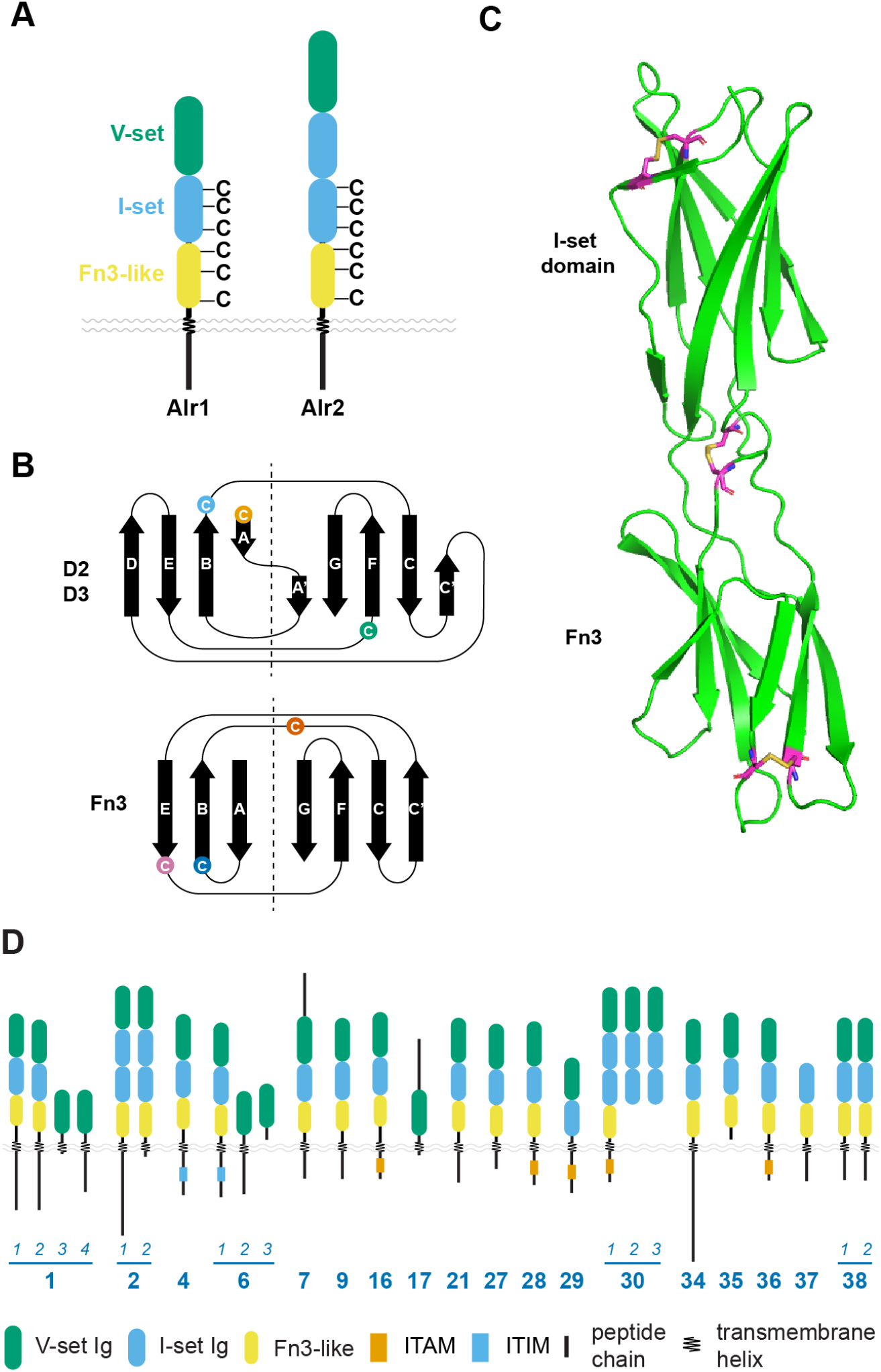
Invariant cysteines in Alr proteins and final domain predictions. (A) Occurrence of invariant cysteine residues found in Alr proteins with an Alr1-like or Alr2-like domain architecture. (B) Position of invariant cysteines in domains 2, 3 (I-set), and the ECS fold (Fn3-like). (C) Structural prediction of tandem I-set and Fn3-like domains of Alr1 shows each invariant cysteine forms a disulfide bond. (D) Domain architecture and motifs of Alr proteins

The two remaining invariant cysteines were in the EF loop of the Ig domain and the BC loop of the Fn3-like domain. These domains appear in tandem in 28 of the Alr proteins, raising the possibility that they are stabilized by a disulfide-bridge. To test this, we predicted structures for the tandem I-set Ig and Fn3-like domains from each Alr protein. All were predicted with high confidence (*SI Appendix*, Table S9), and in all models the two domains were predicted to be linked by a disulfide bridge (e.g., **Fig. 8C**).

### Alr orthologs may exist in other hydrozoans

To search for orthologs of the *Alr* gene family, we first queried *Hydra vulgaris* and *Clytia haemispherica*, the only hydrozoan species with publicly available proteomes. BLASTP searches against the predicted proteomes of each species returned hits with significant E-values, many of which were predicted to be single-pass transmembrane proteins (*SI Appendix* Table S10). Only two hits in *Clytia* were identified as having Ig domains by HMMER/Pfam. To broaden our search, we next queried the non-redundant protein database at NCBI via BLASTP, and excluded hydrozoans from the returned hits. For each Alr protein, the top scoring alignment was between its Ig domains and one or more Ig domains in the hit. No full-length hits were found. Notably, proteins from the anthozoans *Nematostella* and *Acropora* were not among the top hits. Further examination of cnidarian genomes will be required to determine the phylogenetic distribution of the *Alr* gene family.

## Discussion

We have shown that the *Hydractinia* ARC contains a family of genes homologous to *Alr1* and *Alr2*, which we have named the *Alr* gene family. Most encode proteins with a receptor-like topology and domain architecture (**Fig. 8D**). Despite this similarity, individual Alr proteins have low sequence identity when compared to each other, suggesting the gene family is old, has experienced high rates of molecular evolution, or both.

Our data indicate that the N-terminal domains of Alr proteins are Ig domains. Specifically, domain 1 is either part of the V-set family or represents a new family within the IgSF that is most closely related to V-set domains. Likewise, domains 2 and 3 are either part of the I-set family or represent a new family most closely related to I-set domains. We propose including the Alr domains within existing V-set and I-set families. One reason for doing so is that the sequence profiles often used to identify Ig domains may be biased against recognizing them in non-bilaterians. Indeed, profiles for V-set and I-set domains were originally defined using a handful of sequences, mostly from vertebrates and model organisms (43, 55). Today V-set and I-set profiles in pfam contain thousands of sequences, but are dominated by sequences from chordates and, to a much lesser extent, arthropods (33). Although there is considerable sequence diversity within these sequences, they still represent the descendants of only two evolutionary nodes within metazoans. A consequence, suggested by this study, is that many V-set and I-set Ig domains go undetected in non-bilaterians and other understudied taxa. As genomes continue to be sequenced, our ability to detect these domains has been enhanced by new methods for detecting remote homologs and producing highly accurate structural models (37–39). These new data will undoubtedly enhance our understanding of how Ig domains evolved and may force us to revisit how these families are defined.

Indeed, to our knowledge, this is the first time a V-set Ig domain has been identified outside of bilaterians. Our data therefore suggest the last common ancestor of *Hydractinia* and bilaterians had distinct V-set and I-set Ig domains. These domains likely followed different evolutionary trajectories to arrive at their current sequences. Some Alr I-set domains appear to have subsequently lost the C’ strand. Testing this hypothesis (or alternatives involving convergence) will require careful analysis of Ig domains from other metazoans, especially non-bilaterians.

We have also discovered that the region previously referred to as the ECS actually encodes a domain with an Fn3-like fold. However, the sequences of Alr and Fn3 domains differed substantially between strands D through G, and HHpred did not identify them as homologs with high confidence. It therefore remains unclear whether this Fn3-like domain belongs to the fibronectin III superfamily.

Many Alr Ig domains lack the disulfide bridge and tryptophan found at the core of most Ig domains. Although there are many examples of Ig domains lacking either the disulfide bridge or the tryptophan, we are unaware of any V-set or I-set Ig domains that lack both. The reason for this loss is unclear. Intriguingly, the Alr I-set domains, along with their neighboring Fn3-like domains, are predicted to have conserved disulfide bridges that link neighboring β-strands within one β-sheet. There is also a conserved disulfide bridge between neighboring I-set and Fn3-like domains, which may stabilize their orientation.

One obvious function for the newly described *Alr* genes is allorecognition. It is therefore significant that many *Alr* genes are located outside the genomic region originally mapped by Cadavid et al. (12) and Powell et al. (13). If these other *Alr* genes rendered homozygous in the inbred lines used for mapping, any effect they might have as an allodeterminant would have been missed. In nature, however, these genes might be as polymorphic as *Alr1* and *Alr2*. Such undiscovered allodeterminants could explain the appearance of unexpected allorecognition responses between outbred colonies (14, 15).

*Alr6* is a good candidate for such an allodeterminant because it is very similar to *Alr1. Alr6* occupies a location in cluster A that is roughly syntenic with the location of *Alr1* in Cluster B. It has an *Alr1-*like intron/exon structure, and it encodes a cytoplasmic tail that is clearly homologous to that of *Alr1*. In addition, *Alr6* is alternatively spliced to generate isoforms that have a single Ig domain and an alternative cytoplasmic tail, a feature shared only with *Alr1*. We hypothesize this similarity could extend to *Alr6* functioning as a third allodeterminant in the ARC.

*Alr30* could also be an allodeterminant. *Alr30* is immediately upstream of *Alr2* and was previously considered a pseudogene (15, 19). Here, we show that *Alr30* encodes a transmembrane protein with an extracellular region with recognizable sequence similarity to *Alr2*, but a cytoplasmic tail that has no detectable similarity to other *Alr* genes. Since this gene resides within the genomic interval previously defined in Nicotra et al. (14), it is formally possible that it, too, contributes to allorecognition phenotypes.

The domain architecture of most Alr proteins also suggests alternative functions in extracellular protein-protein interactions. Tandem Ig domains are commonly found in cell adhesion molecules, proteins involved in cell-to-cell communication, and immune receptors (56). An adhesive function would also be consistent with that already described for Alr1 and Alr2 (17).

The ITAM motifs in some Alr cytoplasmic tails are also potentially significant. ITAM-mediated signaling activates inflammation and cellular immune responses in vertebrates (24). It also activates phagocytosis of unwanted or damaged cells in *Drosophila* (23, 57) and appears to promote immune responses to bacteria in oysters (58). An immune function for some ITAM-bearing Alr proteins therefore seems plausible.

Could ITAM-mediated signaling play a role in allorecognition responses? One possibility is that Alr proteins with ITAM motifs activate rejection responses when they bind non-polymorphic, *Hydractinia-*specific ligands on opposing tissues. This rejection response would then be inhibited if polymorphic allodeterminants bind a compatible ligand. At present, this model is only supported by two seemingly disparate observations. First, *Hydractinia* mount the most vigorous and sustained rejection responses against other *Hydractinia* (59). This indicates colonies can identify the type of tissue they encounter. Second, the initial stages of rejection and fusion are morphologically indistinguishable (60). In both responses, nematocytes migrate to the point of contact and arrange their nematocysts as batteries pointed at their opponent. In rejection, the batteries fire, but in a fusion, the nematocytes migrate away as the tissues merge. This suggests rejection could be the default allorecognition response in *Hydractinia*. In this model, Alr proteins with ITAMs would activate rejection, which would be inhibited later by homophilic binding between compatible allodeterminants. This model would be analogous to the balance of ITAM and ITIM-mediated signaling that determines whether natural killer (NK) cells become activated in the vertebrate immune system. If true, it could also indicate a deep evolutionary relationship between invertebrate and vertebrate self-recognition systems.

Our decision to classify some *Alr* sequences as putative genes relied heavily on whether the genes were expressed and correctly spliced. Two caveats are associated with our expression data. First, because the RNAseq experiment was primarily intended to guide our annotation, it did not include biological or technical replicates. The resulting expression levels should therefore be viewed as rough estimates. Second, the RNA used to generate these reads was extracted from a pool of feeding and reproductive polyps. Mat and stolon tissue — the normal sites of allorecognition responses — were not included because we and others have been unable to isolate high quality RNA from these tissues (unpublished data; Uri Frank, personal communication). It is also possible that some *Alr* genes are expressed at developmental time points not represented in our dataset. Therefore, we expect to update our classification with additional data in the future.

The sequences of the Alr genes themselves do not appear to be orthologous to other invertebrate allorecognition genes. Nonetheless, this new ARC sequence reinforces similarities between *Hydractinia* and other species in which allorecognition is controlled by genomic clusters of related genes (5). This clustering may enhance the efficiency and coordination of allorecognition gene expression. It may also facilitate the generation of sequence diversity via gene conversion or unequal crossing over. Moreover, these invertebrate allorecognition complexes are remarkably similar to the complexes that control self/non-self recognition in vertebrates, namely the major histocompatibility complex (MHC) (61), leukocyte receptor complex (LRC) (62), and NK complex (NKC) (63). Identifying evolutionary links between these systems may become possible in the future as we survey a broader swath of metazoan genomes and simultaneously deepen our molecular understanding of how invertebrate allorecognition works.

## Materials and Methods

### Sequencing and assembly of ARC

Colony 236-21 was maintained on glass microscope slides in 38 liter aquaria filled with artificial seawater as previously described (64). DNA was extracted as detailed in the *SI Appendix*. PacBio and Illumina libraries were constructed and sequencing performed at the NIH Intramural Sequencing Center (NISC) via a whole-genome shotgun approach. All raw reads are available through BioProject PRJNA802249. The genome was assembled with the Celera Assembler version 8.3r2 (65). We then used NUCmer from the MUMmer package (ref) to align the resulting assembly to the previously sequenced ARC BACs, and merged these sequences to create a reference sequence for the ARC. Genome sequencing and assembly are detailed in the *SI Appendix*.

### RNA extraction, sequencing, and mapping

RNA was harvested and extracted from a mixture of gastrozooids and gonozooids as detailed in the *SI Appendix*. RNA-Seq libraries were constructed and sequenced at NISC as detailed in the *SI Appendix*. Raw reads are available through BioProject PRJNA802249. To calculate expression levels of our annotated Alr genes, paired-end RNA-seq reads were mapped to the entire genome assembly using HISAT2 (66) and transcript abundance estimated with Cufflinks (67) with a correction for multiple read mappings as detailed in the *SI Appendix*.

### Annotation of *Alr* genes and sequence comparisons and analyses

*Alr* genes were annotated using Apollo (68). Methods for multiple sequence alignments, pairwise sequence alignments, sequence clustering, protein sequence annotation, and phylogenetic tree construction are provided in the *SI Appendix*.

### Structural prediction and alignment

For single domain predictions, we generated a custom multiple sequence alignment, as detailed in the *SI Appendix* which was submitted to Colabfold via the “AlphaFold2_mmseqs2” notebook, version 1.1 (37). The secondary structure of each model was determined with STRIDE (44). Structure comparisons were performed with DALI (41) and PDBeFOLD (https://www.ebi.ac.uk/msd-srv/ssm/) (42). Models were visualized in Pymol 2.3 (69). Further details can be found in the *SI Appendix*.

## Acknowledgements

We thank Leo Buss for sharing colony 236-21 and for helpful discussion. This work was funded by National Science Foundation grant 1557339 awarded to (MLN), National Science Foundation grant 1923259 awarded to (CES, MLN), National Institutes of Health, Intramural Research Program of the National Human Genome Research Institute, grant ZIA HG000140 (ADB), and National Institutes of Health, Intramural Research Program of the National Human Genome Institute, grant ZIA HG200398 to (AMP). ALH and SMS were supported by National Institutes of Health grant T32 AI074490.

## Notes

### Competing Interest Statement

The authors have declared no competing interest.

### Summary of Updates

In this version, the title and abstract were revised, and several figure panels and the detailed methods were were moved to the supplement.

