## Supplemental Data for "A family of unusual immunoglobulin superfamily genes in an invertebrate histocompatibility complex"

PNAS

Affiliations:

**This PDF file includes:**

Supplementary Materials and Methods
Figures S1 to S18
Tables S1 to S10

Legends for Datasets S1 to S15

SI References

**Other supplementary materials for this manuscript include the following:**

Datasets S1 to S15

### Supplementary Information Text

### Material and Methods

#### Sequencing and assembly of the genome of an ARC homozygous animal

Colony 236-21 was maintained on glass microscope slides in 38 liter aquaria filled with artificial seawater (Reef Crystals) as previously described (Cadavid et al., 2004). It was starved three days prior to nucleic acid extraction. Tissue was scraped from the slide with a sterile razor blade and snap-frozen by transferring it to a mortar filled with liquid nitrogen. The frozen tissue was then ground into a fine powder with a pestle. UEB1 buffer (7 M urea, 0.3125 M NaCl, 0.05 M Tris-HCl, 0.02 M EDTA, and 1% w:v N-lauroylsarcosine sodium salt) was added to the mortar, where it froze. The frozen UEB1-tissue mixture was ground into a powder and transferred to a 50 ml centrifuge tube containing room temperature UEB1 buffer. This was mixed by gentle inversion. An equal volume of equilibrated phenol:chloroform:isoamyl alcohol (25:24:1) was added and mixed by gentle inversion. This was centrifuged for 10 minutes at 3000 x g. The aqueous layer was transferred to a 15 ml centrifuge tube with a wide bore pipette tip. Total nucleic acid was precipitated by adding 0.7 volume isopropyl alcohol. Precipitated nucleic acid was then spooled onto a pipette tip and transferred to a clean 15 ml tube, where it was washed twice with 70% EtOH and twice with 100% EtOH. The precipitated material was then gently brought to the bottom of the tube by briefly centrifuging, air dried, and immediately resuspended in 1X TE (10 mM Tris-HCl, pH 8.0; 1 mM EDTA, pH 8.0). RNA was then digested by adding RNAses (RNAse cocktail, Ambion, #AM2286) and incubating at 37°C for 15 minutes. DNA was then extracted by adding 1 volume equilibrated phenol:chloroform:isoamyl alcohol, centrifugation at 12,000 x g, and transfer of the aqueous layer to a new tube. This was followed by precipitation with 2.5 volumes of 100% ethanol and 1/10 volume 5 M sodium acetate (pH 5.2). The precipitate was pelleted, washed with 70% ethanol, and resuspended with 1X TE. The resuspended DNA was then stored at -20°C.

PacBio and Illumina libraries were constructed and sequencing performed at the NIH Intramural Sequencing Center (NISC) via a whole-genome shotgun approach. Both the high-throughput Illumina HiSeq2500, run as 250 base paired-end reads, and PacBio RSII long-read sequencing platforms were used. Filtered subreads from two PacBio libraries (a total of 37 SMRT cells) were corrected with the Celera Assembler version 8.3r2 (Berlin et al., 2015). Specifically, the PBcR.pl script was used with parameter sensitive=1, which is recommended for datasets with <50x coverage, increases MHAP sensitivity, and uses the slower but more accurate pdbagcon consensus algorithm to generate the corrected reads. The corrected reads were assembled with runCA.pl parameters batOptions=”-el 5000 -eg 0.025 -Eg 4.00 -em 0.025 -Em 4.00 -o asm -RS -NS -CS -repeatdetect 6 150 15” which reduces the default error rate, increases the minimum overlap size, and increases the splitting thresholds. The resulting assembly was polished with the PacBio reads using the ArrowGrid parallel wrapper (Chin et al., 2013) followed by polishing with the Illumina short read data using the PilonGrid parallel wrapper (Walker et al., 2014).

### Assembly of the ARC

To assemble the full reference sequence for the ARC, NUCmer from the MUMmer package (v3.23) was used align the BAC contigs with the newly assembled whole genome sequence to identify the contigs which matched the known ARC sequence (Kurtz et al., 2004). First, the query and reference sequences were aligned using NUCmer (nucmer -p <output.file> <reference.file> <query.file>). The resulting file was then filtered (delta-filter) to only show matching hits in one direction on the strands (-r) and to remove all hits less than 1000 base pairs (-l #). Finally, the output was appended into a tab-delimited file (-T) sorted by the reference sequence (-r), with a minimum length of 1 kb or 10 kb (-L #), the sequence length (-l), and the percent coverage between two sequences (-c). The tabular files were manually inspected to assess overlapping contigs. Overlapping regions were then inspected by alignment with BLAST+ version 2.6.0 (Camacho et al., 2009) and dot plots generated in YASS (Noe and Kucherov, 2005). The genome assembly and BAC sequences were then merged to create a reference sequence of the ARC-F haplotype (File S1).

### RNA extraction, sequencing, and mapping

Thirty polyps were severed from colony 236-21 with a scalpel, moved to an eppendorf tube and briefly centrifuged. Remaining water was removed with a pipette. Tissue was immediately lysed with 0.5 mL of TRIzol (Invitrogen) and ground vigorously with a small pestle. Lysate was incubated for less than five minutes at room temp. One hundred µl chloroform was added and the tube was shaken vigorously for 15 seconds, followed by a three minute incubation at room temp. The sample was then centrifuged at 12,000 x g for 15 minutes at 4°C. RNA was then extracted from the aqueous phase with a PureLink RNA Mini Kit (Invitrogen). RNA quality and quantitation was assessed by Tapestation and Qubit, respectively, at the University of Pittsburgh Genomics Core. Final sample was frozen and stored at -80°C until sequencing by NISC. RNA-Seq libraries were constructed from 1 µg RNA using the Illumina TruSeq Stranded mRNA kit. The resulting cDNA was fragmented using a Covaris E210 focused ultrasonicator. Library amplification was performed using 10 cycles to minimize the risk of over-amplification. The library was sequenced on an Illumina HiSeq4000 to generate 75 base paired-end reads.

To calculate expression levels of our annotated Alr genes, paired-end RNA-seq reads were mapped to the entire genome assembly using HISAT2 (Kim et al., 2019). The mapping was performed under the most stringent conditions (only concordant mappings with zero mismatches were kept) while allowing for multiple alignments. The resulting mapping file was processed and sorted using samtools (Li et al., 2009) before proceeding to quantitation. Using the reference annotations of the Alr genes, transcript abundance of the Alr genes was estimated with Cufflinks (Trapnell et al., 2010). Abundance estimates were corrected for multiple read mappings.

### Annotation of *Alr* genes

*Alr* genes were annotated using Apollo (Dunn et al., 2019) installed on a local computer running Ubuntu 18 LTS. Tracks displaying the results of BLASTX searches and RNAseq mapping were imported and used as a guide for manual annotation of *Alr* gene models. To generate BLAST results, repeats in the genomic sequences were first masked using the protein-based repeat masking option on the Repeatmasker website (https://www.repeatmasker.org) (Smit et al., 2015). Masked DNA sequences were then divided into 32 kb segments with 2 kb overlaps. These segments were used as BLASTX queries against a database of Alr1 and Alr2 proteins (to identify *Alr*-like sequences), and the swissprot database (to identify highly conserved genes). BLAST results were then concatenated, and a custom perl script was used to adjust their coordinates to those of the unsegmented genome sequence. To generate RNAseq alignments, the assembled RNA-seq dataset was aligned to the genome using HISAT2 (v2.1.0) through the Galaxy platform (Kim et al., 2015, 2019). The parameters used RNAseq alignments included paired-end reads and no alignments for individual mates, and only 1 primary alignment. The output file (.bam) was then imported to Apollo for visualization during annotation.

**Alr sequence comparisons**

Alignments between Alr proteins were performed using MAFFT (Katoh and Toh, 2010). The L-INS-i alignment strategy was used for all alignments except those involving only domains 1, 2, and 3, which used G-INS-i. Pairwise sequence alignments were done using the modified Needleman-Wunsch algorithm available in Jalview [15]. Clustering was performed with CD-HIT [16] using the psi-cd-hit.pl script, with 20% sequence identity cutoff and 0.1 e-value cutoff. Neighbor joining trees were constructed in Jalview using the BLOSUM62 scoring matrix. Trees were visualized in iTOL [17], exported as scaled vector graphics files, and annotated in Adobe Illustrator.

**Protein sequence analysis**

Signal peptides were predicted with SignalP 5.0 [18]. Transmembrane helices were predicted with TMHMM 2.0 [19]. Conserved protein domains were identified with the Pfam database [20] using HMMER3 (http://hmmer.org/). For domain prediction by HHpred, sequences were submitted to the MPI Bioinformatics Toolkit [21]. The query MSA was generated via three iterations of HHblits against the Uniref30 database, with an e-value threshold of $1 \times 10^{-3}$ for inclusion. HHpred was then used to search the SCOPe70_2.07 database.

**Structural prediction and alignment**

For single domain predictions, we generated a custom multiple sequence alignment which was submitted, along with the query sequence, to Colabfold via the “AlphaFold2_mmseqs2” notebook, version 1.1 [22]. The input multiple sequence alignment was generated as follows. The sequence of the query domain was aligned to the same domain type from all *bona fide* Alr proteins using MAFFT with the G-INS-i setting. This alignment was then submitted as a query to HHblits via the MPI Bioinformatics Toolkit, which was run for two iterations against the Uniref30 database, with an e-value threshold of $1 \times 10^{-3}$ for inclusion. A reduced representation alignment of the resulting Query MSA was then downloaded and submitted as a custom multiple sequence alignment to Colabfold. The secondary structure of each model was determined with STRIDE [23]. The Alr structural model with the highest average pLDDT was then submitted to DALI [24] and PDBeFOLD (https://www.ebi.ac.uk/msd-srv/ssm/) [25] to identify similar structures in the PDB. For multi-domain predictions, the query sequence was submitted directly to Colabfold, with msa_mode = “Mmseqs2 (Uniref + Environmental)”. Models were visualized in Pymol 2.3 [26].

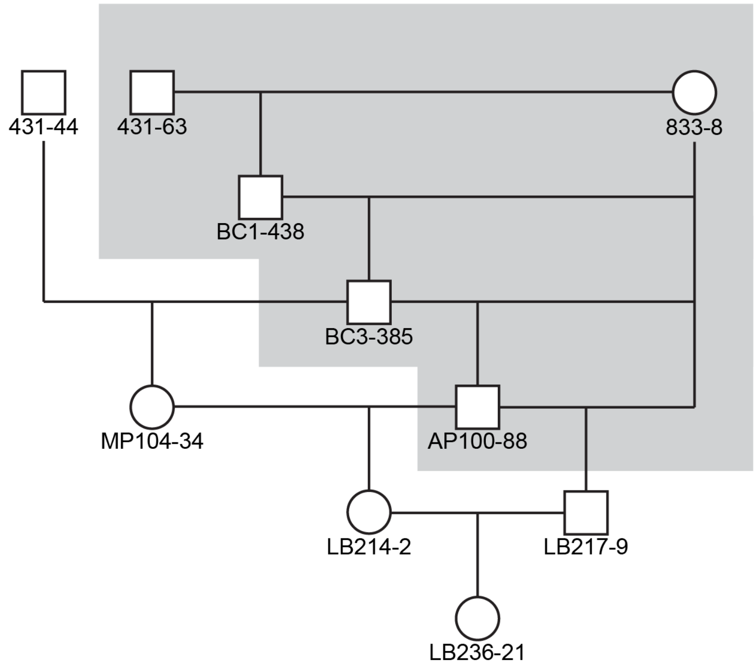


**Fig. S1. Pedigree of colonies used to generate ARC-F reference sequence.**
The pedigree of colony LB236-21 can be recreated by concatenating previously published pedigrees (shaded area) [1,27]. Colony AP100-88 is from the mapping population in Powell *et al*. [27]. Colony 431-44 is from the mapping population in Cadavid *et al*. [1].

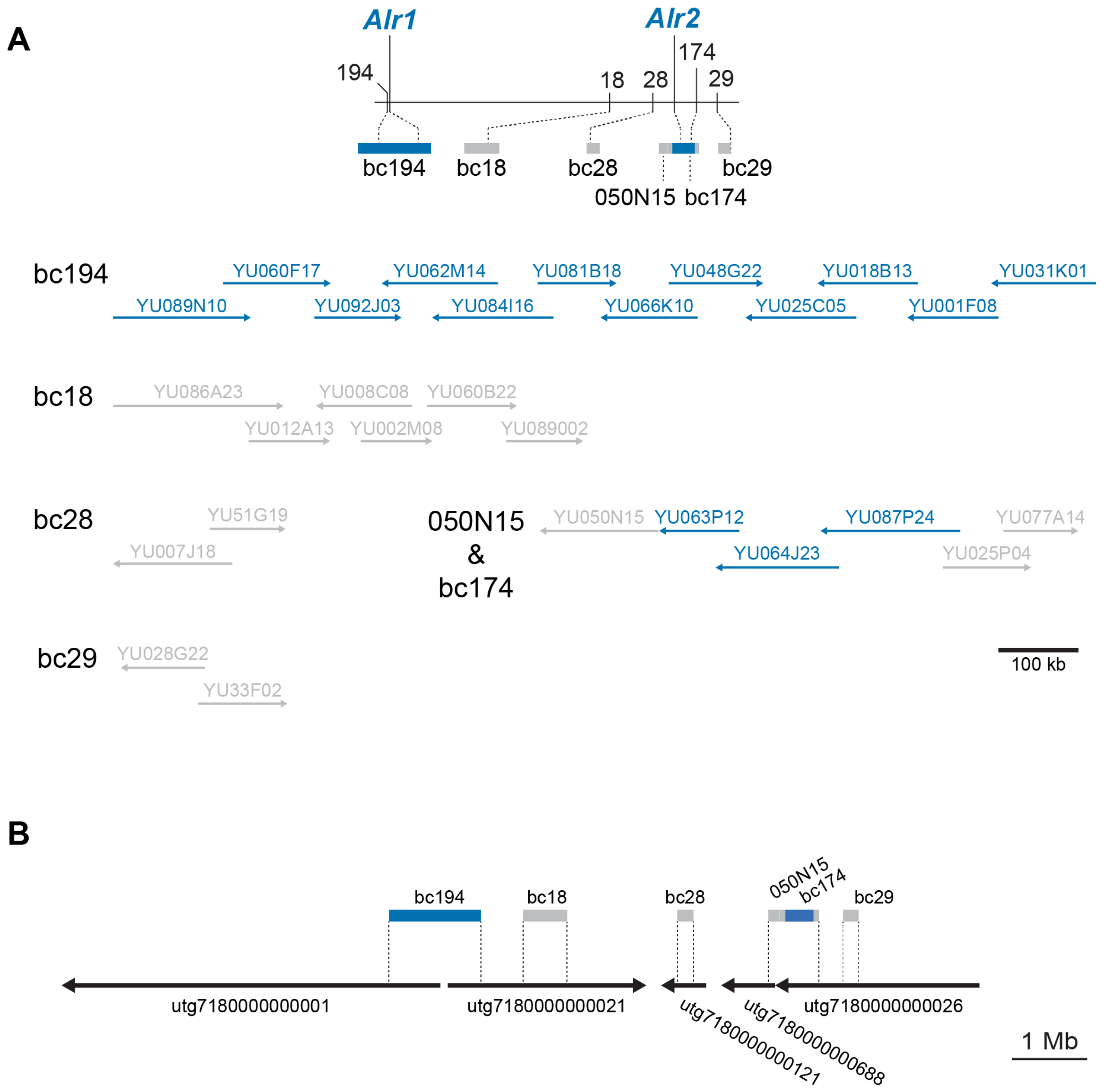


**Fig. S2. Detail of ARC reference assembly.**

(A) Minimum tiling path of sequenced BAC clones resulting from chromosome walks from five markers in the ARC linkage map. Clone names are indicated above an arrow indicating their orientation. Sequences reported in [28] or [29] are in navy blue. Unpublished sequences are in gray.
(B) Overlap between BAC contigs and genome contigs.

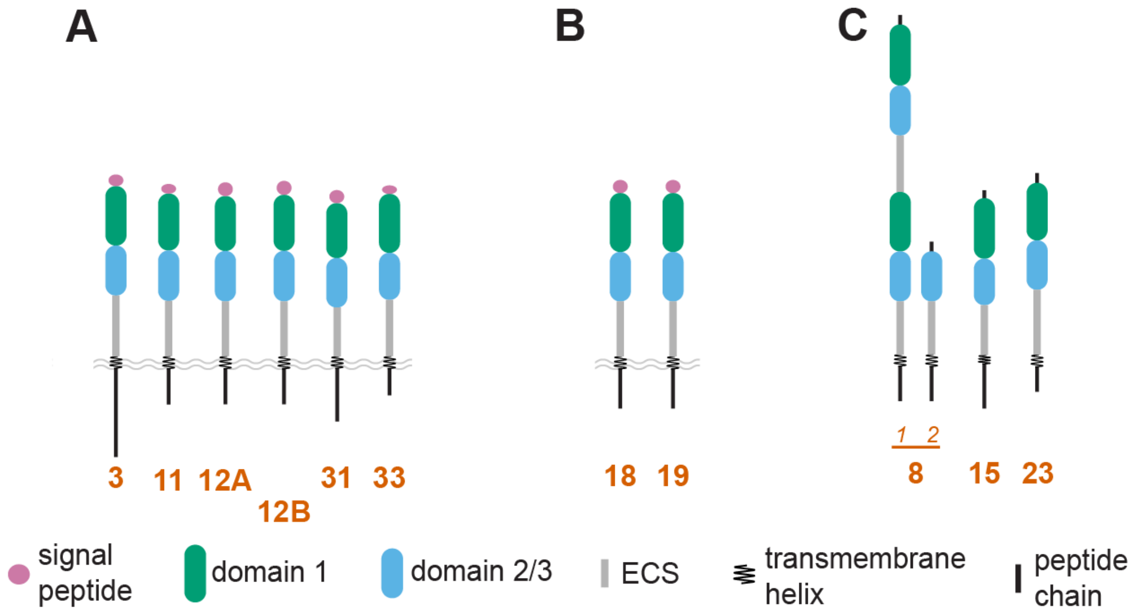


**Fig. S3. Predicted domain architectures encoded by putative *Alr* genes.**
(A) Domain architecture of unexpressed putative genes.
(B) Domain architecture of partially expressed putative genes.
(C) Domain architecture of putative genes lacking a predicted signal peptide.
In (A) and (B), signal peptides are expected to be cleaved, but are shown to indicate their presence.

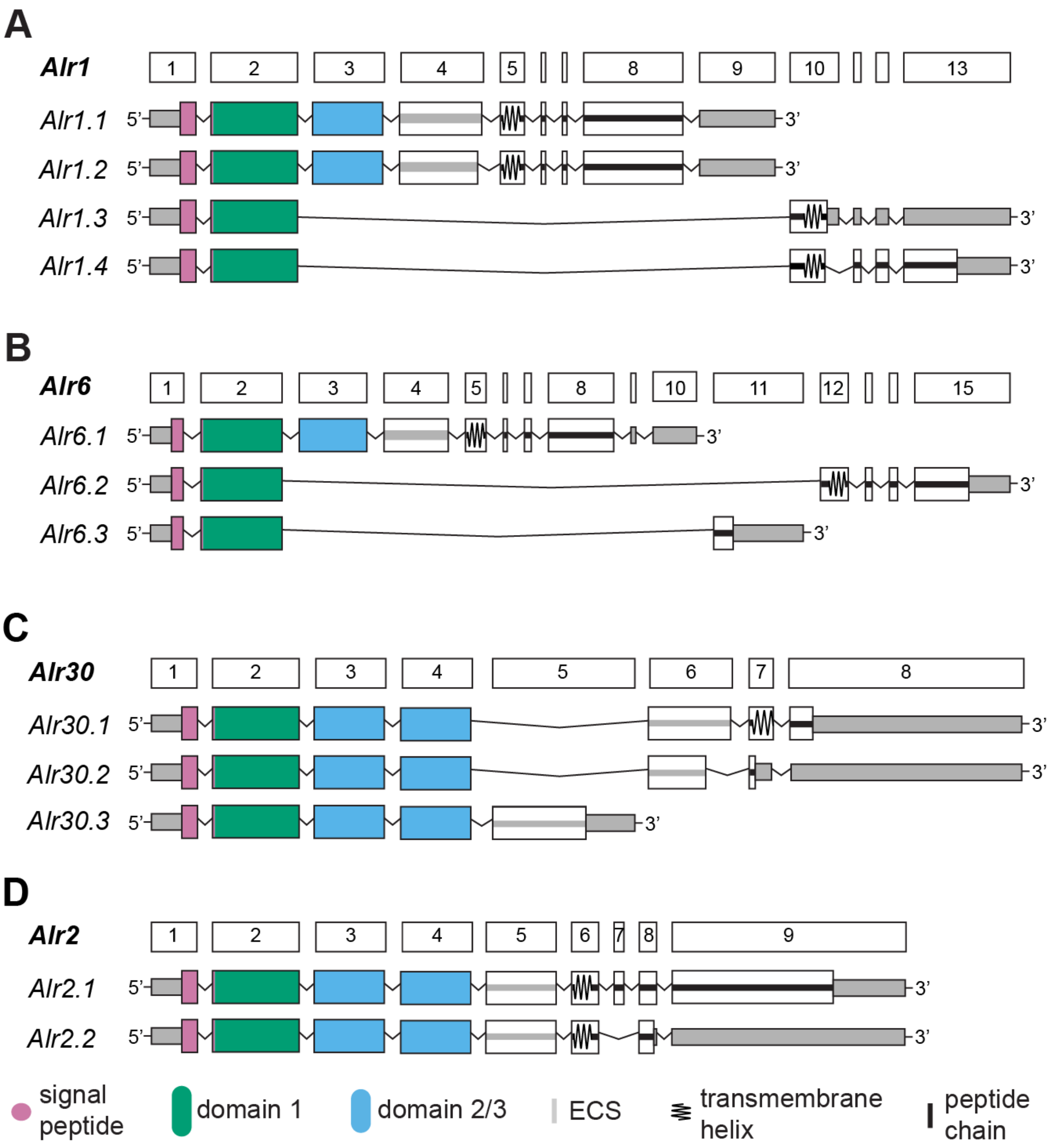


**Fig. S4. Alternative splicing of *Alr* genes**
In all panels, exons are colored according to the type of domain/region they encode. (A) alternative splicing of *Alr1*. (B) Alternative splicing of *Alr6*. (C) Alternative splicing of *Alr30*. (D) Alternative splicing of *Alr2*.

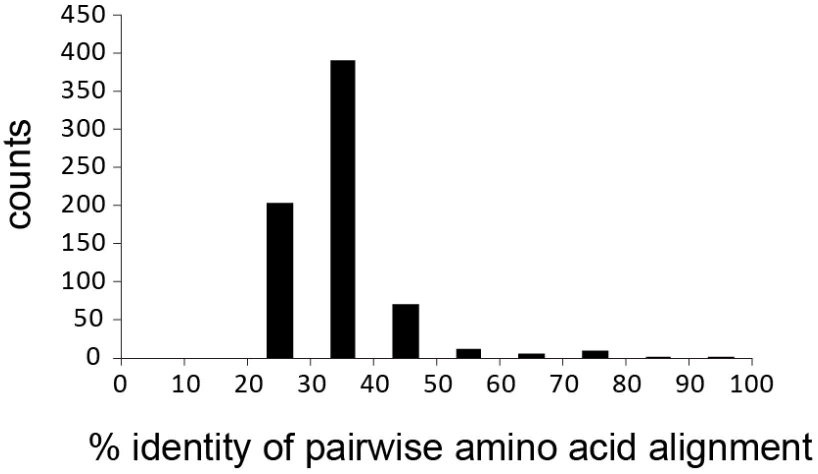


**Fig. S5. Pairwise amino acid identities between *Alr* genes**
Histogram of amino acid percent identities for pairwise alignments of *Alr* genes and putative genes. Alignments were performed using the modified Needleman-Wunsch algorithm available in Jalview.

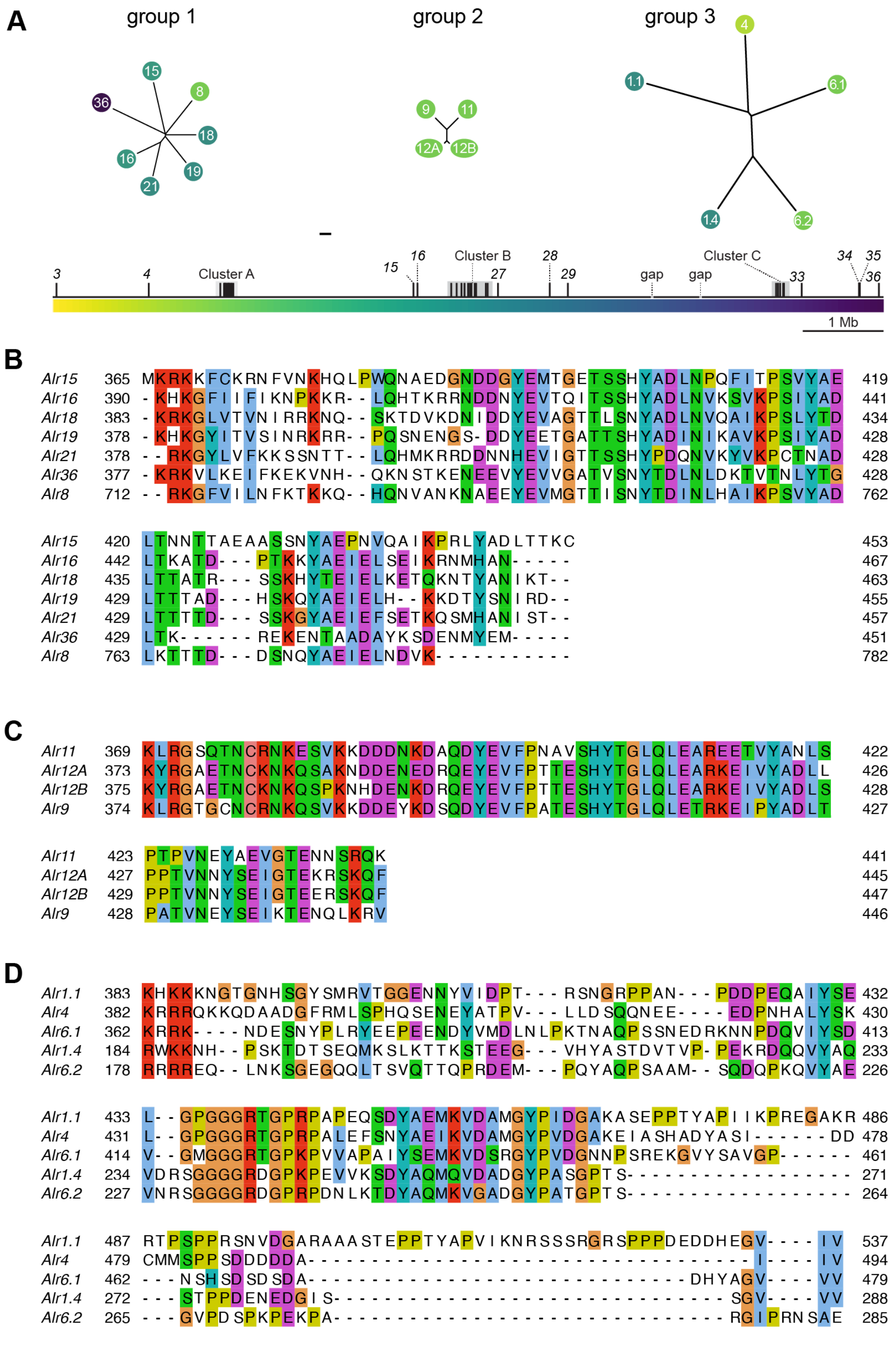


**Fig. S6**

(A) Alr cytoplasmic tails grouped by CD-HIT at 20% similarity. Neighbor-joining trees are shown. Leaves are color coded according to their genomic position. Branch lengths calculated according to the BLOSUM26 matrix. Scale bar = 100 units.

(B) Alignment of group 1 cytoplasmic tails.

(C) Alignment of group 2 cytoplasmic tails.

(D) Alignment of group 3 cytoplasmic tails.

**A**

```
Alr16   390  -KHKGFIIFIKNPKKR--LQHTKRRNDDNYEVTQITSSHYADLNVKSVKPSIYADLTKATD---PTKKYAEIELSEIKRNMHAN----- 467
Alr21   378  --RKGYLVFKKSSNTT--LQHMKRRDDNNHEVIGTTSSHYPDQNVKYVKPCTNADLTTTTD---SSKGYAEIEFSETKQSMHANIST-- 457
Alr36   377  -KRKVLKEIFKEKVNH--QKNSTKENEEVYEVVGATVSNYTDLNLDKTVTNLYTGLTK------REKENTAADAYKSDENMYEM----- 451
Alr8    712  --RKGFVILNFKTKKQ--HQNVANKNAEEYEVMGTTISNYTDINLHAIKPSVYADLKTTTD---DSNQYAEIELNDVK----------- 782
Alr15   365  MKRKKFCKRNFVNKHQLPWQNAEDGNDDGYEMTGETSSHYADLNPQFITPSVYAELTNNTTAEAASSNYAEPNVQAIKPRLYADLTTKC 453
Alr18   383  -KRKGLVTVNIRRKNQ--SKTDVKDNIDDYEVAGTTLSNYADLNVQAIKPSLYTDLTTATR---SSKHYTEIELKETQKNTYANIKT-- 463
Alr19   378  -KHKGYITVSINRKRR--PQSNENGS-DDYEETGATTSHYADINIKAVKPSIYADLTTTAD---HSKQYAEIELH--KKDTYSNIRD-- 455
```

**B**

```
Alr9    374  KLRGTGCNCRNKQSVKKDDEYKDSQDYEVFPATESHYTGLQLETRKEIPYADLTPATVNEYSEIKTENQLKRV  446
Alr11   369  KLRGSQTNCRNKESVKKDDDNKDAQDYEVFPNAVSHYTGLQLEAREETVYANLSPTPVNEYAEVGTENNSRQK  441
Alr12A  373  KYRGAETNCKNKQSAKNDDENEDRQEYEVFPTTESHYTGLQLEARKEIVYADLLPPTVNNYSEIGTEKRSKQF  445
Alr12B  375  KYRGAETNCKNKQSPKNHDENKDRQEYEVFPTTESHYTGLQLEARKEIVYADLSPPTVNNYSEIGTEERSKQF  447
```

**C**

```
Alr31   357  SVTIKELEENENVEENNDYKDLNCLDISEETYSKLTPPKSRLHKARVQLRYSKGKIKSFKKHSTKSTHEYEVPPSLLANMSKDKTAD  463
```

**Figure S7. ITAMs in putative Alr proteins**

(A) Alignment of group 1 cytoplasmic tails, showing ITAMs (orange shading). Overlapping ITAMs are shown with heavier shading. Bona fide *Alr* genes (blue gne names) are included in the alignment for comparison.

(B) Alignment of group 2 cytoplasmic tails, showing ITAMs. Shading as in (A).

(C) ITAM in putative *Alr* gene *Alr31.*

```
Alr1 Domain 1
 |A-|   |A'|      |--B--|          |--C--|   |--C'-|  |C"|            |-D--|    |--E-|          |--F--|     |----G----|
LNVKPVLPTISTTFNATVELQWQVTLEPSEAIASMTVRELPSLITILTGAVDGFNVAQGGRKLFGDRVSGIFNNRNNIVTLTLQNIQYNETLTFQLLVGSSTLVKAANISIVEIS
 EEEETTTTEEEETTTb EEEEEEEE TTTT   EEEEEEEETTTEEEEEEEETTEEEE HHHHHHHTTTEEEEEEETTTTEEEEEEETTbTTTT  EEEEEEEETTTEEEEEEEEEEEE

Alr2 Domain 1
  A    |A'|      |--B--|           |--C--|  |-C'-|    C"          |D|        |- E-|        |---F---|   |---G----|
LSLTSPAIIEEVVGRSVTITYVTDVADVADDVVVNFRINYNDSRIAEGTKTVFDKIPPTPFGNRLRTSTPLQNQKEYSLLIDNLEYNDTGLFFAKIEIFKPNVEANSTTNLIIY
 EE TTEEEETTTb EEEEEEE  TTTTTTT EEEEEEEETTEEEEEETTTTEEETTTTTTTTTTEEE   TTTT EEEEEETTb GGG EEEEEEEEETTTTEEEEEEEEEEE

Alr3 Domain 1
 |A|    |A'|    |--B---|    |---C---|        |-C'-|  |C"|            |-D--|  |-E--|        |----F----|       |----G-----|
AVLYTDSPTISGIFNQSAIISFKATITANDNRTVKSFYVELPPSTQQIAIGSNNVLQALPVPPFNSRLTASAANQEYLLQVNPLKFSDDYTFRGVLIYLLNNDINNPIAVEQDVKLDVF
 EEETTTTEEEETTTb EEEEEEEE TTTT EEEEEEEEEETTTTT EEEEEEETEEEETTTTTTTTTTTEEEEEEETTEEEEEEETTbTTTTTEEEEEEEEEEEEETTTTTTTEEEEEEEEEEEE

Alr4 Domain 1
 |A|     |A'|   |--B---|      |---C----|    |C'-|    C"           |-D--|    |---E--|        |----F----|       |----G-----|
NVEFLNGQSEVTGMLDKDITLQWQITFLKGEMLQSHDIYLPNRTKIVSNQPPELTPVGKRMYGTRLVPVFDADAAVFKLLKNVKFTDSSHNFTLVVAFERKDDFNRRTGVADINIVNVE
 EEEGGG  EEEETTTEEEEEEEEEE   TTTEEEEEEEEEEEETTTTEEEEEEEETTTTTE  HHHHHHHTTEEEEEEEGGG EEEEEEEEETTTTTTTEEEEEEEEEEEETTT EEEEEEEEEEEE

Alr6 Domain 1
 |A-|   |A'|      |--B--|     |---C----|     |-C'-|    C"            |--D-|  |-E--|          |----F----|  |-----G------|
GTVEAVKTNFEVPVNQTAQLQWRVVPDAGEQVFGVEVFVLGLPNVQIIKLTSTVTIAGKERFGNRISGSLTNGIYTLSIKKIQFNEKKSFNLKVVFYKEPDYYPKNSTVVIKKVV
 EEEETTTTEEEETTTb EEEEEEEE TTTTEEEEEEEEEETTTTEEEEEETTTTEE  HHHHHHHTTTEEEEEEETTEEEEEETTbTTTT EEEEEEEEEETTEEEEEEEEEEEEE

Alr7 Domain 1
    |-A|   |A'-|     |---B--|     |----C----|      |-C'-|   |C"|          |--D-|     |-E--|         |----F----|     |-----G-----|
ASSGKVTVVVKNVFERVQIKSTVVMEWKIAHSDNQTNKVLKLYVLPDRDRPVFSSYGKYQSSQGKGRQTFGNRLSATFSKIKGKYTVTLKRIQYNENYTFQLKVIFINKKSVTETKVADIRIKNVV
  EEEETTTEEEETTTb EEEEEEEE TTTEEEEEEEEEEEETTTTTEEEEEEEETTEEEE HHHHHHHHTTTEEEEEEGGG EEEEEETTbTTTT EEEEEEEEEEEETTT EEEEEEEEEEEE

Alr8 Domain 1a
  |A|    |A'|     |--B---|      |---C----|      |C'-|   C"             |--D-|  |--E-|         |---F----|        |----G-----|
YGGTITAAATTVTVNFQSTLKLEWTAIPSPAEQIVVLKVFILPDTNNGVLTNTNPPVLLPTGMSLFGRNRLSATFINSKYTLMLKNVTYNDSCTFQMYAIFRKPAAYSVYQVNRSNIKVIVT
  EEEETTTEEEETTTb EEEEEEEE TTTTEEEEEEEEEETTTTTEEEEEETTTEEE HHHHHHHTTTEEEEEEETTEEEEEETTb GGG EEEEEEEEETTTTTTTEEEEEEEEEEEE

Alr8 Domain 1b
 |-A-|    |A'|    |--B--|      |----C---|       |C'|     C"           |-D--|  |-E--|          |----F----|  |-----G-----|
VNDETISDTASTLDIVYNTTLKLDWNMTLSAGEVIVGVQVYVLPDTTNRIITDTSPTVLSKGISIFGENRLSATFTNSRYRLMIRNIRYNESFTFQLYVIYGAGDHLSSSRFNIQVTVK
 EEEEE  TTEEEETTTbEEEEEEE   TTTTEEEEEEEEEEEETTTTTEEEEETTTEE HHHHHHHTTTEEEEEEETTEEEEEETTbTTTT EEEEEEEEEEETTEEEEEEEEEEEEE

Alr9 Domain 1
 |A-|   |A'|      |--B--|       |---C---|   |-C'|    C"            |--D-|  |--E-|          |----F---|   |-----G------|
GSVEAIIKNIEATDNSTAELSWRVETNKDGERVFGVELSEGGVVVILDKSGTVTEAGRNKFGGRLSASFSNNVYKMFIKKIQYNEAKSFTLTAAFYWKSALDPVNDTATITSVK
 EEEETTTEEEETTTb EEEEEEEE  TTTTEEEEEEEEEETTEEEEEETTTTEE HHHHHHHTTTEEEEEEETTEEEEEETTbTTTT  EEEEEEEEEETTEEEEEEEEEEEEE

Alr11 Domain 1
 |A-|   A'       |--B--|       |----C---|    |C'-|    C"            |--D-|  |-E--|          |--F----|    |-----G------|
GTVEAVKTNLELPFNETLNLYWRIVLDVGEEISTANVYVLGSPNVQIIKFISTITAAGNAMFGNRLSGSLSKDIYALSIKNIQYNEQKSFNLKVIFQSPVLHAKNATVVIKEVV
 EEETTT EEETTTb EEEEEEEE TTTTEEEEEEEEEEETTTTEEEEEEETTEEE HHHHHHHTTTEEEEEEETTEEEEEETTbTTTTT EEEEEEEEEETTEEEEEEEEEEEEE

Alr12A Domain 1
   |A-|      |-B--|        |---C----|      |C'-|    C"           |-D--|  |-E--|          |----F----|   |-----G------|
VKTNLEVPVNQTAQLQWRVVPVAGEQVFGVEVFVLGSTNVQIIKLTSTLTISGKKRFGNRLSGSLRNGIYTLSIKKIQFNEKKSFNLKVVFYKAPEYYPKNSTVVIKEVV
   EEEETTTb EEEEEE  TTTTEEEEEEEEEETTTTEEEEEEETTEE HHHHHHHTTTEEEEEEETTEEEEEETTbTTTT EEEEEEEEEEETTEEEEEEEEEEEEE

Alr12B Domain 1
 |-A--|     |-B--|        |---C---|      |--C'-|  |C"|            |-D--|  |-E--|          |---F---|    |----G-----|
VKTNLEVPVNQTAQLKWRVVLDLGEKITTVNVFILESPKVLIVLGTHLASATTPAGKTMFGDRLSGSLSNGIYTLSIKKIQFSEEKSFNLEVLFESQPFYLKNATVVIREVG
 EEEEETTTb EEEEEE  TTTTEEEEEEEEEEETTTTEEEEEEEETEEEE HHHHHHHTTTEEEEEEETTEEEEEEETTbTTTTT EEEEEEEEEETTTEEEEEEEEEEEE

Alr15 Domain 1
    |A-|     A'     |--B---|     |--C---|  |-C'-|         |-C"-|  |---D----|   |---E---|        |----F----|    |-----G------|
DGQIAVTLANNKKEASKGKDCTVYWNISNAENIFHLKLYRNKTNIKSNNPASWKSLELFTVNKTSISIETFMEGDKMCYMMTIHNVSYADDASEYVMKVIYVTLEPFQKYEEKATFHLNVI
    EEEETTT  EETTTb EEEEEEEETGGGEEEEEEEEETTEEEEEE GGGTTTEEEEEETTEEEEEEEEEETTTEEEEEEEEEETb GGGTTEEEEEEEEEEEETTTTTEEEEEEEEEEEEE

Alr16 Domain 1
 |A-|   |A'|      |--B--|      |---C---|      |-C'-|   |C"|            |-D--|   |-E--|          |----F----|   |-----G-----|
ADVRISTSFLEATYGNTVDMHWKIILDTGQTINSFTLSLKSRPEDSIIFGSANYQNIANKGKELFGNRLSAVYDKTTSTYRVSIKNIQYNETLSFQLTTTLSNPLFGKQTTIEIKDVK
 EEEETTTEEEETTTb EEEEEEEE TTTTEEEEEEEEEETTTTTEEEEEEEETEEEE HHHHHHHTTTEEEEEEGGG EEEEEETTbTTTT EEEEEEEEEEETTEEEEEEEEEEEEE

Alr17 Domain 1
    |A-|   |A'|     |--B---|      |---C----|      |-C'-|  |C"|          |--D-|     |--E-|         |---F---|    |-----G----|
HVSEVKPVKSVLEVTHGSTVNMKWNITLSKNQVITGFSLVVLPDVGNPVVSGNVNSQMVQEKGRELFGNRLSATFNKTAGLYIATLGNIQDNETYAFRLMTTFSLPDQLEGNIIEIRNIT
   EEEETTTEEEETTTb EEEEEEEE TTTTEEEEEEEEEETTTTTEEEEEEEETEEEE HHHHHHHTTTEEEEEEGGG EEEEEETTbTTTT  EEEEEEEEEETTTEEEEEEEEEEEEE

Alr18 Domain 1
 |A-|   |A'|      |--B--|         |---C---|  |--C'-|  |C"|            |--D-|    |--E-|        |----F---|    |---G----|
LNVKPVLPTISTTFNATVELQWKVTLEPSEAIASMTVRELPSLIGILTGTVDGFNVAQGGTKLFGDRISGIFNNSSNIATLTLQNIQYEETLTFELVVVSSKFVIKAANISIEQIS
 EEEETTTTEEEETTTb EEEEEEEE TTTT EEEEEEEETTTEEEEEEEETTEEEE HHHHHHHTTTEEEEEEETTTEEEEEEEEE   bTTTTEEEEEEEETTT EEEEEEEEE

Alr19 Domain 1
 |A-|   |A'|      |--B--|          |--C--|   |--C'-|  |C"|            |-D--|    |-E--|          |--F--|     |---G----|
LNVKPVLPTISTTFNATVELQWQVTLEPSEAIASMTVRELPSLITILTGAVDGFNVVQGGRKLFGDRVSGIFNNRNNIVTLTLQNIQYNETLTFQLLVGSSTLVKAANISIVEIS
 EEEETTTTEEEETTTb EEEEEEEE TTTT   EEEEEEEETTTEEEEEEEETTEEEE HHHHHHHTTTEEEEEEETTTTEEEEEEETTbTTTT  EEEEEEEETTTEEEEEEEEEEEE

Alr21 Domain 1
 |A-|   |A'|      |--B--|      |----C---|      |-C'-|   |C"|           |-D--|    |--E-|         |---F----|   |-----G----|
ASVEFIAMTKCAVTNTTVKLTWKVKLEPSEGIIYIKLHKLPDDENNIVTYASQKLTVNIKGRQLFGKRLSANYSKHYENVTLTLQNIKYNENVTFYFVARFFPGFVLGTGAITIKCVT
 EEEETTTEEEETTTb EEEEEEEE TTTTEEEEEEEEEETTTTTEEEEEEEETEEEE HHHHHHHTTTEEEEEEGGG EEEEEETTbTTTT EEEEEEEEEEETTEEEEEEEEEEEEE

Alr23 Domain 1
 |-A-|   |A'|      |---B---|    |---C----|  |--C'--|   C"            |-D--|  |-E--|         |----F---|    |----G-----|
NGVTITTKSNNKIVLNGNTLELEWTVNVSPGDTIALRSVYLSPDFISPILVDGSVIEKGRELFGGNRLSATYGNTTYTMSLMDVRYNESGTFRLFVFFRNNGLKETKSDIHVRVT
 EEEEETTTEEEETTTb EEEEEEEEETTTTEEEEEEEEEEEEEEEEEEEEETTEE HHHHHHHTTTEEEEEEETTEEEEEEETTbTTTTTEEEEEEEEEEETTTEEEEEEEEEEEE

Alr27 Domain1
 |A-|      A'     |--B---|     |--C---|   |-C'-|         |-C"-|  |---D----|   |---E---|        |----F----|    |-----G-----|
QIDVSDANNKNEASKGKDYIVHWNISNAEKIFLLELSKNKTNIKSNNPGNWKSLELFTVNKTSISIETFMEGDKMCCIMTVHNVSYDDDASEYVMQVTYILYNPIREEKKNATYRLDVT
 EEEETTT  EETTTb EEEEEEEETGGGEEEEEEEEETTEEEEEE GGGTTTEEEEEETTEEEEEEEEEETTTEEEEEEEEEETb GGGTTEEEEEEEEEEEETTTTTEEEEEEEEEEEEE

Alr28 Domain 1
          |-A'|    |-B--|       |---C--|  |--C'--|     |C"|      |-D--|    |-E--|        |----F----|  |-----G------|
YFKAECAEQNIIGKAEKGNNVTIYWSLNVTKRTYTKATLRFDNIVVSTCNAAGGNGFTCVTPKNDKYEASFNKTDKQITLTILNLTYSDSGDYLVALDYKHGKQTRPERLTNTTIQVR
          EEEEETTTb EEEEEE  TTTT EEEEEEEETTEEEEEEEE TTTTEEEE  TTTTTEEEEEEEEGGG EEEEEETTb GGG EEEEEEEEEEETTEEEEEEEEEEEEE

Alr29 Domain 1
   |A-|    |A'|   |--B---|    |--C---|  |-C'-|    C"           |--D-|   |-E--|         |---F---|    |----G----|
IGQHIAVNKTNIKATYNRNISITFFLTSTNPIKNIEFSWEEIIIATYQSSFEKVEPNYFKDRLSFSSEKSRNFKLHIRNTQYTDAGIFRFKVLFKIPDPHSATASTLMQVK
   EEEETTTEEEETTTb EEEEEEEE   EEEEEEEETTEEEEEETTTEEETTTTTTTTTEEEEEEETTTEEEEEEETTb GGG EEEEEEEETTTTEEEEEEEEEEEE

Alr30 Domain 1
 A   |A'|      |--B--|       |--C--|  |--C'-|   C"           |--D-|    |--E--|        |---F---|    |---G----|
SLISPAIIEKVVGRSVTITYVTDVADNIDAIFKIYYNDSQIGEGTKRLFETPPTPFGVRLRTSTSLQNQKKYTLHIDNLKYNDTGLFFAKIEIFKPNVEANSTTYLIIY
 EE TTEEEETTTb EEEEEEE  TTTT EEEEEEEETTEEEEEEEEEEEE   TTTGGGEEEEEETTTTEEEEEEEETb GGG EEEEEEEEETTTTEEEEEEEEEEE

Alr31 Domain 1
  |A|      |-B--|       |---C--|  |--C'-|    C"          |--D-|   |--E-|         |---F---|            |----G----|
APQVNVIAGEDLNITYYFNDEVKNWYNLRIFADNVLLFEKEKYGTTVLHNTSYGDRLVITFQEYVQALLWINNMTYTDSKIMTFEYSSSPENSSIGHIKTTNTRFPVDVK
   EEEETTb EEEEEE TTTTTTEEEEEEEEEETTEEEEEETTT EEE  TTTTTTEEEEEETTTEEEEEEETTbTTTTTEEEEEEEEEE TTTTTTT  EEEEEEEEEEE

Alr33 Domain1
 |-A-|     |-A'-|      |---B---|    |-C--|  |--C'-|  |C"|          |--D--|   |---E--|         |---F---|        |----G----|
LSQSINLANITKEITITAVHGEELKLSILMPLHSQRLLVYKLRRPIVIFTKSQLKFLDKKSFENGLTTFSESESNVIKYGLWAPKLSLNDQQRLTIVAVNNTMPDDWVTNITLNIKVY
 EEEEEETTTT  EEEEEETTTb EEEEEEEETTT EEEEEETTEEEEEEETTEEEETTTTTTTTTT EEEEEEETTEEEEEEEEEETb GGG EEEEEEEEE TTTTTEEEEEEEEEEE

Alr34 Domain 1
 |A-|   |A'|     |---B--|     |---C----|      |C'-|     C"           |-D--|    |-E--|         |----F----|   |-----G----|
GEIIIKSQITGVVRESVDISFIIDTFNGESLLSVKVFQLPDKLNPVAVGSSSAFTVAKGGSFGGRIQANVEPAKGRYTLAISTLKYVDEAAYEITATFYESTAEVRVLRKTIDLKVQ
 EEEETTTEEEETTTb EEEEEEEE TTTEEEEEEEEEEETTTTTTEEEEEETTTTEEETTTT TTTTEEEEEEGGG EEEEEETTb GGG EEEEEEEEEETTT EEEEEEEEEEEE

Alr35 Domain 1
 |A-|   |A'|     |--B---|     |---C----|      |--C'--|  |C"|          |-D--|    |-E--|         |---F---|      |----G-----|
GTVSVANKNMNVNKDNPVSLSYFIQYPPEEKFNVIDIYLKDPVSSVETILAQGLSNSLKLQGSFGGKYSTSVDVSAKKYTLTINNAQYTDSASYKAIAVFLTKTGELSVKNVEIALEVN
 EEEETTTEEEETTTb EEEEEEEE TTTTEEEEEEEEETTTTT EEEEEEEEETTEEEE   TTTTEEEEEGGG EEEEEETTb GGG EEEEEEEEEETTTT EEEEEEEEEEE

Alr36 Domain 1
 |A-|   |A'|     |--B---|      |--C--|        |C'     |C"|          |-D-|   |-E--|          |---F---|     |----G----|
GNVTAISNSMTAKLNSTVVLQWKLFIFPRMDEIRLYAEPNLVEPLIKVRYYRRGVVEILNASIEMFGNRISATYNGSLFTVLLNKVVYTDSHKYLLRVMKYYFPMRLQSTVTLNVT
 EEEETTTEEEETTTb EEEEEEEETTTT  EEEEEEETTTTTEEEEEEGGGTTEEEE HHHHHHHTTTEEEEEEETTEEEEEEETTb GGG EEEEEEEEETTTTEEEEEEEEEEE

Alr38 Domain 1
      |A-|   |A'|     |---B--|      |---C----|   |-C'-|    C"          |-D--|    |-E--|          |---F---|    |----G-----|
AEVRRAKLNVVQQDLIGLKGEDVEVTLITTMLVGQRLLSIKVFQLPGNIELATGSERAFLLQNETLRGIKFETKIDGEKGIYRFGIKSLDYDDGTMYLATAIFHDGYTRYHKDDVRVKVNVL
      EEEETTTEEEETTTb EEEEEEEE TTTTEEEEEEEEEETTTTEEEEEETTTEEEE  GGGTTTTTEEEEEEGGG EEEEEETTb GGG EEEEEEEE  EEEEEEEEEEE
```

**Fig. S8. STRIDE secondary structure predictions for Alr domain 1**
For each domain, the top line shows beta-strands labeled according to their position in the primary amino acid sequence. The middle line shows the sequence of the domain. The bottom line shows the STRIDE secondary structure predicted from the Colabfold model. (H = alpha helix, G = 3-10 helix, I = PI-helix, E = beta-strand extended conformation, B = isolated bridge, T = turn.)

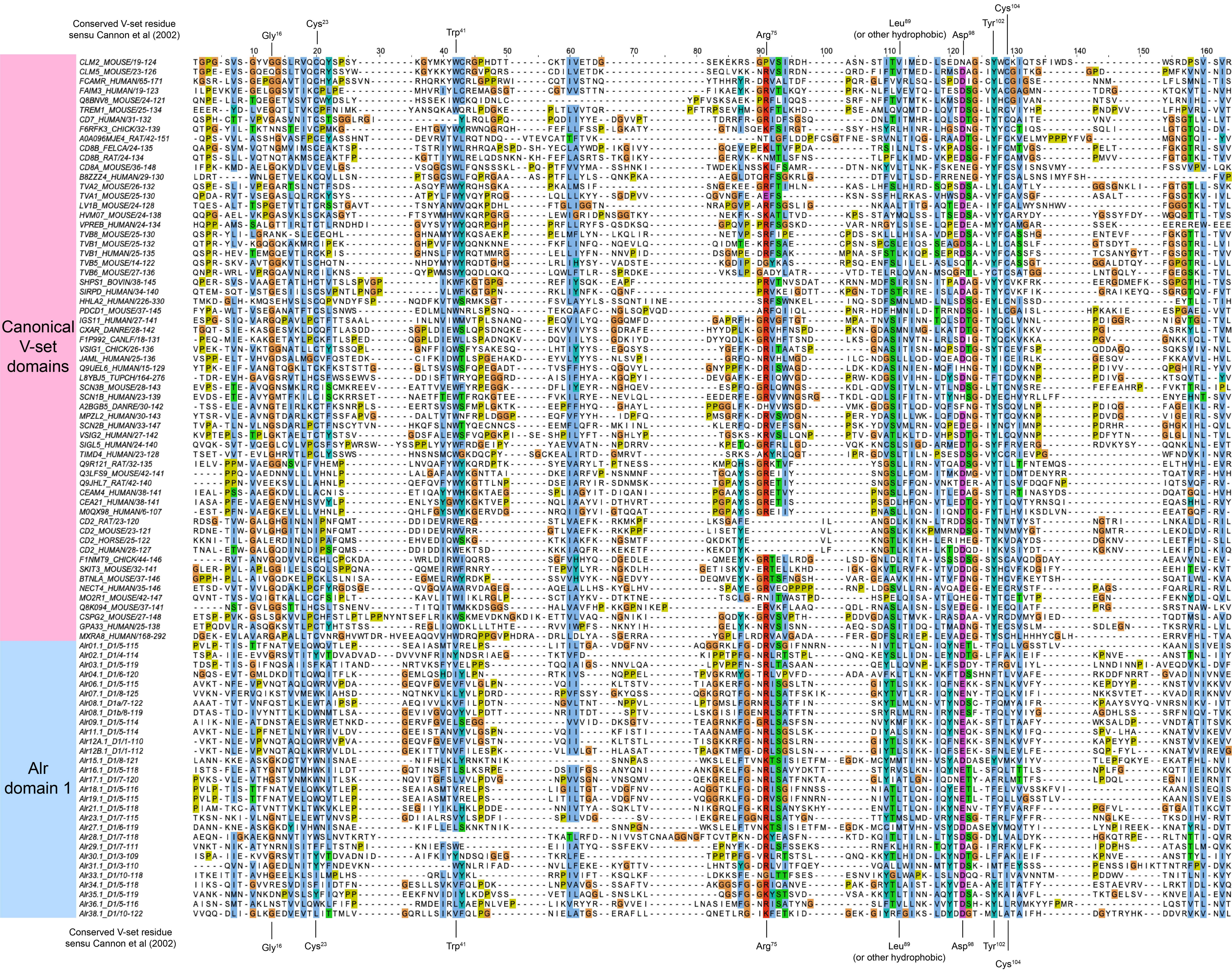

Conserved V-set residue sensu Cannon et al (2002)
Gly16
Cys23
Trp41
Arg75
Leu89 (or other hydrophobic)
Asp98
Tyr102
Cys104
Canonical V-set domains
Alr domain 1

**Fig. S9. Multiple sequence alignment of V-set Ig domains and Alr domain 1.**
Alr domain 1 sequences V-set Ig domains from pfam (pf07686). The positions of conserved V-set residues according to the nomenclature of Cannon et al [30] are shown above and below the alignment. Residues are highlighted by sequence conservation and chemical property with CLUSTALX colors as implemented in Jalview.

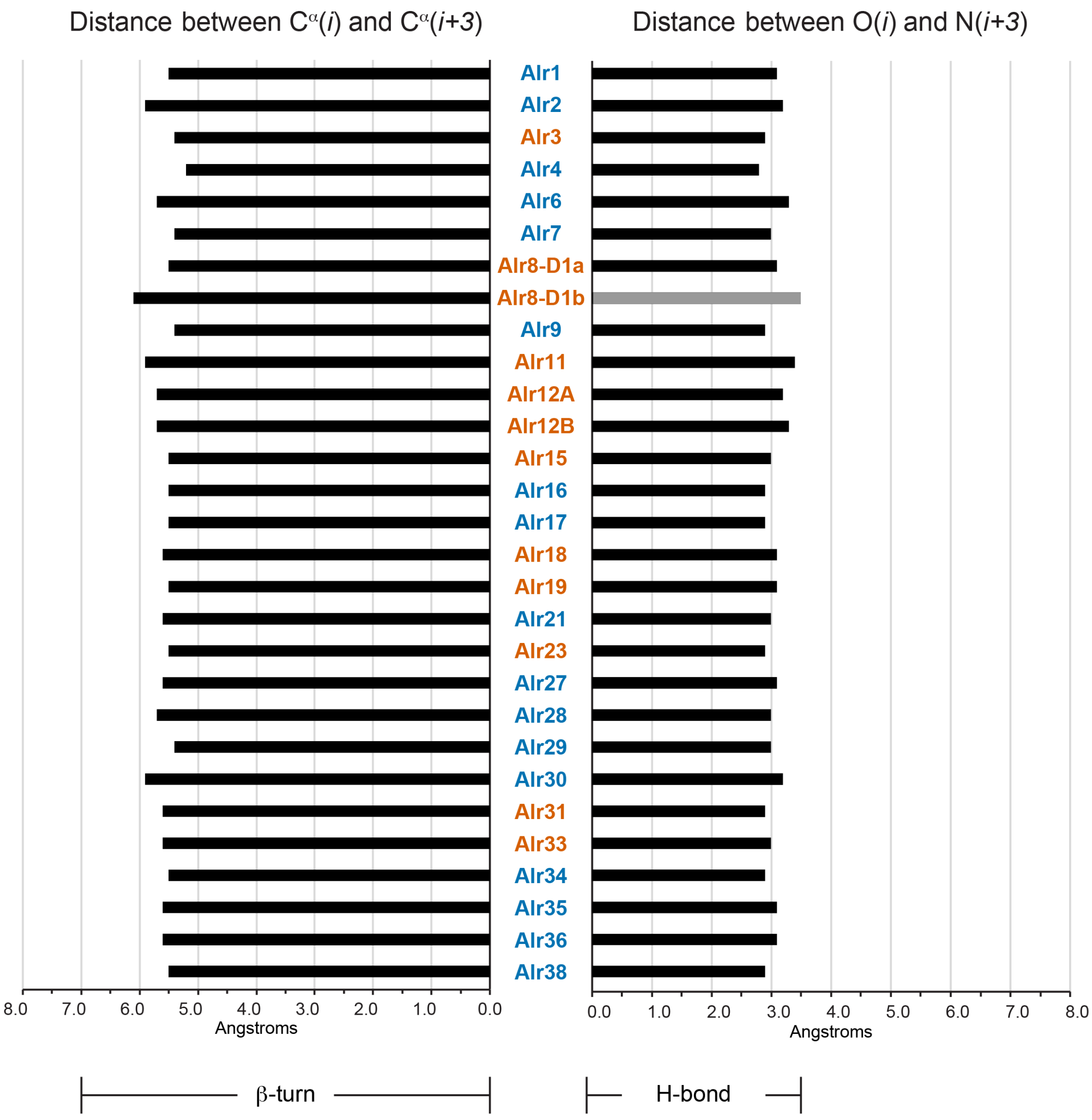


**Fig. S10. Measurements of the turn from strands A' to B in the predicted structures of domain 1.**
Left graph shows distance between the alpha carbons of the first and fourth residues in the turn. In a beta-turn, the distance between these two atoms is less than 7.0 Å. Turns <7.0 Å are shown in black bars. Those >7.0Å are showing with gray. Right graph shows distance between the oxygen atom of the first residue and the nitrogen atom of the fourth residue. A distance of <3.5Å is considered small enough for hydrogen bonding to occur. Black bars indicate distances <3.5Å. Gray bars indicate distances >3.5Å. Gene names are color-coded according ot whether they are *bona fide* (blue) or putative (orange) genes.

```
Alr1 Domain 2
   |A-|     |A′|      |--B--|     |--C-|  |-C′| |--D-|    |--E--|          |---F---|  |----G----|
GSPRVCGRKLKSNYTVNEGDFRNITQDICGYPKPKVSWTLGQENAGSSTSFAVNNATRQYEYHYKTRPFNRSDCGSNIAFIAKNTLGSINGNAWIDVD
   EEEEb    EEEETTTb  EEEEEEETTTTTEEEEEETTEEEEE EEEEEEGGG EEEEEEE   b GGGTTEEEEEEEEEETTEEEEEEEEEEE

Alr2 Domain 2
   |A-|    |A′|   |----B----|     |-C|       |--D--|   |----E----|     |---F--|    |---G----|
GGPSIISNLSSTYTVEENTTNYKIPIILQGHPKPQVTWEFVGNKNVRVETIDEKKRKYRYILTIPMITREMNGKVLSYEAVGNVSKITETAKLNVTY
   EEEET TTEEEETTEEEEEEEEEEETTTTTEEEETTTTTTEEEEEEEGGG EEEEEEEEEEE GGGTTEEEEEEEEETTTTEEEEEEEEEE

Alr2 Domain 3
  A       |A′|      |--B--|     |--C-|  C′  |--D--|  |--E--|           |---F---|  |----G----|
SPTFCDTAVKVIEFEEVASAMFTTHVCGNPMPTVSMEFGGQSVAVTHNTAEKDKYKYTANLTSYLKPSRCGRELIATAENSIGEKTSKVVLKYK
  EETTT   EEEETTTT  EEEEEEETTTT EEEEEETTEEE EEEEEEETTEEEEEEE GGG  GGGTTEEEEEEEEEETTEEEEEEEEEEE

Alr3 Domain 2
   |A-|     |A′|     |---B---|    |--C-|  C′  |--D--|      |---E---|         |---F--|    |----G---|
GGPHVCGDDLPSNITLNATFSVVLSIHLCGRQKPLVTWSIDNETVTTHNSNTTENTTADAQYVYSVVLPKVSASMCGKTLKYIATGYESDKIGTALLDTQ
   EEEEb  TTEEEETTT  EEEEEEEEETTT EEEEEETTEE  EEEEEEE  TTTTEEEEEEEEETTT GGGTTEEEEEEEETTTTEEEEEEEEEE

Alr4 Domain 2
   A       |A′|     |--B---|     |--C-|      C′  |--D-|    |---E--|         |---F---|  |----G----|
GGPKLCGDNITAITREEGSMLVAIQDVCGKPTPLVTWKFASEKNFKNSTSSGLINKEIQLYRYTYMHQNLSREQCWEPIIFKAISRLGSVSGRAMVNVT
   EE      EEEETTTb EEEEEEEETTTT EEEEEETTT   EETTEEEEEEEGGG EEEEEEEE  b GGGTTEEEEEEEEEETTEEEEEEEEEEE

Alr6 Domain 2
   A        |A′|   |----B---|     |--C-|      C′  |--D-|    |----E----|      |---F--|     |---G---|
GGPLFCGANISKSYNVTEGAALTLKQDVCGNPKPNVQWKMSSDSAYMPTKSSVLIENITRTYQYTFTTKALTRTNCGHIIEFRASGHKPDITGQTMININ
   EE      TTEEEETTTEEEEEEEEEEETTTT EEEEEETTT   EE  EEEEEETTTTEEEEEEEEEEE GGGTTEEEEEEEE     EEEEEEEEE

Alr7 Domain 2
   A        |A′|   |----B---|     |--C-|  |-C′|  |--D-|    |----E----|      |---F---|  |----G----|
GGPDVCGKGMESTYTAKEGQSLSIVQDICGHPKPVVQWKLNKDMFVSSLKSTLINDTIKQFRYSYGSRSLRRSDCGKYITFNASNDVAAIEQNAMIDVV
   EE      TTEEEETTTEEEEEEEEEEETTTTTEEEEEETTEEEEE  EEEEEETTTTEEEEEEEEEEE GGGTTEEEEEEEEEETTEEEEEEEEEEE

Alr8 Domain 2a
   A        |A′|   |----B---|     |--C--|       |---D--|    |----E----|      |---F---|  |----G----|
GVPRVCGKGLASKIIIAEKESLTITQDICGHPKPDFKWKFKGDKIWRILLCSTILDDATKQYRHEFKTRPLTRADCGKTIIFNASNELGSLAGETSVDVM
   EE      TTEEEETTTEEEEEEEEEEETTTTTEEEEEEEETT TTTTEEEEEEEEEGGG EEEEEEEEEEE GGGTTEEEEEEEEEEETTEEEEEEEEEEE

Alr8 Domain 2b
   A        |A′|      |--B--|     |--C-|   |-C′-| |--D-|    |--E--|          |---F---|  |----G----|
GTPKICGVRLKSNYTVVENSTITFTQDVCAHPKPVAEWKLDQEKLYKKPSNTSLINTEQRKYRFTYQTRKLTRNDCGAKFILKATNAMGSVEETVKVDVI
   EE      TTEEEETTTb  EEEEEEETTTT EEEEEETTTEEEEEE EEEEEEGGG EEEEEEE   b GGGTTEEEEEEEEEETTEEEEEEEEEEE

Alr9 Domain 2
            |A′|   |---B----|     |--C-|                    |-----E----|      |---F---|  |----G----|
GGPDQCGISLNSSYTVHEGKQLSLLSEICGNPKPILTWKLQNELGYSYSSDFMLMDIFSMRYRYVYKTRRLVTREDCGTKLAFNATGASGTIQGYAILDVT
  TTT   B TTEEEETTTEEEEEEEEEEETTTT EEEEEETTB     BTTB   TTTTTEEEEEEEEEEEEE GGGTTEEEEEEEEEETTEEEEEEEEEEE

Alr11 Domain 2
   |A-|     |A′|      |--B--|     |--C-|      C′  |--D-|    |--E--|           |---F---|  |----G----|
GGPDVCGISLKSSYIVNEGKKLSFLSEVCGNPKPILTWKMENELEYSYSADIRYMDKSTMRYQYVYKTRSHITRKDCGTKIIFNATGANKMITGEAVISVT
   EEEEb  TTEEEETTTb  EEEEEEETTTT EEEEEETTT   EETTEEEEEEEGGG EEEEEEE    b GGGTTEEEEEEEEEETTEEEEEEEEEEE

Alr12A Domain 2
   A        |A′|   |----B----|    |--C-|  C′  |----D---|  |------E-----|      |---F---|  |----G----|
GGPEACGITLNTSYAVNEGKQLSVTTEVCGNPKPVLTWQLQEELEYSYSTEVAPVNISIMRYRYVYKTRRLVTREDCETKLVFNATGANGMIKEETLIDVT
   EE      TTEEEETTTEEEEEEEEEEEETTT EEEEEETTEEE EEEEEEEEEETTEEEEEEEEEEEEEE GGGTTEEEEEEEEEETTEEEEEEEEEEE

Alr12B Domain 2
            |A′|   |----B----|    |--C-|  |C′| |---D---|  |-----E------|      |---F---|  |----G----|
GGPEACGITLNTSYAVNEGKQLSVTTEVCGNPKPVLTWQLQEELEYSYSTEVAPVNISIMRYRYVYKTRRLVTREDCGTKLVFNATGANGMIKGETLIDVT
   TTT  B TTEEEETTTEEEEEEEEEEEETTT EEEEEETTEEEE EEEEEEEEETTEEEEEEEEEEEEEE GGGTTEEEEEEEEEETTEEEEEEEEEEE

Alr15 Domain 2
   |A|      A′     |---B---|   |--C-|   C′  |--D--|    |--E--|       |---F--|     |---G---|
GGPEFCGMKLPNQVNVNNEKAEVEICGHPKPEVNFYIDEGKRLPGACKLTDERLKKYKCEVELTDLNCGEMLYLEGRGYDTMKLTSSSLLIAN
   EEEb   TTEEEGGG EEEEEEEEETTTEEEEEEETTTEEE EEEEEEEGGG EEEEEEE  GGGTTEEEEEEEE TTTTEEEEEEEEE

Alr16 Domain 2
   A       A′       |---B---|     |--C-|     C′   |--D-|    |---E---|        |---F---|  |---G---|
GSPRICGKSLKSIYTASDTAVLTVTQDICGHPKPFVKWKLERDNTFSDSFSSVLMSNTSRKYRYSFATRNIIRSDCGEKIMFNARNKFGNFNGSLTIYIS
   EE      EEE  TTTTEEEEEEEEEETTTTTEEEEEETTTT EEE  EEEEEETTTTEEEEEEEEE   GGGTTEEEEEEEEEETTEEEEEEEEE

Alr18 Domain 2
   |A-|     |A′|   |----B---|     |--C-|         |--D-|    |----E----|     |---F---|  |----G----|
GSPRFCGPKVEFSYSVTEGSILTILQDICGNPKPDVKWKVGEDIFTSSSSSSVINNATRKYEYTFTTRPLNRNDCGINITVFASNTLGYFERNTMVHVE
   EEEEb  TTEEEETTTEEEEEEEEEEETTTTTEEEEEEETTB      EEEEEEGGG EEEEEEEEEEE GGGTTEEEEEEEEEETTEEEEEEEEEEE

Alr19 Domain 2
   A        |A′|   |----B---|     |--C-|     C'  |--D-|    |----E----|     |---F---|  |----G----|
GSPSLCGRNLESIYTVNQDNILNVVQFICGYPNPEVKWKVGDDNFSKSYSLSEINTAMRQYKYTFRTRPITRNDCGQTLIFVANNTVGSIQRNAELDVE
   EE      TTEEEETTTEEEEEEEEEEETTTTTEEEEEEETTT  EE  EEEEEEGGG EEEEEEEEEEE GGGTTEEEEEEEEEETTEEEEEEEEEEE

Alr21 Domain 2
   A        |A′|   |----B---|     |--C-|      |---D---|    |----E----|     |---F---|  |----G----|
GPPNVCGKSVKSRYYITDTATITVAQDICGHPKPFVEWKLEDTSFSSSSRFMLIDDATKKYRYSFTTENIVRSNCGKKIMYHARNEFGNVEGYSMIYIS
   EE      TTEEEETTTEEEEEEEEEEETTTTTEEEEEEETTT   EEEEEEEEEGGG EEEEEEEEEEE GGGTTEEEEEEEEEETTEEEEEEEEEEE

Alr23 Domain 2
   |A-|     |A′|   |----B---|     |--C-|     C′    |D-|    |----E----|     |---F---|  |----G----|
GSSIFCDISSKYHHTVNESEVLTVVQNVCGYPIPELKWKVGDDNFSTASSPSVINNETRQYEYSFKTRHITRKDCGRNLTLVTSNTIGSMEINAKIDVK
   EEEETTTT EEEETTEEEEEEEEEEEETTTTTEEEEEEETTT  EE    EEEEGGG EEEEEEEEEEE GGGTTEEEEEEEEEETTEEEEEEEEEEE

Alr27 Domain 2
   |A |     A′     |--B--|     |--C-|   C′  |--D--|    |--E--|       |---F--|     |---G---|
GVPEFCGTKLPNEINLINKNAEVEICGHPKPEVNFYIHESKRLPGACELTEGRLKKYKCKVELTDLKCGEKLNLKIRGYQNMKLSSSSILIAN
   EEEEb  TTEEEGGGTEEEEEEETTTTTEEEEEEETTTEEE EEEEEEETTTTEEEEEEETTTTTTEEEEEEEE GGG EEEEEEEEE

Alr28 Domain 2
   |A-|     |A′|      |--B--|     |--C-|      |---D--|  |--E--|          |---F---|  |----G----|
GPPQICDSNFMTSYEFAQHYQPIVNITICGYPTPRFSWAFGQNTKPDISIRSIKAHKHVFSAVLSNLTSSMCGSKLTFKASNKFGKVSTSAVINVS
   EEEE   TTEEEETTTb  EEEEEEETTTTEEEEEEETTTB  EEEEEEEETTEEEEEEEETTTb GGGTTEEEEEEEEEETTEEEEEEEEEEE

Alr29 Domain 2
   A        A′       |--B--|     |--C-|  C′  |--D--|    |--E--|        |---F--|      |---G--|
GGPDICGKQLPLQISFRNTSSRIISTNVCGNPPPQIFWSMDGMKLQGTRKERDQSRKKFEYSVNLTGFDLCDKELSYKIVGALSENTLVGSIKLT
   EE      TTEE        EEEEEEETTTT EEEEEETTEEE EEEEEEEGGG EEEEEEETTTT TTTEEEEEEEETTTT  EEEEEEEE

Alr30 Domain 2
   |A-|    |A′|   |----B----|     |--C-|      |---D--|  |----E----|     |---F--|     |----G---|
GGPSIISNLSSTYTVEENTTNYNVPIILQGHPKPQVTWEFAGNKNVRMEAIAEKKRKYKYILTIPIITREMNGKVLSYKAVGYNDNEVVGSTKLNVTY
   EEEET TTEEEETTEEEEEEEEEEEETTTT EEEEEETTTTTEEEEEEEEETTEEEEEEEEEEE GGGTTEEEEEEEEETTTTEEEEEEEEEE

Alr30 Domain 3
  A        |A′|      |--B--|     |--C-|  C′  |--D-|  |--E--|           |---F---|  |----G----|
SPTFCDTAVIEVIEFAEVASAVFTTHVCGNPMPAVYMEFDGEPLATVLDAEKDKYKYTVNLTPYLLPSRCGKKLIITAKNNKGEGTRRIVLKYK
  EE TTTT  EEEETTTT  EEEEEEETTTT EEEEEETTEEE EEEEEETTEEEEEEE GGGTTGGGTTEEEEEEEEEETTEEEEEEEEEEE

Alr31 Domain 2
   A        |A′|      |--B--|     |--C-|  C′   |--D-|    |--E--|          |---F---|     |----G---|
GGPSICGALPSPVTYLKSRDQTTLSVFLCGHPTPEVIWKVDNRFLNGTVKEIDEETELFKYTLDFKNYLQPKKCSVKVSLMARGHSEEEQVFQSEIQYG
   EE      TTEEEETTTTTTEEEEEEEETTTT EEEEEETTEEE  EEEEEETTTTEEEEEEE GGG TTTT EEEEEEEEE     EEEEEEEEEE

Alr33 Domain 2
   A              |---B--|     |--C-|  C′    |--D-|  |---E--|         |---F--|     |---G--|
GGPFVCNKERKIVIRTNPLRVKDIICSNPKPMVTWYVDNNLISGGDVLKKQLTKYSVRKIFTTQDVSLCGKRISYVATGRNGISLNRSTLLVFP
   EETTT      TTTEEEEEEEEETTTT EEEEEETTEEE   EEEEEETTEEEEEEEE GGGGGGTTEEEEEEEEETTTTEEEEEEEE

Alr34 Domain 2
   |A-|     |A′|   |----B---|     |--C-|  C′   |-D-|    |----E---|     |---F---|  |----G----|
GGPDNCGERLSPITLVEEEQASVFVAFLCGNPKPTVHWTIGRRQISSYVDNTPSTGMYKYSTNIKITLETCGETVRYVAAGYGRELTSSTVIKVS
   EEEEb  TTEEEETTTEEEEEEEEEEETTTT EEEEEETTEEE  EEEEETTT EEEEEEEEEE GGGTTEEEEEEEEEETTEEEEEEEEEEE

Alr35 Domain 2
   A        |A′|      |--B--|     |--C-|  C′  |-D-|     |--E--|         |---F---|  |----G----|
GGPEFCNLKPPETLTVISHERPRVRVSVCGNPEPQITWEYLGNNLKSTIIKGKKPKEFIFETLLPQITPPNCGSSLTYKATGHGRGVSDRTTLNVK
   EETTT  TTEEEETTTb  EEEEEEETTTTEEEEEEETTEEE EEEEE  TTTEEEEEEE   b GGGTTEEEEEEEEEETTEEEEEEEEEEE

Alr36 Domain 2
   |A-|     |A′|     |---B--|     |--C-|         |--D--|   |---E--|         |---F---|  |----G----|
GGPDDCGDSLKSMYNVEEFEKLLVTKEICGHPKPTVRWKFQRNRFYSPSASSQLINNSTRLYRYSFEWRNITRANCGRKLLFSASGYGESLSENAWLNVV
   EEEEb  TTEEEETTTb EEEEEEEETTTTTEEEEEEETTT  TTTEEEEEEEETTTEEEEEEEEE  b GGGTTEEEEEEEEEETTEEEEEEEEEEE

Alr37 Domain 2
     |----A-----|  |---B---|     |--C-|  |-C′|  |--D-| |---E--|       |---F--|     |---G--|
IFEKPYRCGHRTLQQISSSSFQVKETICGQRIPSLVWYLDGKLIGNGTAREIEKHKYVFTKTFKDMTACGRNLSYVITSFDGYTATYYATLSFN
     EEEEEEEEEEETTEEEEEEEEEETTTT EEEEEETTEEEE EEEEEETTEEEEEEEE TTTTTTEEEEEEEETTTT EEEEEEEE

Alr38 Domain 2
   |A-|     |A′|   |----B---|     |--C-|  C′   |D-|     |----E---|     |---F---|  |----G----|
GGPQLCGSTLPAVTMVKRNTTVRFSSVVCGNRPPSMSWMLGCESLKTNVTKGLKPQEYVHHVDLKITTRMCGLLLRYVASGHDGEITGSTKLIEK
   EEEEb  TTEEEETTTEEEEEEEEEEETTTT EEEEEETTEEE  EEEE  TTTEEEEEEEEEETTTTTEEEEEEEEEETTEEEEEEEEEEE
```

**Fig. S11. STRIDE secondary structure predictions for AIr domains 2 and 3**
For each domain, the top line shows beta-strands labeled according to their position in the primary amino acid sequence. The middle line shows the sequence of the domain. The bottom line shows the STRIDE secondary structure predicted from the Colabfold model. (H = alpha helix, G = 3-10 helix, I = PI-helix, E = beta-strand extended conformation, B = isolated bridge, T = turn.)
Figure S9 – I-set alignment

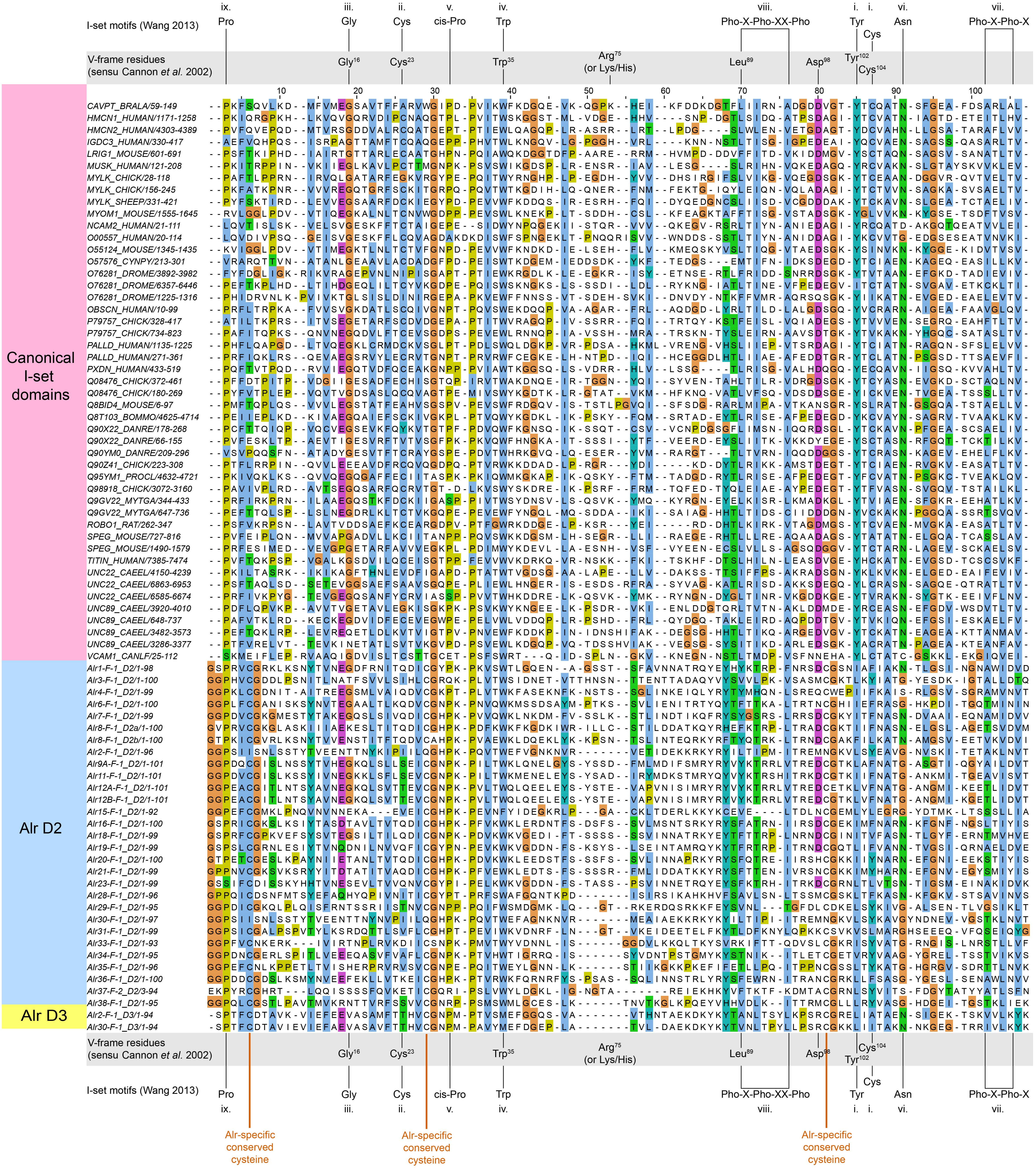

I-set motifs (Wang 2013)
ix. Pro
iii. Gly
ii. Cys
v. cis-Pro
iv. Trp
viii. Pho-X-Pho-XX-Pho
i. Tyr
i. Cys
vi. Asn
vii. Pho-X-Pho-X
V-frame residues (sensu Cannon et al. 2002)
Gly16
Cys23
Trp35
Arg75 (or Lys/His)
Leu89
Asp98
Tyr102
Cys104
Canonical I-set domains
Alr D2
Alr D3
CAVPT_BRALA/59-149
HMCN1_HUMAN/1171-1258
HMCN2_HUMAN/4303-4389
IGDC3_HUMAN/330-417
LRIG1_MOUSE/601-691
MUSK_HUMAN/121-208
MYLK_CHICK/28-118
MYLK_CHICK/156-245
MYLK_SHEEP/331-421
MYOM1_MOUSE/1555-1645
NCAM2_HUMAN/21-111
O00557_HUMAN/20-114
O55124_MOUSE/1345-1435
O57576_CYNPY/213-301
O76281_DROME/3892-3982
O76281_DROME/6357-6446
O76281_DROME/1225-1316
OBSCN_HUMAN/10-99
P79757_CHICK/328-417
P79757_CHICK/734-823
PALLD_HUMAN/1135-1225
PALLD_HUMAN/271-361
PXDN_HUMAN/433-519
Q08476_CHICK/372-461
Q08476_CHICK/180-269
Q8BID4_MOUSE/6-97
Q8T103_BOMMO/4625-4714
Q90X22_DANRE/178-268
Q90X22_DANRE/66-155
Q90YM0_DANRE/209-296
Q90Z41_CHICK/223-308
Q95YM1_PROCL/4632-4721
Q98918_CHICK/3072-3160
Q9GV22_MYTGA/344-433
Q9GV22_MYTGA/647-736
ROBO1_RAT/262-347
SPEG_MOUSE/727-816
SPEG_MOUSE/1490-1579
TITIN_HUMAN/7385-7474
UNC22_CAEEL/4150-4239
UNC22_CAEEL/6863-6953
UNC22_CAEEL/6585-6674
UNC89_CAEEL/3920-4010
UNC89_CAEEL/648-737
UNC89_CAEEL/3482-3573
UNC89_CAEEL/3286-3377
VCAM1_CANLF/25-112
Alr1-F-1_D2/1-98
Alr3-F-1_D2/1-100
Alr4-F-1_D2/1-99
Alr6-F-1_D2/1-100
Alr7-F-1_D2/1-99
Alr8-F-1_D2a/1-100
Alr8-F-1_D2b/1-100
Alr2-F-1_D2/1-96
Alr9A-F-1_D2/1-101
Alr11-F-1_D2/1-101
Alr12A-F-1_D2/1-101
Alr12B-F-1_D2/1-101
Alr15-F-1_D2/1-92
Alr16-F-1_D2/1-100
Alr18-F-1_D2/1-99
Alr19-F-1_D2/1-99
Alr20-F-1_D2/1-100
Alr21-F-1_D2/1-99
Alr23-F-1_D2/1-99
Alr28-F-1_D2/1-96
Alr29-F-1_D2/1-95
Alr30-F-1_D2/1-97
Alr31-F-1_D2/1-99
Alr33-F-1_D2/1-93
Alr34-F-1_D2/1-95
Alr35-F-1_D2/1-96
Alr36-F-1_D2/1-100
Alr37-F-2_D2/3-94
Alr38-F-1_D2/1-95
Alr2-F-1_D3/1-94
Alr30-F-1_D3/1-94
Alr-specific conserved cysteine
Alr-specific conserved cysteine
Alr-specific conserved cysteine

**Fig. S12. Multiple sequence alignment of I-set Ig domains and Alr domains 2 and 3.**

Alr domain 2 and 3 sequences were aligned to I-set Ig domains from pfam. Residues in the alignment are highlighted by sequence conservation and chemical property with CLUSTALX colors as implemented in Jalview. The positions of conserved V-frame residues are shown above and below the alignment with gray background. Motifs common to I-set domains are also indicated. The position of invariant cysteine residues is shown in red lettering. Note that domain 3 is the most membrane-proximal Ig domain in Alr2 and Alr30, hence the conserved cysteine appears there and not in domain 2.

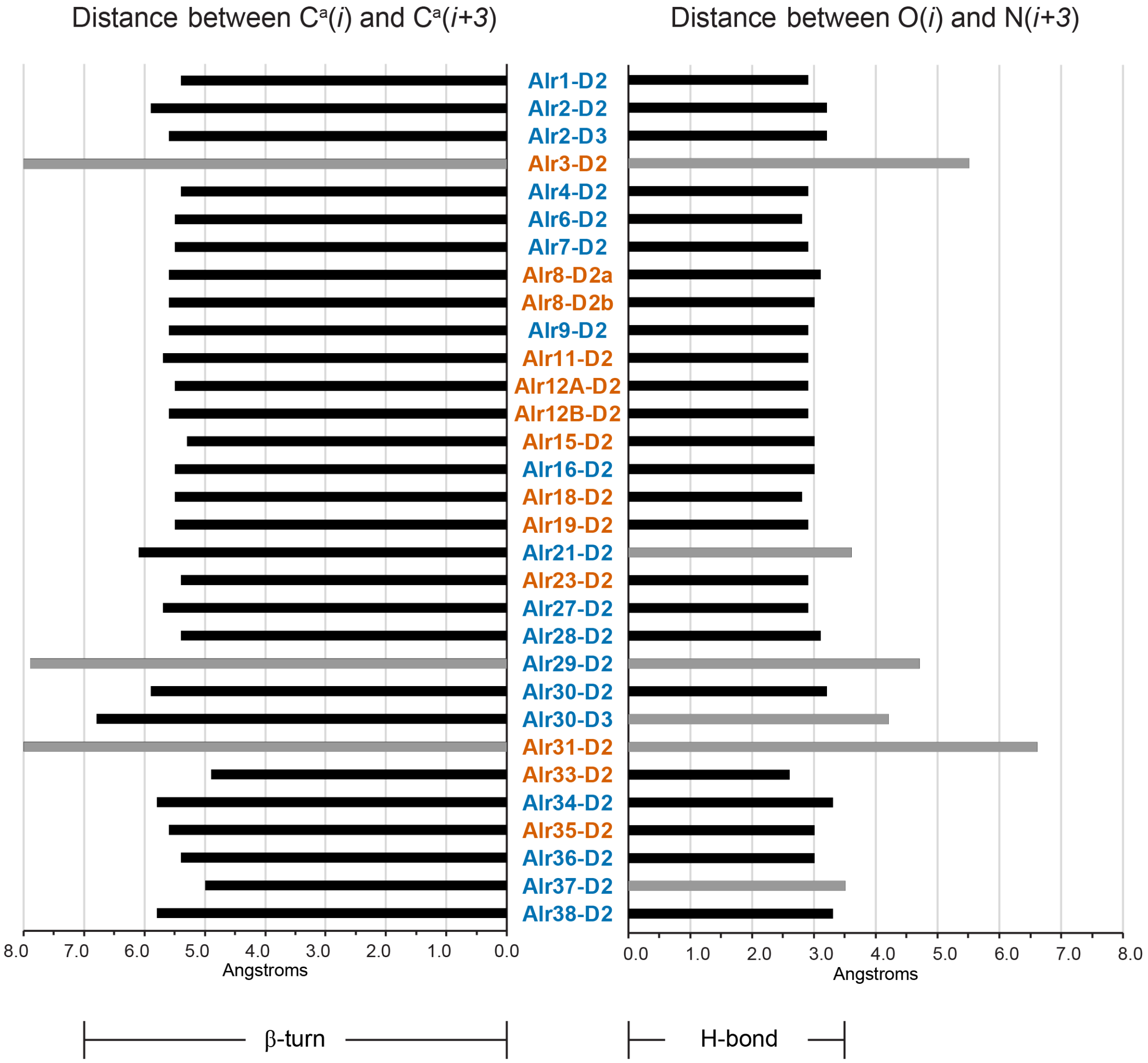


**Fig. S13. Measurements of the turn from strands A' to B in the predicted structures of domains 2 and 3.**

Left graph shows distance between the alpha carbons of the first and fourth residues in the turn. In a beta-turn, the distance between these two atoms is less than 7.0 Å. Turns <7.0 Å are shown in black bars. Those >7.0Å are showing with gray. Right graph shows distance between the oxygen atom of the first residue and the nitrogen atom of the fourth residue. A distance of <3.5Å is considered small enough for hydrogen bonding to occur. Black bars indicate distances <3.5Å. Gray bars indicate distances >3.5Å. Gene names are color-coded according ot whether they are *bona fide* (blue) or putative (orange) genes.

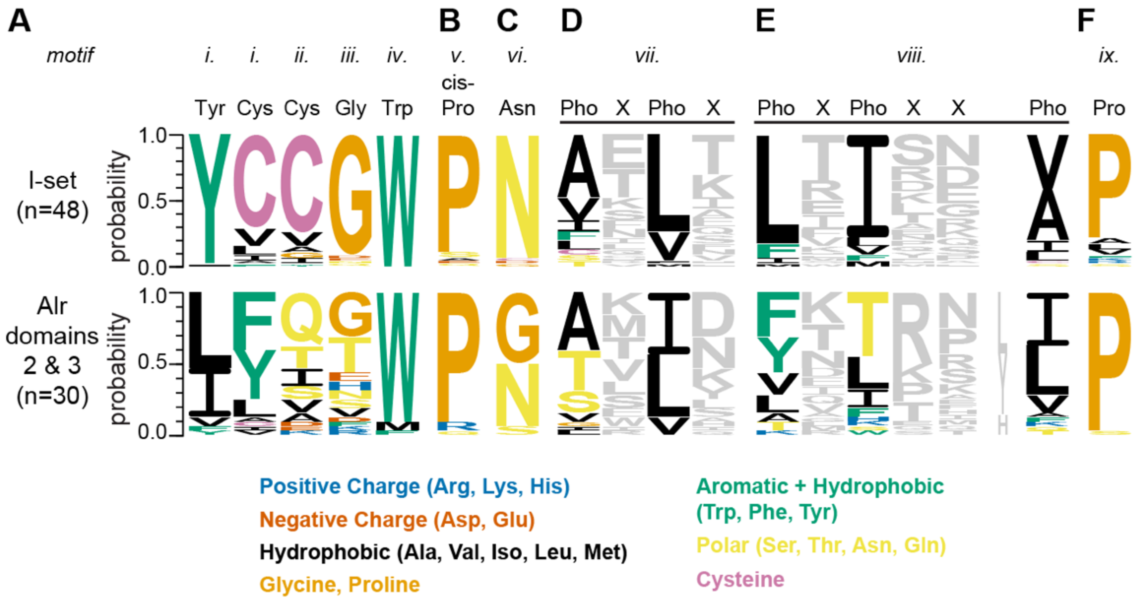


**Fig. S14. Sequence logo of I-set-specific motifs in classical I-set domains and Alr domains 2 and 3.**
Sequences of Alr domains 2 and 3 and I-set domains from the pfam I-set sequence profile (pf07679) were aligned in MAFFT. Sequence logos for then created to represent the motifs identified by Wang (2013) for the Alr or pfam sequences.

(A) Motifs *i*, *ii*, *iii*, and *iv*, which are discussed in the main text because they are also part of the V-frame as described by Cannon et al (2002).

(B) Motif *v* is a conserved proline in the BC loop, and motif *vi* is a conserved asparagine in the FG loop. These two residues form a hydrogen bond that stabilizes the BC and FG loops in a closed position. In domains 2 and 3, we found the conserved proline residue in 28/31 sequences but the asparagine was only present in 13/31 sequences (see also Figure S9). Thus, motif *v* is present in domains 2 and 3, but motif *vi* and the structural motif it forms with motif *v* do not appear to be a common feature of these domains.

(C) Motif *vii* is a Pho-X-Pho-X pattern of amino acids (where Pho represents a hydrophobic residue), located approximately 10-12 residues downstream of motif *vi*. It is found in beta-strand G and denotes the C-terminal end of an I-set (and also V-set) domain. This pattern was found in 30/31 of domain 2 and 3 sequences (See also Figure S9).

(D) Motif *viii* is a set of hydrophobic and hydrophilic residues, Pho-X-Pho-X-X-Pho, located at the bottom of beta-strand E. The last two hydrophobic residues typically contact the tyrosine in the tyrosine corner, while the two consecutive hydrophilic residues between them form a beta bulge. In our alignments, 26/31 domain 2 and 3 sequences had this motif, although the first and second hydrophobic residues often had polar or aromatic side chains (see also Figure S9). As noted above, the tyrosine corner is not present in domains 2 and 3. However, a beta bulge was predicted to occur at the end of the E strand in 28/31 structural models. Motif *viii* is therefore present in most domains, but the structural consequences of this motif are likely to differ from traditional I-set domains.

(E) Motif *viiii* is a proline ~23-26 residues upstream of the B-strand cysteine. This proline defines the beginning of an I-set domain and, in domains 2-3, it was found in 30/31 sequences (see also Figure S9).

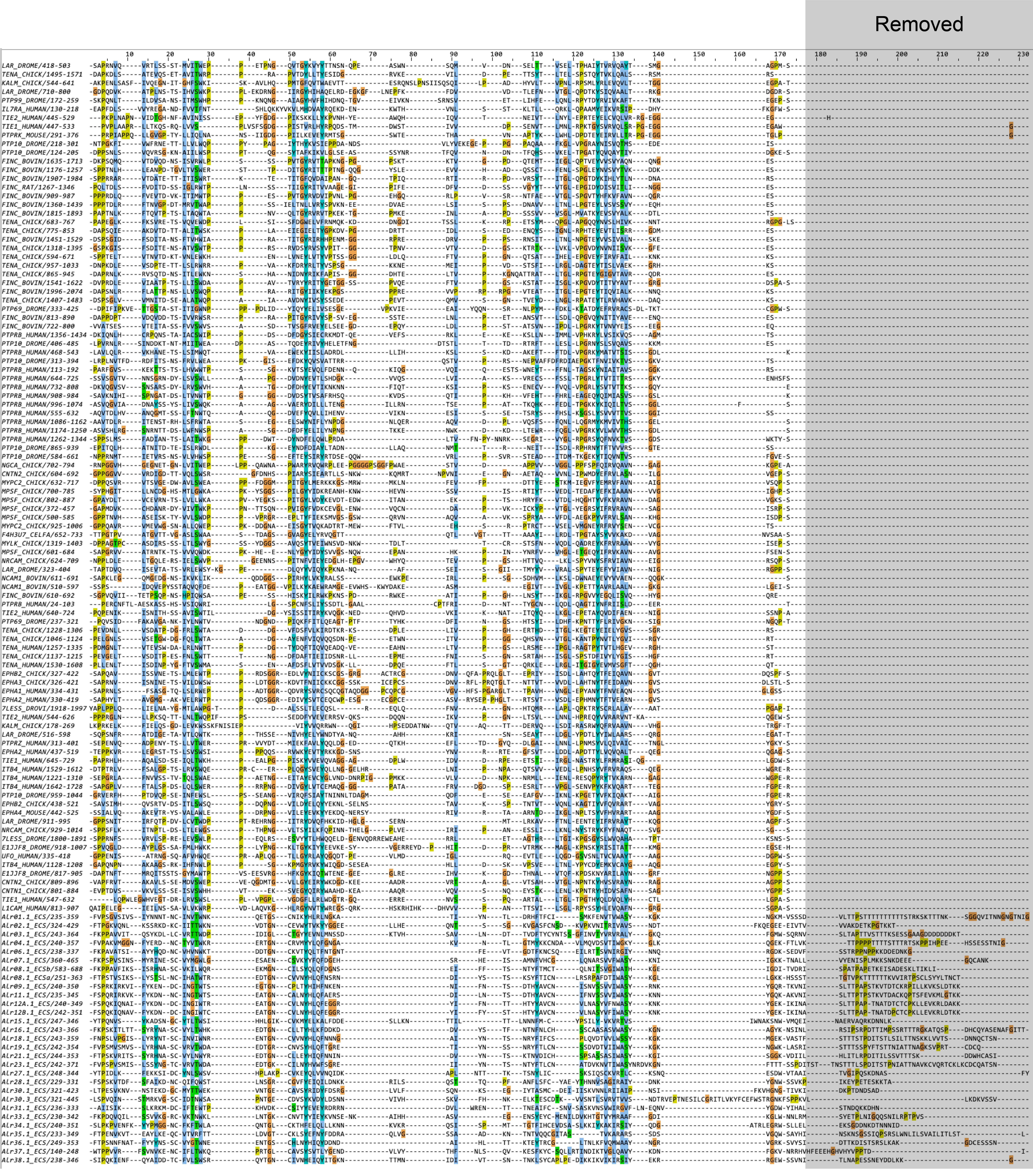


**Figure S12. Sequence removed from the ECS prior to structural prediction.**

**Fig. S15. Amino acid sequence removed from the ECS prior to structure prediction.**

```
Alr1 ECS-trimmed
      |--A---|   |--B--|         |--C-|     |C’|  |-E-|      |--F---|  |---G---|
FVPSGVSIVSIYNNNTNCINVTWNKQETGSCNIKYHLRLNGKATIYNTLDRHFTFCISMKFENVTVWASYKGKNGKMVSSSD
      EEEEEEEETTEEEEEEEE   TTTT  EEEEEETTT  EEEE  EEEEE      EEEEEEEETTEEEEEEEEE

Alr2 ECS-trimmed
      |--A---|  |--B--|        |---C---|  |--C’--|     |-E-|         |--F---|      |--G---|
FTPGKVQNLKSSRKDKCIITTWKNVDTGNCEVWYTVKYYGGEELLHEQNTSSGKVGASFCNSDKVPKVNITRIVAVSNDTFKQEGEEEIVTV
     TEEEEEEEETTEEEEEEE   TTTT EEEEEEEEETTEEEEEEEETTTT EEEEETTTTGGG  EEEEEEEE TTTT EEEEEEEE

Alr3 ECS-trimmed
      |--A---|  |--B--|         |--C---|   |--C’--|     |-E-|           |--F---|  |---G---|
FKPPAVVITQSYKDLLCVRTTWDPIDIGLCTGYYEVNLMNSSDKTVHSAVLNDTTVDFTYCYNTSSGFINVTYVRVRALYGKQFGMWSQRNV
      EEEEEEEETTEEEEEEE   TTTT  EEEEEEEETTTEEEEEEEETTTT EEEEE  GGG      EEEEEEEETTEE   EEEE

Alr4 ECS-trimmed
      |--A---|  |--B--|         |--C-|    |C’|   |-E-|         |--F---|       |G-|
FVPAKVMGGNFYERDNCTYVTWKRERTGNCRVMYYLQFGNGAKVNTLGTMYKKCNDAVLMQVDSVTIWGKYGLKQGEKFTLVK
      EEEEEEEETTEEEEEEE   TTTT  EEEEEETTT EEEE   EEEEETTGGGGG  EEEEEEEETTEE   EEEE

Alr6 ECS-trimmed
      |--A---|  |--B--|         |--C-|    |C’-|  |E-|         |--F---|  |---G---|
FKPAVATSIAYYHQDNCVHVNWKTDDTGNCKVTYQLTFNTGNTFEVTGDTFKNCSQEILRTTSVNIRGIYNNQRGDKSEDVF
      EEEEEEEETTEEEEEEE   TTTT  EEEEEETTT EEEEE  EEEE  GGGTTT EEEEEEEETTEEEEEEEEE

Alr7 ECS-trimmed
      |--A---|   |--B--|         |--C-|      C’    |-E-|      |--F---|       |G-|
FKPSPVSINSMYRINESCVYMGWLGESAENCSVKYYFQFDGEHSRHEISAMNFVHCGLQNARSVVFWASYKNIIGKKTNALL
      EEEEEEEETTEEEEEEEE   TTTT  EEEEEETTT   EEE TTEEEEE TTTT EEEEEEEETTEE   EEEE

Alr8 ECSa-trimmed
      |--A---|   |--B--|         |--C-|    |-C’|   |-E-|       |--F---|  |--G--|
FTPSTVSIKSLYSTSLNCIHATWTREDTGNCVVNYHLQFNSRNDIYSTSKSYFTICNLRSRPAFDTIWASYKGLLGKKHSSST
      EEEEEEEETTEEEEEEEE   TTTT  EEEEEETTTTEEEEE   EEEEE TTTT  EEEEEEEETTEEEEEEE

Alr8 ECSb-trimmed
      |--A---|   |--B--|         |-C--|     |C’|   |-E-|      |--F---|  |---G---|
FKPPAVFIKSISRHNASCVKILWQREKMGNCILSYQLQFDGNSEIFFTSNTYFTMCTQLNITSVGIWATHKGEIGDITVDRI
      EEEEEEEETTEEEEEEEE TTTTTT  EEEEEETTT  EEEE   EEEEE TTTT EEEEEEEETTEE   EEEE

Alr9 ECS-trimmed
      |--A---|  |--B--|         |-C--|     |C’|   |-E-|      |--F---|  |---G---|
FSPRKIRKVIFYKENDCINGTWTSEGTGNCPLTYHIHFNKENNIFNTSDTHYAVCNISNVSSVVIWASYRNKYGQRTKVNI
     TEEEEEEEETTEEEEEEE TTTTTT  EEEEEETTT  EEEE   EEEEE TTTT EEEEEEEETTEE   EEEE

Alr11 ECS-trimmed
      |--A---|  |--B--|         |--C-|     |C’|   |-E-|      |--F---|  |---G---|
FSPQRIRKVKFYKDNNCINGTWTSEATGNCALNYHLQFAERSDILNTSDTHYAVCNIFNVSSVVIWASYKNNYGQKAKVNI
     TEEEEEEEETTEEEEEEE   TTTT  EEEEEETTT  EEEE   EEEEETT    EEEEEEEETTEE   EEEE

Alr12A ECS-trimmed
      |--A---|  |--B--|         |--C-|     |C’|   |-E-|      |--F---|  |---G---|
FSPQKIQNAIFYKDNDCINGIWTREATGNCVLNYHLQFEGGRYIFNTTHTYYAVCNVLNASYVFNWASYKNKYGEKIKINA
     TEEEEEEEETTEEEEEEE   TTTT  EEEEEETTT  EEEE   EEEEETTTTT EEEEEEEETTEE   EEEE

Alr12B ECS-trimmed
      |--A---|  |--B--|         |--C-|     |C’|   |-E-|      |--F---|  |---G---|
FSPQKIQNAVFYKDNDCINGIWTCEATGNCVLNYHLQFEGGRYIFYSTHTYYAVCNVLNASYVFIWASYKNKYGEKIKINA
     TEEEEEEEETTEEEEEEE   TTTT  EEEEEETTT  EEEE   EEEEE TTTT EEEEEEEETTEE   EEEE

Alr15 ECS-trimmed
  |---A---|   |--B---|       |----C-----|  |--C’---|   |E-|    |---F----|        |G-|
YTPQNVSYKADSNGCYTLTWSIHHLGIKCVKMYELKALFDDESLLKNTTILTTLNNNFMCYPSILYVKVRTVSIWNAKSNWVMQEI
  EEEEEEEEETTTEEEEEEEETTTTGGGEEEEEEEEEEEETTEEEEEEEEE   EEEE TTTEEEEEEEEEETTT B   EEEE

Alr16 ECS-trimmed
      |--A---|   |--B--|         |--C-|     |C’|   |E-|       |--F---|  |---G---|
FKPSKITLTTSYRYNASCVYLTWHKEDTGNCLLTYHLKFDDKDDVYSTFNTNFNLCHSSCAASASVWASYKGNAGYKNSINL
      EEEEEEEETTEEEEEEEE   TTTT  EEEEEETTT  EEEE   EEEEETTTTT EEEEEEEETTEEEEEEEEE

Alr18 ECS-trimmed
         |-A-|   |-B-|           |--C-|     |C’|   |-E-|      |--F---|  |---G---|
FNPSLVPGISLYRYNTSCINVIWNRENTGNCVVNYHLQFNNNGAIYNTSNRYFIFCSPLQVDTVVLWSSYKGKMGEKVASTF
         EEEEETTTEEEEE     TTTT  EEEEEETTT  EEEE   EEEEE      EEEEEEEETTEE   EEEE

Alr19 ECS-trimmed
      |--A---|   |--B--|         |--C--|    |C’|   |E-|       |--F---|  |---G---|
FVPSMVSMVSLYRHNASCVRVTWDAEDTGRCNVSYHLQFTGRETIYNSSNRYFTLCNSSDVDTVIIWASYKGKNGWKLASRI
      EEEEEEEETTEEEEEEEE   TTTT  EEEEEEETT  EEEE   EEEETTTTTT EEEEEEEETTEEEEEEEEE

Alr21 ECS-trimmed
      |--A---|   |--B--|         |--C-|     |C’|   |E-|       |--F---|  |---G---|
FTPSKVRITSSYRHNASCVYLNWYREDTGNCLLEYHIQFNNINDVYNTSKTNVDICHSPSASSASIWASYKGISGGKVDIIL
      EEEEEEEETTEEEEEEEE   TTTT  EEEEEETTT  EEEE   EEEE  TTTT EEEEEEEETTEEEEEEEEE

Alr23 ECS-trimmed
      |--A---|   |--B--|        |--C---|    |C’|   |-E-|      |--F---|   |---G---|
FVPSPVSMISLSSYNGTCVRVTWDAEDTGKCILNYHVWFSGREIIYNTSNTYFTLCNATDVKNVTIWASYNRDVKGNFTTTSSPDI
      EEEEEEEETTEEEEEEEE   TTTT EEEEEEEETT  EEEE   EEEEETTTTT EEEEEEEETTTEEEEEEEEE

Alr27 ECS-trimmed
  |---A----|  |--B---|       |---C----|  |--C’--|   |-E-|   |---F----|        |G-|
YTPIDLKFEKKSIDCYNLSWSVHPLAKPCVKEYQLVVNQIDKAPLNTTTKTSNYLICSKSIQSCKVRTLCKSNQESDWVTAAI
  EEEEEEEEEETTEEEEEEEE GGGGGGEEEEEEEEEETTEEEEEEEE   EEEEETTTEEEEEEEEEETTT B   EEEE

Alr28 ECS-trimmed
      |--A---|  |--B--|         |--C--|     |-C’--|     |E-|          |----F-----|   |G|
FSPSKVTDFSFAIKDNCQIFQWSTLNSGRCGVFYEIQILDNKKRILSQSTTQPFANFLSFCYAEYTHNNVSAGIRAIYDNKYGNWSSVKP
     TEEEEEEEETTEEEEEEE TTTTTT  EEEEEEETTTT EEEEEEETTTT EEEE TTTTTTTTTEEEEEEEEETTEE   EEE

Alr30p1 ECS-trimmed
      |--A---|  |--B--|        |--C---|     |-C’--|     |E-|          |--F---|        |-G--|
LTPESVKNVNSTEKDGCMYTTWEKVNTGECAVSYRIDYFDSRGLVLFSQNKSEGIYTASMCDEIIISKVNNLRIIAIPNSIFKVHGNGTIVKI
      EEEEEEEETTEEEEEEE  TTTT  EEEEEEEETTTT EEEEEEETTTT EEEE  HHHHH   EEEEEEEE TTTT   EEEEEE

Alr30p3 ECS-trimmed
       |-A--|  |-B--|         |--C---|     |--C’--|    |E-|          |--F---|
LSPVQINSTMRKVGSCIDTNWSAPNTGECDVSYKVDYLDSNNVIVHSKVFHNKELKTESCDTKVVSNTLSVRVTVVSNDTRVEPTNESI
   EE  EEEEEETTEEEEEE   TTTT  EEEEEEEETTT  EEEEEEEE    EEEE  HHHHHH  EEEEEEEE

Alr31 ECS-trimmed
 |---A----|  |--B---|       |--C---|     |-C’--|   |-E-|         |--F---|     |---G---|
AIISIKSLKRKMDCIFTEWTPKHVDKCSIYYEVEYRDRRNDVLWRENTTTNEAIFCSNVSASKVNSVWIRGVFLNEQNVYGDWYIHAL
 EEEEEEEEEETTEEEEEEEE TTTT  EEEEEEEETTTT EEEEEEE   EEEEETTTTGGG  EEEEEEEE TTTTEE   EEEE

Alr33 ECS-trimmed
      |--A---|  |--B--|         |--C---|    |--C’---|   |-E-|         |---F----|  |---G---|
FKPDQVQILSSVVKGRCVKTNWKLLNTGKCNVTYYIEYKVNSENKAIHHASVEGENSYEYCMENILDVKHTGTVYMRAAFGDIKGLWNNLRM
      EEEEEEEETTEEEEEEE   TTTT  EEEEEEEETTTTEEEEEEEEETT EEEEETTTHHHHHHEEEEEEEEEETTEE   EEEE

Alr34 ECS-trimmed
      |--A---|   |--B--|        |---C---|    |-C’--|     |-E-|         |----F-----|  |----G----|
SLPKPVENFKYYPMGGNCFKFTWLAQNTGLCKTHFELQLLLKNNKVKRQSISPMTTGTFIHCEVDSASLKKIYKAKIRSIYQDGATRLEGRWSLLEL
     TEEEEEEEETTEEEEEEEE   TTTT EEEEEEEEETTT EEEEEEETTTT EEEEETTTHHHHHHEEEEEEEEEEEETTEEEE   EEEE

Alr35 ECS-trimmed
      |--A---|  |--B--|        |--C---|     |-C’--|     |-E-|     |--F---|  |---G---|
FTPENVKVTEAYLKEQCVTVRFTTLDVGTCKLSYEFNYFDDRAQLVGSSAADKNTNTVQQCGITASTVKARARSSDSVGQWSAYHI
      EEEEEEEETTEEEEEEE   TTTT EEEEEEEETTTT EEEEEEETTTT EEEEE     EEEEEEEETTEE   EEEE

Alr36 ECS-trimmed
        |-A--|   |-B--|          |-C--|     |C’|   |-E-|      |--F---|  |---G---|
FTPSNNFNATFYYYNSSCVYGTWNEENTGSCHLNYHIQYDDNDAIHLTTKTEYTRCGLTNLKFVQMWAAYNGRVGRKSVYSI
        EEEEEETTEEEEEE     TTTT  EEEEEETTT  EEEE   EEEEE TTTT EEEEEEEETTEE   EEEE

Alr37 ECS-trimmed
       |-A--|  |-B--|          |--C--|     |--C’--|     |-E-|           |--F---|  |---G---|
WTPPVVRFSVNVKENCIFLTWKQPRTGLCAVSYSVTLYGENDVLVYTHFNLSQVKESFKYCSQLLRTINDIKTVGLQAVYKDNFGKVNRRHV
       EEEEEETTEEEEEE    TTTT  EEEEEEE GGG EEEEEEEETTTT EEEEE      GGG  EEEEEEEETTEE   EEEE

Alr38 ECS-trimmed
      |--A---|  |--B--|         |--C---|   |--C’---|    |-E-|         |---F----|  |---G---|
SIPQKIENFQYAIDDTCFVLSWSRQYTGNCIVNHEIQYITGKNTTMNIDIVTSNTNKLSYCAPLPEDIKKIKVIKIRSIYEKRRGEWSSVNI
     TEEEEEEEETTEEEEEEE   TTTT  EEEEEEEETTTEEEEEEEEETTT EEEEETTTGGGGGGEEEEEEEEEETTEE   EEEE
```

**Fig. S16. STRIDE secondary structure predictions for AIr domains 2 and 3**
For each domain, the top line shows beta-strands labeled according to their position in the primary amino acid sequence. The middle line shows the sequence of the domain. The bottom line shows the STRIDE secondary structure predicted from the Colabfold model. (H = alpha helix, G = 3-10 helix, I = PI-helix, E = beta-strand extended conformation, B = isolated bridge, T = turn.)

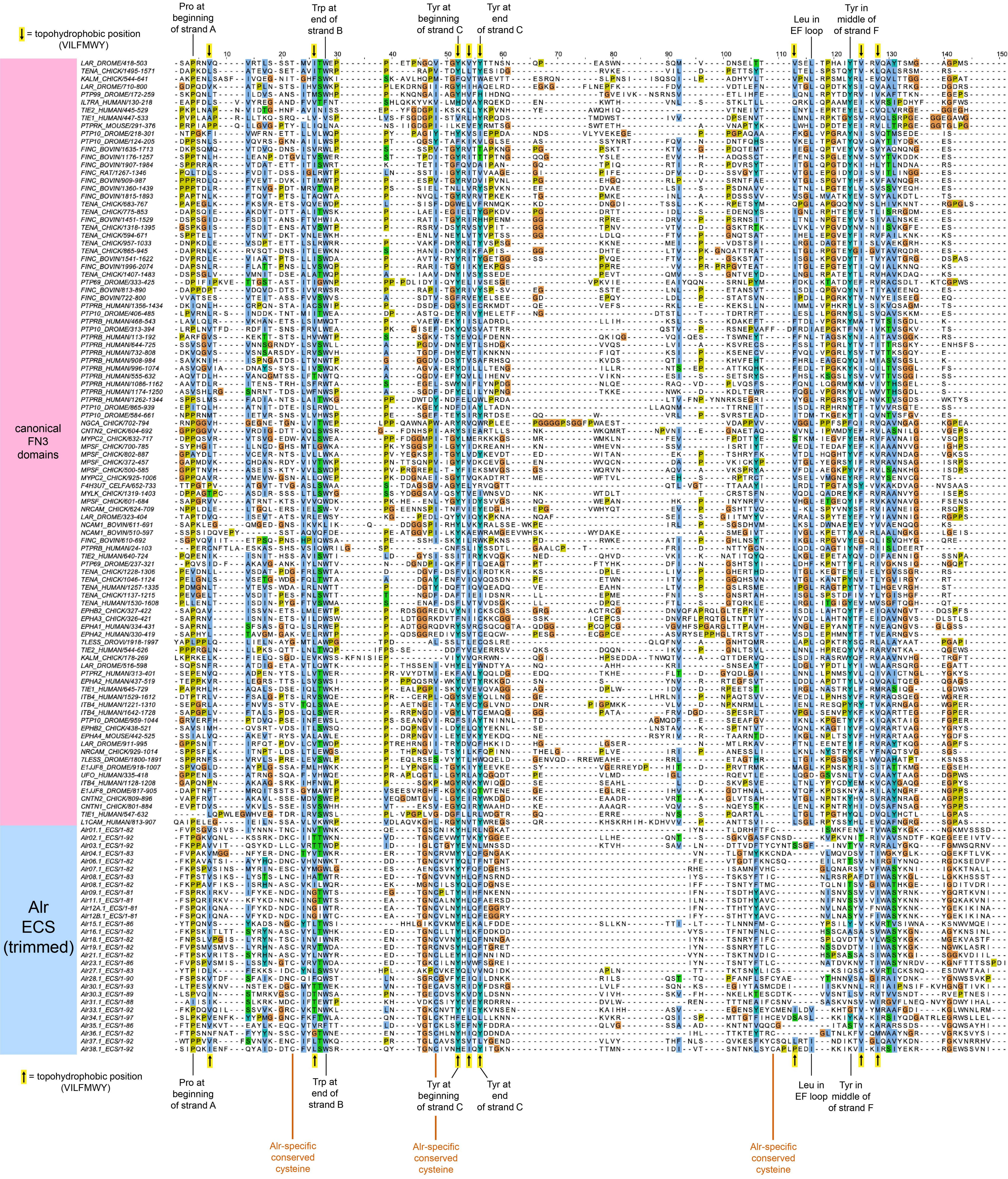

= topohydrophobic position (VILFMWY)
Pro at beginning of strand A
Trp at end of strand B
Tyr at beginning of strand C
Tyr at end of strand C
Leu in EF loop
Tyr in middle of strand F
canonical FN3 domains
Alr ECS (trimmed)
Alr-specific conserved cysteine
Alr-specific conserved cysteine
Alr-specific conserved cysteine

**Fig. S17. Multiple sequence alignment of Fn3 domains and the Alr ECS fold.**
Alr ECS sequences were aligned to Fn3 domains from pfam. Residues in the alignment are highlighted by sequence conservation and chemical property with CLUSTALX colors as implemented in Jalview. The positions of residues typically conserved across Fn3 domains are shown above and below the alignment. The position of invariant cysteine residues is shown in red-orange lettering.

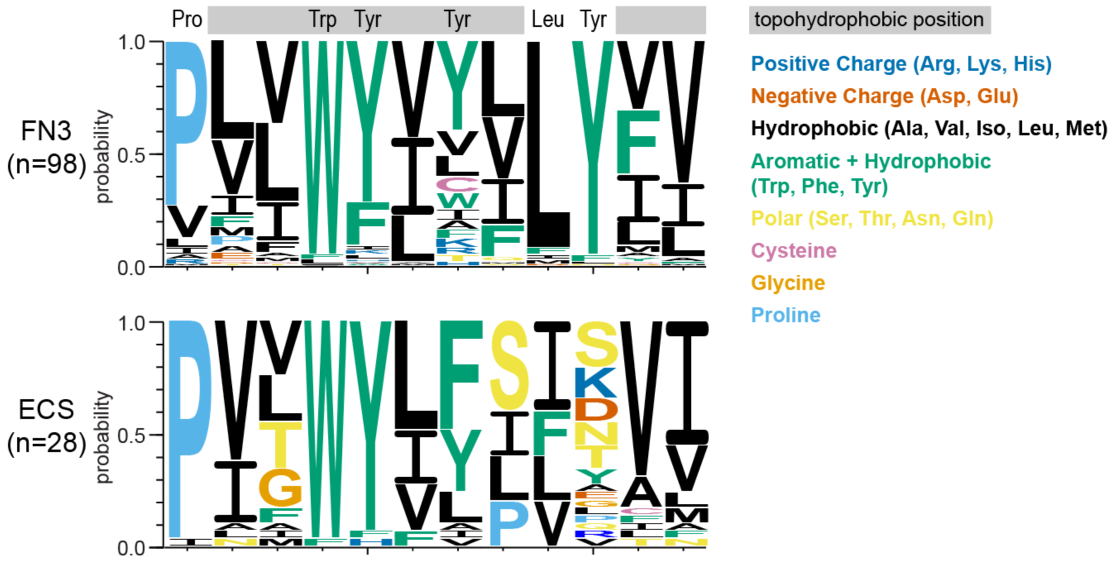


**Fig. S18.** Sequence logo showing residues at conserved positions within Fn3 domains (top) and the ECS fold (bottom).

**Table S1. Overlap coordinates of genomic contigs and BAC contigs used to create reference ARC-F sequence.**

| Genome Assembly ID (length) | start | stop | BAC contigs ID (length) | start | stop |
|---|---|---|---|---|---|
| utg0000000001 (5,049,836 bp) | 687,083 | 1 | bc194 (1,225,536 bp) | 1 | 684,144 |
| utg0000000021 (2,644,760 bp) | 174 | 442,144 | bc194 (1,225,536 bp) | 783,992 | 1,225,536 |
| utg0000000021 (2,644,760 bp) | 1,005,148 | 1,590,913 | bc18 (586,384 bp) | 1 | 586,384 |
| utg0000000121 (601,649 bp) | 386,059 | 170,371 | bc28 (214,692 bp) | 1 | 214,692 |
| utg0000000688 (716,359 bp) | 88,750 | 244 | bc050N15 (147,919 bp) | 1 | 88,530 |
| utg0000000026 (2,721,327 bp) | 2,719,577 | 2,666,436 | bc050N15 (147,919 bp) | 94,782 | 147,919 |
| utg0000000026 (2,721,327 bp) | 2,661,684 | 2,138,742 | bc174 (522,055 bp) | 1 | 522,055 |
| utg0000000026 (2,721,327 bp) | 1,818,447 | 1,610,954 | bc29 (207,512 bp) | 1 | 207,512 |

**Table S2. Gene models classified as *Alr* pseudogenes**

| Gene model name | Expression | Reason for classifying as a pseudogene |
|---|---|---|
| Alr2p2.1 | yes | Partial duplication of exons 1-4 of Alr2. Frame-shift in exon 3 leading to premature stop codons in exon 3. |
| Alr2p2.2 | yes | Partial duplication of exons 1-5 of Alr2. Frame-shift in exon 3 leading to premature stop codons in exon 3. |
| Alr05p | yes | No evidence of splicing between exons 2-3. |
| Alr10 | yes | Improper splicing of exon 4 to downstream exons introduces stop codon in transcript. |
| Alr12C | no | Stop codon in exon 2. |
| Alr13 | no | Stop codon in exon 1. |
| Alr14p | no | No evidence of expression or exon encoding signal peptide. |
| Alr20p | Yes | No evidence of expression or splicing between exons 1 and 2. No evidence of exon encoding a signal peptide. |
| Alr22p | no | No evidence of expression or exon encoding signal peptide. |
| Alr24p | yes | A few reads map to exons 2-3. No evidence of exon encoding a signal peptide. |
| Alr25p | yes | A few reads map to exons 2-5. No evidence of exon encoding a signal peptide. |
| Alr26p | yes | No open reading frame. No evidence of exon encoding a signal peptide. No splicing between exon 2-3. |
| Alr32p | no | Only three exons, which have sequence similarity to Alr31, but are not expressed. |

**Table S3. Sequence homology of Domain 1**

| Protein[a] | Domain | HMMER search of pfam | | | HHpred search of SCOPe | | |
|---|---|---|---|---|---|---|---|
| | | *Accession* | *description* | *e-value*[b] | *Accession* | *SCOPe Family* | *probability*[c] |
| Alr1 | D1 | | | | d5l21b | b.1.1.1: V set domains | 96.4 |
| Alr2 | D1 | PF07686.17 | V-set | 0.0012 | d5e56a | b.1.1.1: V set domains | 95.8 |
| Alr3 | D1 | | | | | | |
| Alr4 | D1 | | | | d5my6b1 | b.1.1.1: V set domains | 96.5 |
| Alr6 | D1 | | | | d5my6b1 | b.1.1.1: V set domains | 96.5 |
| Alr7 | D1 | | | | d5l21b | b.1.1.1: V set domains | 96.0 |
| Alr8 | D1a[d] | | | | d5my6b1 | b.1.1.1: V set domains | 95.7 |
| Alr8 | D1b[e] | | | | d2esve1 | b.1.1.1: V set domains | 96.0 |
| Alr9 | D1 | | | | d5my6b1 | b.1.1.1: V set domains | 96.1 |
| Alr11 | D1 | | | | d5my6b1 | b.1.1.1: V set domains | 96.7 |
| Alr12A | D1 | | | | d4n8pa1 | b.1.1.1: V set domains | 96.6 |
| Alr12B | D1 | | | | d5my6b1 | b.1.1.1: V set domains | 96.9 |
| Alr15 | D1 | | | | | | |
| Alr16 | D1 | | | | d5my6b1 | b.1.1.1: V set domains | 96.9 |
| Alr17 | D1 | | | | d5my6b1 | b.1.1.1: V set domains | 97.1 |
| Alr18 | D1 | | | | d5my6b1 | b.1.1.1: V set domains | 97.0 |
| Alr19 | D1 | | | | d5my6b1 | b.1.1.1: V set domains | 96.4 |
| Alr21 | D1 | PF07686.17 | V-set | 0.0012 | d5my6b1 | b.1.1.1: V set domains | 96.8 |
| Alr23 | D1 | | | | d5my6b1 | b.1.1.1: V set domains | 96.9 |
| Alr27 | D1 | | | | | | |
| Alr28 | D1 | PF07686.17 | V-set | 1.40E-04 | d5o04f1 | b.1.1.1: V set domains | 95.3 |
| Alr29 | D1 | PF07686.17 | V-set | 8.10E-05 | d1yjdc1 | b.1.1.1: V set domains | 95.5 |
| Alr30 | D1 | PF07686.17 | V-set | 0.0031 | d5e56a | b.1.1.1: V set domains | 95.0 |
| Alr31 | D1 | PF17711.1 | DUF5556 | 0.0097 | | | |
| Alr33 | D1 | | | | | | |
| Alr34 | D1 | | | | d1c5db1 | b.1.1.1: V set domains | 88.8 |
| Alr35 | D1 | PF07686.17 | V-set | 1.00E-04 | d5o04f1 | b.1.1.1: V set domains | 93.9 |
| Alr36 | D1 | | | | d5my6b1 | b.1.1.1: V set domains | 93.4 |
| Alr38 | D1 | | | | d5my6b1 | b.1.1.1: V set domains | 69.5 |

[a] proteins encoded by *bona fide* genes in blue, putative genes in red
[b] significance cutoff = 0.01
[c] probability of homology; values <50% not shown; values >95% shaded in green
[d] this is the membrane-distal domain with homology to other domain 1 sequences
[e] this is the membrane-proximal domain with homology to other domain 1 sequences

**Table S4. Predicted structural homology for domain 1**

| Protein[a] | Domain | Colabfold | DALI Top Structural Alignment | | | | | PDBeFold | | | | |
|---|---|---|---|---|---|---|---|---|---|---|---|---|
| | | *plDDT score[b]* | *PDB accession* | *Z-score[c]* | *RMSD* | *% ID* | *Domain Type[d]* | *PDB accession* | *Q-score[e]* | *RMSD* | *% ID* | *Domain Type[d]* |
| Alr1 | D1 | 97.4 | 7kqyE | 15.2 | 1.9 | 10 | V-set | 5uoeN | 0.6356 | 1.589 | 9.9 | V-set |
| Alr2 | D1 | 90.5 | 3oaiA | 15.1 | 2.1 | 20 | V-set | 3m45C | 0.6243 | 1.663 | 14.7 | V-set |
| Alr3 | D1 | 95.2 | 3udwD | 14.8 | 1.9 | 14 | V-set | 3udwD | 0.6095 | 1.565 | 14.6 | V-set |
| Alr4 | D1 | 92.7 | 5imkA | 14.4 | 1.8 | 16 | V-set | 2iceT | 0.6337 | 1.708 | 16.5 | V-set |
| Alr6 | D1 | 97.0 | 6o3bE | 15.7 | 1.5 | 14 | V-set | 5immA | 0.6647 | 1.695 | 13.6 | V-set |
| Alr7 | D1 | 92.5 | 2iceS | 14.9 | 2.1 | 11 | V-set | 1neuA | 0.554 | 1.923 | 15.1 | V-set |
| Alr8 | D1a[f] | 94.7 | 2iceT | 15.2 | 2.1 | 13 | V-set | 5immA | 0.6055 | 1.874 | 13.5 | V-set |
| Alr8 | D1b[g] | 89.3 | 5imkA | 13.5 | 2.1 | 11 | V-set | 5immA | 0.5658 | 1.875 | 12.3 | V-set |
| Alr9 | D1 | 96.9 | 6o3bE | 14.7 | 1.5 | 11 | V-set | 6krzG | 0.6162 | 1.581 | 13.3 | V-set |
| Alr11 | D1 | 96.7 | 6o3bE | 15.7 | 1.6 | 14 | V-set | 2iceS | 0.663 | 1.634 | 16.7 | V-set |
| Alr12A | D1 | 95.4 | 5imkA | 14.7 | 1.8 | 15 | V-set | 2pndA | 0.6482 | 1.708 | 15.1 | V-set |
| Alr12B | D1 | 92.8 | 5imkA | 14.9 | 2 | 16 | V-set | 2iceT | 0.6412 | 1.748 | 15.9 | V-set |
| Alr15 | D1 | 85.6 | 6bj2D | 14 | 2.1 | 15 | V-set | 1u3hE | 0.5279 | 1.952 | 14.0 | V-set |
| Alr16 | D1 | 95.7 | 2iceT | 14.6 | 2.1 | 10 | V-set | 2iceT | 0.6332 | 1.802 | 10.9 | V-set |
| Alr17 | D1 | 95.5 | 2iceS | 15 | 2.1 | 11 | V-set | 2iceT | 0.6197 | 1.879 | 11.7 | V-set |
| Alr18 | D1 | 95.7 | 3qi9D | 14.9 | 1.8 | 10 | V-set | 6oppL | 0.612 | 1.515 | 16.2 | V-set |
| Alr19 | D1 | 97.3 | 1tvdB | 15.3 | 2.2 | 9 | V-set | 5uoeN | 0.6353 | 1.59 | 9.9 | V-set |
| Alr21 | D1 | 95.0 | 6j8gC | 14.4 | 2 | 13 | V-set | 6j8hC | 0.6408 | 1.795 | 11.7 | V-set |
| Alr23 | D1 | 95.9 | 2pndA | 15.3 | 1.7 | 11 | V-set | 5immA | 0.6338 | 1.668 | 12.2 | V-set |
| Alr27 | D1 | 84.2 | 2f53D | 14.1 | 2 | 9 | V-set | 1u3hE | 0.5468 | 1.754 | 11.2 | V-set |
| Alr28 | D1 | 80.6 | 5m2wB | 14 | 2.1 | 9 | V-set | 3ucrA | 0.5733 | 1.817 | 19.2 | V-set |
| Alr29 | D1 | 88.7 | 2iceT | 17.1 | 1.6 | 10 | V-set | 2iceT | 0.7063 | 1.569 | 11.0 | V-set |
| Alr30 | D1 | 92.8 | 3oaiA | 15.8 | 2 | 19 | V-set | 6ol7L | 0.6295 | 1.419 | 19.4 | V-set |
| Alr31 | D1 | 86.7 | 2iceT | 14.2 | 2 | 13 | V-set | 2ptvA | 0.6401 | 1.586 | 11.8 | V-set |
| Alr33 | D1 | 84.3 | 6dleB | 12.6 | 1.6 | 11 | Ig domain | 6arqA | 0.5361 | 1.811 | 13.4 | V-set |
| Alr34 | D1 | 95.7 | 5imkA | 15 | 2.4 | 20 | V-set | 6vi4C | 0.6021 | 2.113 | 20.5 | V-set |
| Alr35 | D1 | 93.5 | 1tvdA | 16.7 | 2.2 | 15 | V-set | 3b9kA | 0.639 | 1.771 | 12.7 | V-set |
| Alr36 | D1 | 92.9 | 6fr6B | 15.3 | 1.9 | 13 | V-set | 3udwA | 0.6006 | 1.884 | 14.9 | V-set |
| Alr38 | D1 | 93.4 | 5imlA | 15.8 | 2.2 | 16 | V-set | 2iccA | 0.6086 | 1.944 | 16.1 | V-set |

[a] *bona fide* genes in blue, putative genes in red
[b] predicted local-distance difference test score; >90 considered highly accurate
[c] Z-score between 8-20 indicates probable homology between query and hit
[d] as annotated in the PDB (rcsb.org)
[e] Q-score = 1 are identical alignments; >0.5 are considered to have homologous structures
[f] this is the membrane-distal domain with homology to other domain 1 sequences in Alr8
[g] this is the membrane-proximal domain with homology to other domain 1 sequences in Alr8

### Table S5. Sequence homology for Domain 2 and 3

| Protein[a] | Domain | HMMER search of pfam | | | HHpred search of SCOPe | | |
|---|---|---|---|---|---|---|---|
| | | *Accession* | *description* | *e-value*[b] | *Accession* | *SCOPe family* | *probability*[c] |
| Alr1 | D2 | PF07679.16 | I-set | 3.30E-05 | d1biha3 | b.1.1.4: I-set domains | 97.6 |
| Alr2 | D2 | PF13927.6 | Ig_3 | 0.0002 | d1biha3 | b.1.1.4: I-set domains | 99.0 |
| Alr2 | *D3* | PF07679.16 | I-set | 0.0023 | d1biha3 | b.1.1.4: I-set domains | 96.7 |
| Alr3 | D2 | | | | d1biha3 | b.1.1.4: I-set domains | 96.6 |
| Alr4 | D2 | | | | d1biha3 | b.1.1.4: I-set domains | 98.7 |
| Alr6 | D2 | PF07679.16 | I-set | 1.70E-05 | d1biha3 | b.1.1.4: I-set domains | 97.0 |
| Alr7 | D2 | PF07679.16 | I-set | 8.80E-05 | d1x44a1 | b.1.1.4: I-set domains | 99.3 |
| Alr8 | D2a[d] | PF07679.16 | I-set | 0.0024 | d1biha3 | b.1.1.4: I-set domains | 97.4 |
| Alr8 | D2b[e] | PF07679.16 | I-set | 2.20E-07 | d1biha3 | b.1.1.4: I-set domains | 98.1 |
| Alr9 | D2 | | | | d1biha3 | b.1.1.4: I-set domains | 97.4 |
| Alr11 | D2 | PF07679.16 | I-set | 0.0034 | d1biha3 | b.1.1.4: I-set domains | 97.1 |
| Alr12A | D2 | PF13927.6 | Ig_3 | 0.0028 | d1biha3 | b.1.1.4: I-set domains | 97.0 |
| Alr12B | D2 | PF13927.6 | Ig_3 | 0.0024 | d1biha3 | b.1.1.4: I-set domains | 97.2 |
| Alr15 | D2 | | | | d1biha3 | b.1.1.4: I-set domains | 84.9 |
| Alr16 | D2 | PF07679.16 | I-set | 0.0021 | d1biha3 | b.1.1.4: I-set domains | 97.4 |
| Alr18 | D2 | PF07679.16 | I-set | 3.50E-06 | d1biha3 | b.1.1.4: I-set domains | 97.4 |
| Alr19 | D2 | PF07679.16 | I-set | 2.50E-06 | d1biha3 | b.1.1.4: I-set domains | 97.4 |
| Alr21 | D2 | PF07679.16 | I-set | 8.10E-05 | d1biha3 | b.1.1.4: I-set domains | 97.6 |
| Alr23 | D2 | PF07679.16 | I-set | 0.0005 | d1biha3 | b.1.1.4: I-set domains | 97.5 |
| Alr27 | D2 | | | | | | |
| Alr28 | D2 | PF07679.16 | I-set | 0.0076 | d1vcaa2 | b.1.1.4: I-set domains | 97.4 |
| Alr29 | D2 | | | | d1ncua1 | b.1.1.4: I-set domains | 85.2 |
| Alr30 | D2 | PF13927.6 | Ig_3 | 0.00027 | d1biha3 | b.1.1.4: I-set domains | 97.8 |
| Alr30 | *D3* | | | | d1biha3 | b.1.1.4: I-set domains | 96.9 |
| Alr31 | D2 | | | | d1ncua1 | b.1.1.4: I-set domains | 87.2 |
| Alr33 | D2 | | | | d1iray3 | b.1.1.4: I-set domains | 54.4 |
| Alr34 | D2 | | | | d1biha3 | b.1.1.4: I-set domains | 97.1 |
| Alr35 | D2 | PF07679.16 | I-set | 0.0066 | d1koaa1 | b.1.1.4: I-set domains | 89.8 |
| Alr36 | D2 | | | | d1biha3 | b.1.1.4: I-set domains | 97.4 |
| Alr37 | D2 | | | | d1biha3 | b.1.1.4: I-set domains | 78.7 |
| Alr38 | D2 | | | | d1iray3 | b.1.1.4: I-set domains | 54.7 |

[a] proteins encoded by *bona fide* genes in blue, putative genes in orange
[b] significance cutoff = 0.01
[c] probability of homology; values <80% not shown; values >95% shaded in green
[d] this is the membrane-distal domain with homology to other domain 1 sequences
[e] this is the membrane-proximal domain with homology to other domain 1 sequences

**Table S6. Predicted structural homology for domains 2 and 3**

| Protein[a] | Domain | Colabfold | DALI Top Structural Alignment | | | | | PDBeFold Top Structural Alignment | | | | |
|---|---|---|---|---|---|---|---|---|---|---|---|---|
| | | *plDDT score*[b] | *PDB accession* | *Z-score*[c] | *RMSD* | *% ID* | *Domain Type*[d] | *PDB accession* | *Q-score*[e] | *RMSD* | *% ID* | *Domain Type*[d] |
| Alr1 | D2 | 95.1 | 2rjmA | 12.9 | 1.7 | 19 | I-set | 3qp3B | 0.5999 | 1.5 | 16 | I-set |
| Alr2 | D2 | 93.7 | 1u2hA | 12.4 | 1.7 | 16 | I-set | 1u2hA | 0.6237 | 1.6 | 16 | I-set |
| Alr2 | D3 | 90.9 | 6efyA | 12.9 | 1.5 | 23 | I-set | 3qp3C | 0.6663 | 1.3 | 15 | I-set |
| Alr3 | D2 | 88.2 | 2rikA | 12.7 | 1.8 | 11 | I-set | 6h4lA | 0.6153 | 1.7 | 13 | I-set |
| Alr4 | D2 | 94.4 | 2j8hA | 13.2 | 1.4 | 18 | I-set | 3pucA | 0.6466 | 1.3 | 15 | I-set |
| Alr6 | D2 | 93.9 | 2rjmA | 12.2 | 1.7 | 15 | I-set | 6h4lA | 0.6019 | 1.5 | 12 | I-set |
| Alr7 | D2 | 92.1 | 2rjmA | 13.0 | 1.5 | 16 | I-set | 4uowK | 0.6176 | 1.5 | 16 | I-set |
| Alr8 | D2a | 88.0 | 2rjmA | 13.4 | 1.8 | 21 | I-set | 1u2hA | 0.6517 | 1.5 | 18 | I-set |
| Alr8 | D2b | 92.5 | 2rjmA | 13.6 | 1.4 | 16 | I-set | 6h4lA | 0.6649 | 1.4 | 17 | I-set |
| Alr9 | D2 | 81.1 | 4pgzA | 10.5 | 2.6 | 16 | I-set | 3j9f8 | 0.5011 | 2.3 | 12 | I-set/C2-set |
| Alr11 | D2 | 88.2 | 2illA | 12.1 | 1.6 | 21 | I-set | 1g1cB | 0.5566 | 1.7 | 19 | I-set |
| Alr12A | D2 | 88.4 | 3pucA | 12.9 | 1.6 | 10 | I-set | 1g1cA | 0.5729 | 1.7 | 23 | I-set |
| Alr12B | D2 | 85.7 | 4of8B | 12.0 | 2.1 | 13 | I-set/C2-set | 2wwmT | 0.5027 | 2.1 | 22 | I-set |
| Alr15 | D2 | 92.6 | 4of8B | 10.8 | 2.1 | 12 | I-set/C2-set | 3rghB | 0.5252 | 2.0 | 5 | filamin |
| Alr16 | D2 | 85.9 | 4uow5 | 11.5 | 2.1 | 19 | I-set | 3j9f8 | 0.4915 | 2.4 | 8 | I-set/C2-set |
| Alr18 | D2 | 89.4 | 2rjmA | 13.1 | 1.7 | 24 | I-set | 4uowG | 0.6208 | 1.6 | 22 | I-set |
| Alr19 | D2 | 92.6 | 2rjmA | 12.1 | 1.8 | 23 | I-set | 1g1cA | 0.588 | 1.5 | 23 | I-set |
| Alr21 | D2 | 87.5 | 6efyA | 12.9 | 2.0 | 12 | I-set | 6h4lA | 0.6162 | 1.5 | 8 | I-set |
| Alr23 | D2 | 92.2 | 4pgzB | 11.6 | 2.3 | 14 | I-set | 6h4lA | 0.5576 | 1.8 | 19 | I-set |
| Alr27 | D2 | 92.4 | 3sbwC | 10.7 | 2.3 | 14 | I-set/C2-set | 4uowB | 0.4968 | 1.8 | 14 | I-set |
| Alr28 | D2 | 91.8 | 4pgzB | 13.0 | 1.9 | 16 | I-set | 6h4lA | 0.6368 | 1.3 | 16 | I-set |
| Alr29 | D2 | 86.9 | 4of8B | 10.6 | 2.2 | 15 | I-set/C2-set | 2wwkT | 0.493 | 1.8 | 16 | I-set |
| Alr30 | D2 | 90.4 | 1u2hA | 12.9 | 1.6 | 23 | I-set | 1u2hA | 0.6853 | 1.4 | 24 | I-set |
| Alr30 | D3 | 92.1 | 2fdbP | 12.3 | 1.8 | 12 | I-set | 4uowE | 0.5936 | 1.7 | 12 | I-set |
| Alr31 | D2 | 84.0 | 3dmkC | 11.4 | 2.4 | 13 | I-set | 6h4lA | 0.5577 | 1.7 | 15 | I-set |
| Alr33 | D2 | 89.0 | 6pv9A | 10.4 | 2.2 | 7 | I-set/C2-set | 2kdgA | 0.5145 | 1.8 | 21 | I-set |
| Alr34 | D2 | 93.4 | 2j8hA | 12.8 | 1.5 | 20 | I-set | 3pucA | 0.6343 | 1.5 | 17 | I-set |
| Alr35 | D2 | 95.5 | 3dmkC | 12.4 | 2.3 | 15 | I-set | 6h4lA | 0.5833 | 1.9 | 16 | I-set |
| Alr36 | D2 | 90.0 | 2rikA | 13.4 | 1.7 | 19 | I-set | 6h4lA | 0.6571 | 1.4 | 13 | I-set |
| Alr37 | D2 | 92.6 | 4uowR | 10.9 | 2.2 | 13 | I-set | 4uowN | 0.5459 | 1.9 | 12 | I-set |
| Alr38 | D2 | 91.7 | 2rikA | 12.5 | 1.7 | 16 | I-set | 1u2hA | 0.6147 | 1.4 | 17 | I-set |

[a] *bona fide* genes in blue, putative genes in red
[b] predicted local-distance difference test score; >90 considered highly accurate
[c] Z-score between 8-20 indicates probable homology between query and hit
[d] as annotated in the PDB (rcsb.org)
[e] Q-score = 1 are identical alignments; >0.5 are considered to have homologous structures
[f] this is the membrane-distal domain with homology to other domain 1 sequences in Alr8
[g] this is the membrane-proximal domain with homology to other domain 1 sequences in Alr8

### Table S7. Sequence homology of the ECS fold

| Protein[a] | Domain | HMMER search of pfam | | | HHpred search of SCOPe | | |
|---|---|---|---|---|---|---|---|
| | | *Accession* | *description* | *e-value*[b] | *Accession* | *SCOPe family* | *Probability*[c] |
| Alr1 | ECS | | | | d1j8ka | b.1.2.1: Fibronectin type III | 88.1 |
| Alr2 | ECS | | | | d1fyhb1 | b.1.2.1: Fibronectin type III | 84.7 |
| Alr3 | ECS | | | | d3s9db1 | b.1.2.1: Fibronectin type III | 76.6 |
| Alr4 | ECS | | | | | | |
| Alr6 | ECS | | | | d1j8ka | b.1.2.1: Fibronectin type III | 88.9 |
| Alr7 | ECS | | | | d1fnfa1 | b.1.2.1: Fibronectin type III | 79.7 |
| Alr8 | ECSa[e] | | | | d1j8ka | b.1.2.1: Fibronectin type III | 85.1 |
| Alr8 | ECSb[f] | | | | | | |
| Alr9 | ECS | | | | d1j8ka | b.1.2.1: Fibronectin type III | 81.6 |
| Alr11 | ECS | | | | d1j8ka | b.1.2.1: Fibronectin type III | 82.4 |
| Alr12A | ECS | | | | d1j8ka | b.1.2.1: Fibronectin type III | 81.3 |
| Alr12B | ECS | | | | d1j8ka | b.1.2.1: Fibronectin type III | 80.2 |
| Alr15 | ECS | | | | | | |
| Alr16 | ECS | | | | d1j8ka | b.1.2.1: Fibronectin type III | 85.2 |
| Alr18 | ECS | | | | d1j8ka | b.1.2.1: Fibronectin type III | 87.8 |
| Alr19 | ECS | | | | d1j8ka | b.1.2.1: Fibronectin type III | 87.8 |
| Alr21 | ECS | | | | d1j8ka | b.1.2.1: Fibronectin type III | 82.0 |
| Alr23 | ECS | | | | d1fnfa1 | b.1.2.1: Fibronectin type III | 79.8 |
| Alr27 | ECS | | | | | | |
| Alr28 | ECS | | | | d1fyhb1 | b.1.2.1: Fibronectin type III | 75.6 |
| Alr29 | ECS | | | | | | |
| Alr30.1 | ECS | | | | d1fyhb1 | b.1.2.1: Fibronectin type III | 70.9 |
| Alr30.3 | ECS | | | | d1fyhb1 | b.1.2.1: Fibronectin type III | 82.4 |
| Alr31 | ECS | | | | d3d85d3 | b.1.2.1: Fibronectin type III | 56.8 |
| Alr33 | ECS | PF07403.13 | DUF1505 | 0.0013 | | | |
| Alr34 | ECS | | | | | | |
| Alr35 | ECS | | | | | | |
| Alr36 | ECS | | | | d1fnfa1 | b.1.2.1: Fibronectin type III | 80.8 |
| Alr37 | ECS | | | | d1fnfa1 | b.1.2.1: Fibronectin type III | 74.3 |
| Alr38 | ECS | | | | d2gysa2 | b.1.2.1: Fibronectin type III | 60.5 |

[a] proteins encoded by *bona fide* genes in blue, putative genes in red
[b] significance cutoff = 0.01
[c] probability of homology; values <50% not shown; values >95% shaded in green
[d] this is the membrane-distal domain with homology to other domain 1 sequences
[e] this is the membrane-proximal domain with homology to other domain 1 sequences

**Table S8. Predicted structural homology for the immunoglobulin-like fold in the ECS**

| Protein[a] | Domain | Colabfold | DALI Top Structural Alignment | | | | | PDBeFold Top Structural Alignment | | | | |
|---|---|---|---|---|---|---|---|---|---|---|---|---|
| | | *plDDT score[b]* | *PDB accession* | *Z-score[c]* | *RMSD* | *% ID* | *Domain Type[d]* | *PDB accession* | *Q-score[e]* | *RMSD* | *% ID* | *Domain Type[d]* |
| Alr1 | ECS | 90.0 | 6h41A | 11.4 | 1.8 | 13 | Fn3 | 7jguA | 0.5781 | 1.5 | 16 | Fn3 |
| Alr2 | ECS | 95.6 | 5fn8A | 12.9 | 1.5 | 22 | Fn3 | 5dc0A | 0.6019 | 1.6 | 12 | Fn3 |
| Alr3 | ECS | 94.1 | 5fn6A | 12.4 | 1.7 | 10 | Fn3 | 5n48D | 0.5855 | 1.8 | 8 | Fn3 |
| Alr4 | ECS | 95.4 | 7e9jB | 12.7 | 1.8 | 15 | Fn3 | 7jgtA | 0.5270 | 2.0 | 13 | Fn3 |
| Alr6 | ECS | 94.0 | 7e9kD | 13.2 | 1.4 | 16 | Fn3 | 1jrhI | 0.5330 | 1.6 | 5 | Fn3 |
| Alr7 | ECS | 90.0 | 5fn6A | 12.2 | 1.7 | 9 | Fn3 | 5n48D | 0.5501 | 1.9 | 10 | Fn3 |
| Alr8[g] | ECSa | 91.1 | 7e9jB | 13.0 | 1.5 | 14 | Fn3 | 2rb8A | 0.5490 | 2.2 | 10 | Fn3 |
| Alr8[h] | ECSb | 92.1 | 5fn6A | 13.4 | 1.8 | 10 | Fn3 | 2rb8A | 0.5852 | 1.8 | 6 | Fn3 |
| Alr9 | ECS | 94.7 | 5fn8A | 13.6 | 1.4 | 16 | Fn3 | 7jguA | 0.5956 | 1.6 | 16 | Fn3 |
| Alr11 | ECS | 95.1 | 5fn8A | 10.5 | 2.6 | 18 | Fn3 | 1tenA | 0.5988 | 1.7 | 11 | Fn3 |
| Alr12A | ECS | 95.3 | 5fn8A | 12.1 | 1.6 | 14 | Fn3 | 7jguA | 0.5975 | 1.6 | 12 | Fn3 |
| Alr12B | ECS | 94.2 | 5x83B | 12.9 | 1.6 | 9 | Fn3 | 7jguA | 0.5947 | 1.6 | 11 | Fn3 |
| Alr15 | ECS | 93.9 | 2geeA | 12.0 | 2.1 | 10 | Fn3 | 3rzwA | 0.6084 | 1.6 | 8 | Fn3 |
| Alr16 | ECS | 93.7 | 5fn6A | 10.8 | 2.1 | 13 | Fn3 | 1tenA | 0.5925 | 1.8 | 6 | Fn3 |
| Alr18 | ECS | 93.1 | 6h41A | 11.5 | 2.1 | 8 | Fn3 | 7jguA | 0.6015 | 1.6 | 12 | Fn3 |
| Alr19 | ECS | 94.5 | 5fn6A | 13.1 | 1.7 | 14 | Fn3 | 7jguA | 0.5960 | 1.7 | 14 | Fn3 |
| Alr21 | ECS | 93.9 | 5fn6A | 12.1 | 1.8 | 13 | Fn3 | 5n48D | 0.5829 | 1.9 | 10 | Fn3 |
| Alr23 | ECS | 84.7 | 6xfiA | 12.9 | 2.0 | 15 | Fn3 | 4wtwB | 0.5724 | 1.6 | 18 | Fn3 |
| Alr27 | ECS | 94.3 | 2geeA | 11.6 | 2.3 | 13 | Fn3 | 5oc7B | 0.6465 | 1.7 | 15 | Fn3 |
| Alr28 | ECS | 92.4 | 3t1wA | 10.7 | 2.3 | 11 | Fn3 | 5n48D | 0.6033 | 1.9 | 12 | Fn3 |
| Alr30.1 | ECS | 88.2 | 3t1wA | 13.0 | 1.9 | 14 | Fn3 | 4wtwA | 0.5870 | 1.8 | 14 | Fn3 |
| Alr30.3 | ECS | 85.9 | 6mojB | 10.6 | 2.2 | 12 | Fn3 | 5n06A | 0.4340 | 2.2 | 15 | Fn3 |
| Alr31 | ECS | 92.4 | 5fn8B | 12.9 | 1.6 | 12 | Fn3 | 5dc0A | 0.5683 | 2.0 | 7 | Fn3 |
| Alr33 | ECS | 89.6 | 3t1wA | 12.3 | 1.8 | 10 | Fn3 | 2rb8A | 0.6074 | 2.0 | 10 | Fn3 |
| Alr34 | ECS | 92.3 | 5n48D | 11.4 | 2.4 | 7 | Fn3 | 5dc0A | 0.5811 | 2.0 | 7 | Fn3 |
| Alr35 | ECS | 96.5 | 5n48D | 10.4 | 2.2 | 7 | Fn3 | 5n48B | 0.6154 | 1.9 | 7 | Fn3 |
| Alr36 | ECS | 93.1 | 5fn8B | 12.8 | 1.5 | 8 | Fn3 | 7jguA | 0.5872 | 1.7 | 20 | Fn3 |
| Alr37 | ECS | 92.0 | 5n48D | 12.4 | 2.3 | 9 | Fn3 | 5n48D | 0.5964 | 1.9 | 10 | Fn3 |
| Alr38 | ECS | 92.2 | 5n48D | 13.4 | 1.7 | 11 | Fn3 | 5n48D | 0.6182 | 1.8 | 9 | Fn3 |

[a] *bona fide* genes in blue, putative genes in red
[b] predicted local-distance difference test score; >90 considered highly accurate
[c] Z-score between 8-20 indicates probable homology between query and hit
[d] as annotated in the PDB (rcsb.org)
[e] Q-score = 1 are identical alignments; >0.5 are considered to have homologous structures
[f] this is the membrane-distal domain with homology to other domain 1 sequences in Alr8
[g] this is the membrane-proximal domain with homology to other domain 1 sequences in Alr8

**Table S9. Structural predictions of tandem I-set and FnIII-like domains**

| Protein[a] | Tandem Domains I-set/FnIII-like | Colabfold *plDDT score*[b] |
|---|---|---|
| Alr1 | D2-ECS | 92.2 |
| Alr2 | D3-ECS | 92.5 |
| Alr3 | D2-ECS | 89.8 |
| Alr4 | D2-ECS | 95.2 |
| Alr6 | D2-ECS | 93.6 |
| Alr7 | D2-ECS | 92.7 |
| Alr8 | D2a-ECSa | 91.4 |
| Alr8 | D2b-ECSb | 92.9 |
| Alr9 | D2-ECS | 92.0 |
| Alr11 | D2-ECS | 93.7 |
| Alr12A | D2-ECS | 92.6 |
| Alr12B | D2-ECS | 93.2 |
| Alr15 | D2-ECS | 76.7 |
| Alr16 | D2-ECS | 91.9 |
| Alr18 | D2-ECS | 92.3 |
| Alr19 | D2-ECS | 94.7 |
| Alr21 | D2-ECS | 92.5 |
| Alr23 | D2-ECS | 85.0 |
| Alr27 | D2-ECS | 91.7 |
| Alr28 | D2-ECS | 90.0 |
| Alr30 | D3-ECS | 87.6 |
| Alr31 | D2-ECS | 74.2 |
| Alr33 | D2-ECS | 87.3 |
| Alr34 | D2-ECS | 88.2 |
| Alr35 | D2-ECS | 95.7 |
| Alr36 | D2-ECS | 93.9 |
| Alr37 | D2-ECS | 86.4 |
| Alr38 | D2-ECS | 88.0 |

[a] *bona fide* genes in blue, putative genes in red

[b] predicted local-distance difference test score; values >90 are considered highly accurate

### Supplemental Table 10. BLASTP results for full-length Alr protein sequences

| Query[a] | *vs. Hydra* Accession | SP[b] | TM[c] | E-value[d] | *vs. Clytia* Accession | SP[b] | TM[c] | E-value[d] | *vs. nr, excluding Hydrozoans* Description | Accession | E-value[d] |
|---|---|---|---|---|---|---|---|---|---|---|---|
| Alr1 | XP_012557852 | - | - | 1E-36 | 00052857 | - | - | 8E-19 | hemicentin-2-like [Ctenocephalides felis] | XP_026479882 | 0.29 |
| Alr2 | XP_012563475 | - | - | 4E-05 | 00064849 | - | - | 1E-13 | unnamed protein product [Brachionus calyciflorus] | CAF0791325 | 6E-6 |
| Alr3 | XP_012563475 | + | + | 6E-10 | 00052857 | + | + | 8E-25 | hemicentin-2 [Sparus aurata] | XP_030273182 | 4.9 |
| Alr4 | XP_012557852 | + | + | 8E-32 | 00052857 | + | + | 2E-15 | uncharacterized protein [Saprochaete ingens] | XP_031856852 | 0.086 |
| Alr6 | XP_012557852 | + | + | 1E-26 | 00052857 | + | + | 9E-18 | unnamed protein [Tenebrio molitor] | CAG9032869 | 0.004 |
| Alr7 | XP_012563475 | + | + | 3E-12 | 00068309 | + | + | 4E-19 | matrilin-2-like [Branchiostoma floridae] | XP_035689613 | 4E-10 |
| Alr8 | XP_012563475 | - | + | 1E-10 | 00063453 | - | + | 8E-15 | hypothetical protein [Anopheles sinensis] | KFB45464 | 8E-4 |
| Alr9 | XP_012563475 | - | + | 2E-09 | 00063453 | - | + | 1E-15 | lachesin-like [Bemisia tabaci] | XP_018915454 | 0.11 |
| Alr11 | XP_012563475 | - | + | 1E-18 | 00052857 | - | + | 9E-16 | Lachesin like protein [Argiope bruennichi] | KAF8793681 | 0.044 |
| Alr12A | XP_012564659 | - | + | 3E-12 | 00052857 | - | + | 7E-15 | | | |
| Alr12B | XP_012563475 | - | + | 5E-09 | 00052857 | - | + | 7E-12 | unnamed protein product [Arctia plantaginis] | CAB3245167 | 2.4 |
| Alr15 | XP_012563475 | - | + | 4E-12 | 00052857 | - | + | 2E-14 | | | |
| Alr16 | XP_012563475 | - | + | 3E-05 | 00011113 | - | + | 5E-13 | kalirin isoform X7 [Xenopus laevis] | XP_041434124 | 1.9 |
| Alr17 | XP_012557852 | - | + | 3E-17 | 00063453 | - | + | 1E-14 | | | |
| Alr18 | XP_012564224 | - | + | 0.79 | 00052857 | - | + | 0.73 | hypothetical protein [Capitella teleta] | ELT96018 | 0.025 |
| Alr19 | XP_012557852 | - | + | 5E-08 | 00052857 | - | + | 3E-11 | protein amalgam-like [Eurytemora affinis] | XP_023337364 | 0.089 |
| Alr21 | XP_012557852 | - | + | 1E-14 | 00052857 | - | + | 2E-16 | titin [Strongylocentrotus purpuratus] | XP_030830392 | 0.063 |
| Alr23 | XP_012557852 | - | + | 4E-15 | 00064849 | - | + | 1E-13 | titin [Epinephelus lanceolatus] | XP_033471474 | 0.003 |
| Alr27 | XP_012564659 | - | + | 5 E-07 | 00064849 | - | + | 5E-09 | | | |
| Alr28 | XP_012563475 | - | + | 2E-05 | 00011113 | - | + | 7E-09 | Obscurin [Araneus ventricosus] | GBM92276 | 0.002 |
| Alr29 | XP_012563475 | - | + | 2E-09 | 00052857 | - | + | 1E-14 | CAM2-like [Biomphalaria glabrata] | XP_013096776 | 0.061 |
| Alr30 | XP_012563475 | - | + | 0.003 | 00005207 | - | + | 7E-06 | Lachesin [Sarcoptes scabiei] | KAF7491539 | 9E-6 |
| Alr31 | XP_012564659 | - | + | 6E-06 | 00064849 | - | + | 9E-11 | myosin light chain kinase, smooth muscle-like [Mizuhopecten yessoensis] | XP_021370306 | 0.36 |
| Alr33 | XP_012563475 | - | + | 1E-06 | 00011113 | - | + | 7E-12 | unnamed protein [Brugia pahangi] | VDN84797 | 0.34 |
| Alr34 | XP_012563475 | - | + | 8E-05 | 00052857 | - | + | 2E-11 | unnamed protein [Coregonus sp. 'balchen'] | CAB1323731 | 3E-10 |
| Alr35 | XP_012563475 | + | + | 7E-09 | 00052857 | + | + | 1E-21 | hypothetical protein [Rhynchophorus ferrugineus] | KAF7270589 | 4E-4 |
| Alr36 | XP_012563475 | + | + | 1E-17 | 00052857 | + | + | 2E-18 | neurotrimin [Drosophila subpulchrella] | XP_037726156 | 0.006 |
| Alr37 | XP_012557852 | + | + | 5E-10 | 00052857 | + | + | 1E-11 | titin-like [Cyprinodon tularosa] | XP_038159991 | 0.095 |
| Alr38 | XP_012563475 | + | + | 3E-08 | 00011113 | + | + | 1E-06 | | | |

[a] proteins encoded by *bona fide* genes in blue, putative genes in orange
[b] Signal Peptide predicted by SignalP
[c] Transmembrane helix predicted by TMHMM
[d] E-value cutoff = 10

**Dataset S1.** **FASTA formatted sequence of the ARC-F reference. Two gaps of unknown physical size are denoted with N's.**

**Dataset S2.** **GFF3-formatted annotations of *Alr* genes in the ARC-F reference sequence.**

**Dataset S3.** **FASTA formatted sequence of contig utg7180000000456, which contains *Alr37*.**

**Dataset S4.** **GFF3-formatted annotation of *Alr37* on contig utg7180000000456.**

**Dataset S5.** **FASTA-formatted sequence of contig utg7180000000115, which contains *Alr38*.**

**Dataset S6.** **GFF3-formatted annotation of *Alr38* on contig utg7180000000115**

**Dataset S7.** **FASTA-formatted cDNA sequences of bona fide genes in the *Alr* gene family.**

**Dataset S8.** **FASTA-formatted amino acid sequences of Alr proteins encoded by *bona fide* genes.**

**Dataset S9.** **FASTA-formatted cDNA sequences of putative genes in the *Alr* gene family.**

**Dataset S10.** **FASTA-formatted amino acid sequences of Alr proteins encoded by putative genes.**

**Dataset S11.** **FASTA-formatted MAFFT alignment of amino acid sequences for domain 1 from *bona fide* and putative *Alr* genes.**

**Dataset S12.** **FASTA-formatted MAFFT alignment of amino acid sequences for domains 2 and 3 from *bona fide* and putative *Alr* genes.**

**Dataset S13.** **FASTA-formatted MAFFT alignment of amino acid sequences for the ECS from *bona fide* and putative *Alr* genes.**

**Dataset S14.** **Zip file containing PDB files of the top predicted Colabfold models for all *bona fide* and putative domain 1, domains 2 and 3, and the trimmed ECS.**

**Dataset S15.** **FASTA-formatted amino acid sequences of the trimmed ECS used for structural predictions and alignment to fibronectin III domains.**
